# Genomic diversity generated by a transposable element burst in a rice recombinant inbred population

**DOI:** 10.1101/2020.08.24.265538

**Authors:** Jinfeng Chen, Lu Lu, Sofia M.C. Robb, Matthew Collin, Yutaka Okumoto, Jason E. Stajich, Susan R. Wessler

**Author notes:** J.C. and L.L. contributed equally to this work. To whom correspondence should be addressed. Susan R. Wessler, University of California Riverside, Department of Botany and Plant Sciences, Genomics 4107A, Riverside, CA 92521, Voice, (SRW).

## Abstract

Genomes of all characterized higher eukaryotes harbor examples of transposable element (TE) bursts - the rapid amplification of TE copies throughout a genome. Despite their prevalence, understanding how bursts diversify genomes requires the characterization of actively transposing TEs before insertion sites and structural rearrangements have been obscured by selection acting over evolutionary time. In this study rice recombinant inbred lines (RILs), generated by crossing a bursting accession and the reference Nipponbare accession were exploited to characterize the spread of the very active *Ping*/*mPing* family through a small population and the resulting impact on genome diversity. Comparative sequence analysis of 272 individuals led to the identification of over 14,000 new insertions of the *mPing* miniature inverted-repeat transposable element (MITE) with no evidence for silencing of the transposase-encoding *Ping* element. In addition to new insertions, *Ping*-encoded transposase was found to preferentially catalyze the excision of *mPing* loci tightly linked to a second *mPing* insertion. Similarly, structural variations, including deletion of rice exons or regulatory regions, were enriched for those with breakpoints at one or both ends of linked *mPing* elements. Taken together, these results indicate that structural variations are generated during a TE burst as transposase catalyzes both the high copy numbers needed to distribute linked elements throughout the genome and the DNA cuts at the TE ends known to dramatically increase the frequency of recombination.

**Significance Statement:** Transposable elements (TEs) represent the largest component of the genomes of higher eukaryotes. Among this component are some TEs that have attained very high copy numbers with hundreds, even thousands of elements. By documenting the spread of *mPing* elements throughout the genomes of a rice population we demonstrate that such bursts of amplification generate functionally relevant genomic variations upon which selection can act. Specifically, continued *mPing* amplification increases the number of tightly linked elements that, in turn, increases the frequency of structural variations that appear to be derived from aberrant transposition events. The significance of this finding is that it provides a TE-mediated mechanism that may generate much of the structural variation represented by pan-genomes in plants and other organisms.

## Introduction

Transposable elements (TEs) represent the largest component of the genomes of higher eukaryotes, comprising almost 50% of the human genome and over 80% of many plant genomes (1–4). Making up the TE component are hundreds, sometimes thousands of TE families, containing autonomous elements (encoding the enzymes that catalyze transposition) and the more numerous nonautonomous members (5). A subset of TE families attain very high copy numbers with hundreds, thousands, even tens of thousands of copies (5). Phylogenetic analysis of high copy number family members often reveals a star-phylogeny, indicative of a burst of amplification that is repressed (either by mutation or host-mediated silencing) and then the sequences of all copies drift into oblivion (5).

The prevalence of evidence for ancient bursts in plant and animal genomes implies that TE families have evolved mechanisms to insert copies throughout the genome while avoiding host silencing (6). To unravel these mechanisms requires the identification of TE families that are in the midst of a burst - where the biochemical features responsible for successful amplification are still active and their insertion sites have not been obscured by negative selection acting over evolutionary time.

To determine the features of a successful burst we characterized the *Ping*/*mPing* TE family, found to be amplifying in four accessions (EG4, HEG4, A119, A123) of *Oryza sativa* (rice) (7–9). The TE family is comprised of the autonomous *Ping*, a member of the *PIF/Harbinger* superfamily of Class II elements, and the miniature inverted-repeat transposable element (MITE), *mPing* (10). MITEs are a subset of nonautonomous DNA elements characterized by their short length (<600 bp) and ability to amplify from one or a few elements to hundreds or thousands of copies (11, 12). MITEs are the most abundant TE associated with the genes of higher plants where they populate noncoding regions (introns, 5’ and 3’ flanking sequences) and are a major contributor to allelic diversity (13, 14). While most characterized rice accessions have 0 to 1 *Ping* and 1-50 *mPings* (15), the four accessions have 7-10 *Pings*, ~230-500 *mPings*, and up to ~40 new *mPing* insertions per plant/generation (7–9).

Prior studies of the four inbred accessions where the *Ping*/*mPing* family has been active over decades revealed key features of successful bursts (9). First, although *mPing* has a preference for genic insertion sites, it can dramatically increase in copy number without having a major impact on the phenotype because of its preference for AT rich target sites while rice exons are GC rich (8, 16). Thus, *mPing* rarely inserts into rice exons (8). Second, because *mPing* is a deletion derivative of *Ping* but does not share any coding sequences, host recognition of *mPing* does not silence *Ping* expression (9). Despite robust host epigenetic regulation, the bursts have been ongoing for decades and are likely to continue until *Ping* transposes into a region that elicits host silencing or the TE load destabilizes the genome (9).

The focus of this study is another dimension of the *mPing* burst, that being impacts of its spread through a population. While TE bursts are rare phenomena, invasions of naïve populations by TE bursts in real time are exceedingly rare, with the most prominent example being the worldwide invasion of *Drosophila melanogaster*, and more recently, *Drosophila simulans* by *P* elements (17, 18). All available evidence indicates that bursts of both *P* elements and the *Ping/mPing* family occurred during the past century (9, 17, 18), however significant increases in *mPing* copy number is likely restricted to a few related accessions as no bursts were detected in over 3,000 sequenced rice accessions (15, 19). This likely reflects the fact that unlike *D. melanogaster*, which is a wild species and an outcrosser, rice is a domesticated crop that propagates by self- or sib-pollination.

In this study, a RIL population, previously constructed to assess the phenotypic consequences of *mPing* insertions (20), was repurposed to model the spread of the *Ping/mPing* burst. The parents of the population were the reference Nipponbare (where the *Ping/mPing* family rarely transposes) and HEG4, one of the 4 bursting accessions. F1s from this initial cross were used to establish 272 inbred lines following 10 generations of self- or sib-pollination (Fig. 1). As such, the RIL population serves as a proxy for the spread of a TE burst in a largely selfing species. Of note was that burst activity was maintained throughout the population as there was no evidence of silencing. In addition, comparative analysis of RIL genome sequences provided the first high-resolution picture of the extent of diversity generated by a TE burst including *mPing* insertions, excisions and structural rearrangements.

**Figure 1.**
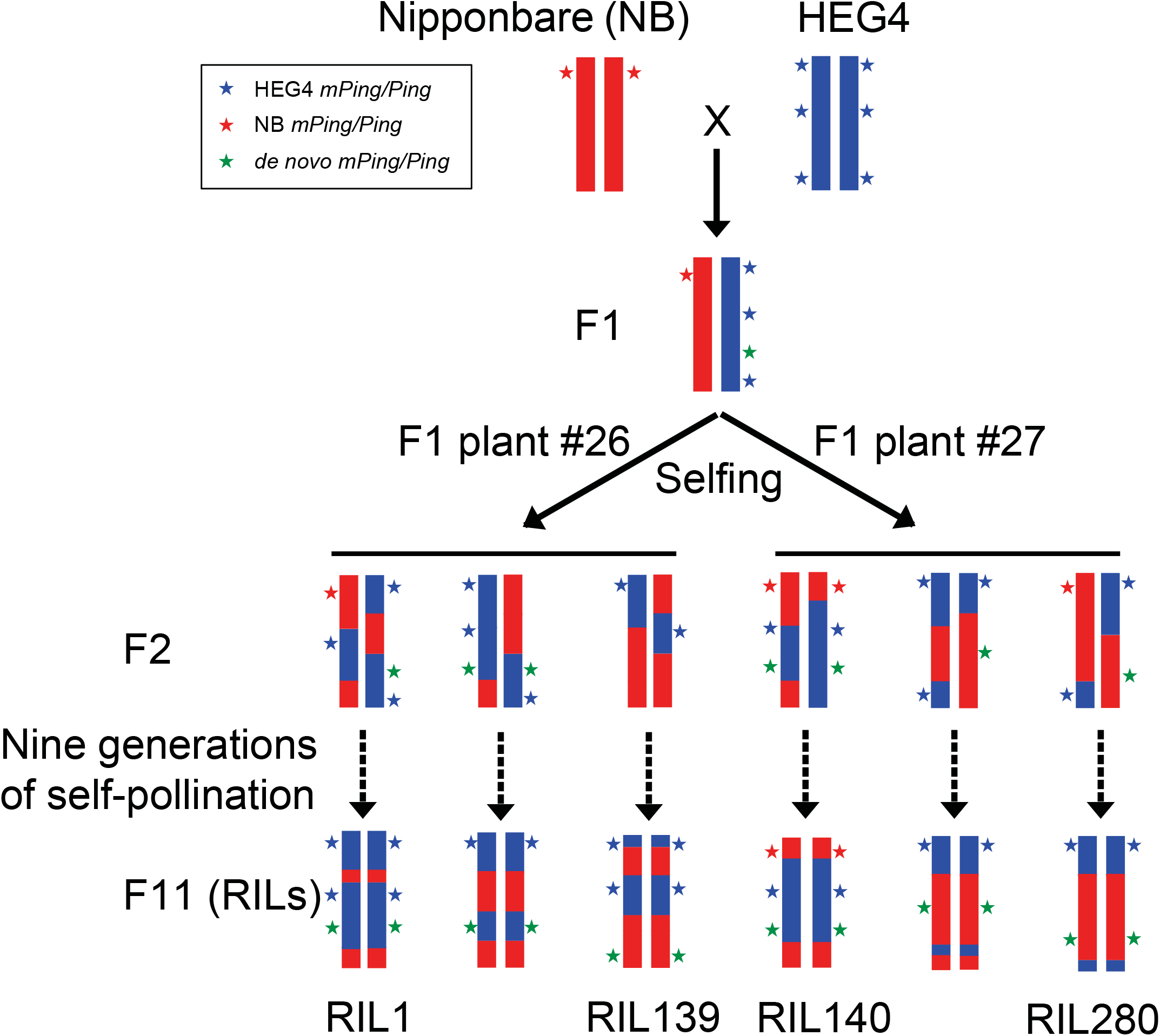
Schematic diagram of RIL construction. The RIL population was constructed by crossing Nipponbare (maternal) with HEG4 (paternal). Two F1 plants (#26 and #27) were used to breed F2s via self-pollination. F2 progeny were self-crossed for 9 generations to develop the RILs. HEG4 contains 7 *Pings* and 422 *mPings*, whereas Nipponbare contains 1 *Ping* and 51 *mPings*. After 10 generations of self-pollination, *Ping* and *mPing* elements from HEG4 (blue stars) and Nipponbare (red stars) segregated in the RILs and new *mPing/Ping* transpositions (green stars) are not in the RIL parents.

## Results

### Sequence and analysis of recombination map in the RIL population

A collection of 272 RILs derived from a cross between accessions HEG4 and Nipponbare (see Methods) was sequenced using Illumina paired-end reads. A total of ~1 Tb of sequences were generated with an average depth of ~11 -fold coverage per RIL (Supplementary Figure 1A and Supplementary Data 1). Sequence reads from each RIL were aligned to the Nipponbare reference genome (MSUv7) (21) and the genotype scored for the 105,900 high-quality SNPs that distinguish the parents (9). This approach determined 93.5% of parental SNP genotypes in every RIL, ranging from 60.9% to 99.7% (Supplementary Figure 1B and Supplementary Data 1). Recombination bins were inferred using a Hidden Markov Model (HMM) approach (22). The resulting recombination map contained 2,572 bins and an average bin length of 142.76 kb (Supplementary Figure 1C and Supplementary Table 1), which is comparable to other sequenced RIL populations in rice (23, 24) and enabled efficient quantitative trait loci (QTL) dissection of genetic loci controlling *mPing* transposition.

### *de novo mPing* and *Ping* insertion sites

The genomic locations of *mPing* elements in each RIL were determined with RelocaTE2 (25). A total of 87,450 *mPing* insertions were identified across the 272 RILs and simplified to 16,914 unique *mPing* loci in the population (Table 1 and Supplementary Data 2). Of these 16,914 loci, 3% are parental while the remaining 16,448, or 97% of *mPing* loci in the population are nonparental and defined herein as *de novo* insertions. Among the *de novo* insertions, 88% (14,534/16,448) are unique to a single individual indicating that most new insertions occurred during or after the F2 generations that produced the recombinant inbreds (Table 1). The rest of the *de novo* insertions (1,914/16,448) are shared among 2 to 145 RILs representing either early insertions that occurred in F1 or late insertions that share target sites in the genome (Supplementary Figure 2). Of those found in a single individual, 72% (10,527/14,534) are homozygous, and likely transposed in early generations of RIL self-pollination while the remaining 28% (4,007/14,534) are heterozygous (Table 1) and represent either more recent *mPing* insertions or *mPing* alleles that are lethal or less fit when homozygous. To distinguish these possibilities SNPs flanking heterozygous *mPing* loci were analyzed to determine genotypes of genomic regions underlying *mPing* insertions as recent insertions are more likely to be in homozygous (inbred) regions. Of the 4,007 heterozygous insertions, we were able to unambiguously genotype 2,964, of which 96% (2,864/2,964) were homozygous (Supplementary Table 2). These data suggest that the vast majority of heterozygous *mPing* insertions are late events. Finally, like previously reported *de novo mPing* insertions (7, 8), 45% of all insertions (both homozygous and heterozygous) identified in this study are within 5kb upstream of protein-coding genes (Supplementary Figure 3).

**Table 1.** Classification of parental and *de novo mPing, Ping*, and *Pong* loci in RILs

| Classification | Number of<br>*mPing* loci | Number of<br>*Ping* loci | Number of<br>*Pong* loci |
| --- | --- | --- | --- |
| Parental | 466 | 8 | 6 |
| -shared | 7 | 0 | 5 |
| -unique HEG4 | 415 | 7 | 0 |
| -unique NB | 44 | 1 | 1 |
| *de novo* | 16,448 | 17 | 0 |
| -shared | 1,914 | 0 | 0 |
| -unique homozygous | 10,527 | 17 | 0 |
| -unique heterozygous | 4,007 | 0 | 0 |

Analysis of *Ping* elements identified only 17 nonparental (*de novo*) *Pings* and over half of these (8 of 17) are within 5kb of protein-coding genes (Table1 and Supplementary Table 3). Among the population, RIL270 has two *de novo Ping* insertions (Supplementary Data 2), while each of 15 RILs had a single new *Ping* insertion. *de novo Ping* insertions were only found in RILs with two or more parental *Ping* loci (Supplementary Data 2).

### Correlation between *Ping* copy number and number of new *mPing* insertions

Among the 272 RILs, significant variation (0 to 138) was found in the number of unique *mPing* insertions (from here on restricted to *de novo* unique homozygous *mPing* insertions) (Supplementary Data 2), suggesting that multiple genetic loci control *mPing* transposition activity. To identify these loci, QTL mapping was performed using the number of unique *mPing* insertions as the trait (Supplementary Figure 4). Three major loci were identified with the logarithm of the odds (LOD) scores greater than 3.31 and accounting for 49% of the phenotypic variation (Supplementary Figure 5 and Supplementary Table 4). Two of these loci contain multiple *Ping* elements (*PingA, PingB* on chromosome 1 and *PingE, PingF, PingG* on chromosome 9)_confirming prior data that all bursting accessions had multiple *Pings* (7-10 *Pings*) (9). A third QTL located on chromosome 4 (accounting for 11.82% of phenotypic variance) does not contain a *Ping* element (Supplementary Figure 5 and Supplementary Table 4), suggesting that additional factors may contribute to the rate of transposition. These potential host factors are beyond the scope of this study and will not be further discussed.

To date, quantification of the impact of *Ping* loci on *mPing* transposition has been restricted to accessions with either the single *Ping* locus in Nipponbare *(PingH)* or the collective impact of the 7-10 *Ping* loci in the bursting accessions (HEG4, EG4, A123, A119) (9). In Nipponbare, *mPing* rarely transposes and *Ping* transcript levels are very low (9). In contrast, there is significant amplification of *mPing* and transcription of *Ping* in the accessions with 7-10 *Pings*, including the RIL parent, HEG4 (9). This population allows for higher resolution analyses of *Ping* copy number and transposition and transcription activity because the 8 parental *Ping*s are segregating, and, theoretically, should contain most combinations of *Ping* loci (Supplementary Figure 6). In fact, several RILs were found to contain from 0 to 8 *Ping* loci, and a positive correlation was found to exist between *Ping* copy number and the number of new *mPing* insertions (Fig. 2A, two-tailed Pearson’s correlation test, *r* = 0.71, *P* = 1e-42). For example, there are ~5 unique *mPing* insertions in RILs with one *Ping*, whereas RIL with 7 *Ping* elements have, on average, 65 unique *mPing* insertions. In addition, transcript levels of both *ORF1* (blue) and *TPase* (red) increased linearly with *Ping* copy number (Fig. 2B, two-tailed Pearson’s correlation test, *ORF1*: *r* = 0.85, *P* = 6e-14; *TPase*: *r* = 0.9, *P* = 4e-11), suggesting that a simple dosage relationship exists between *Ping* expression and transposition activity.

**Figure 2.**
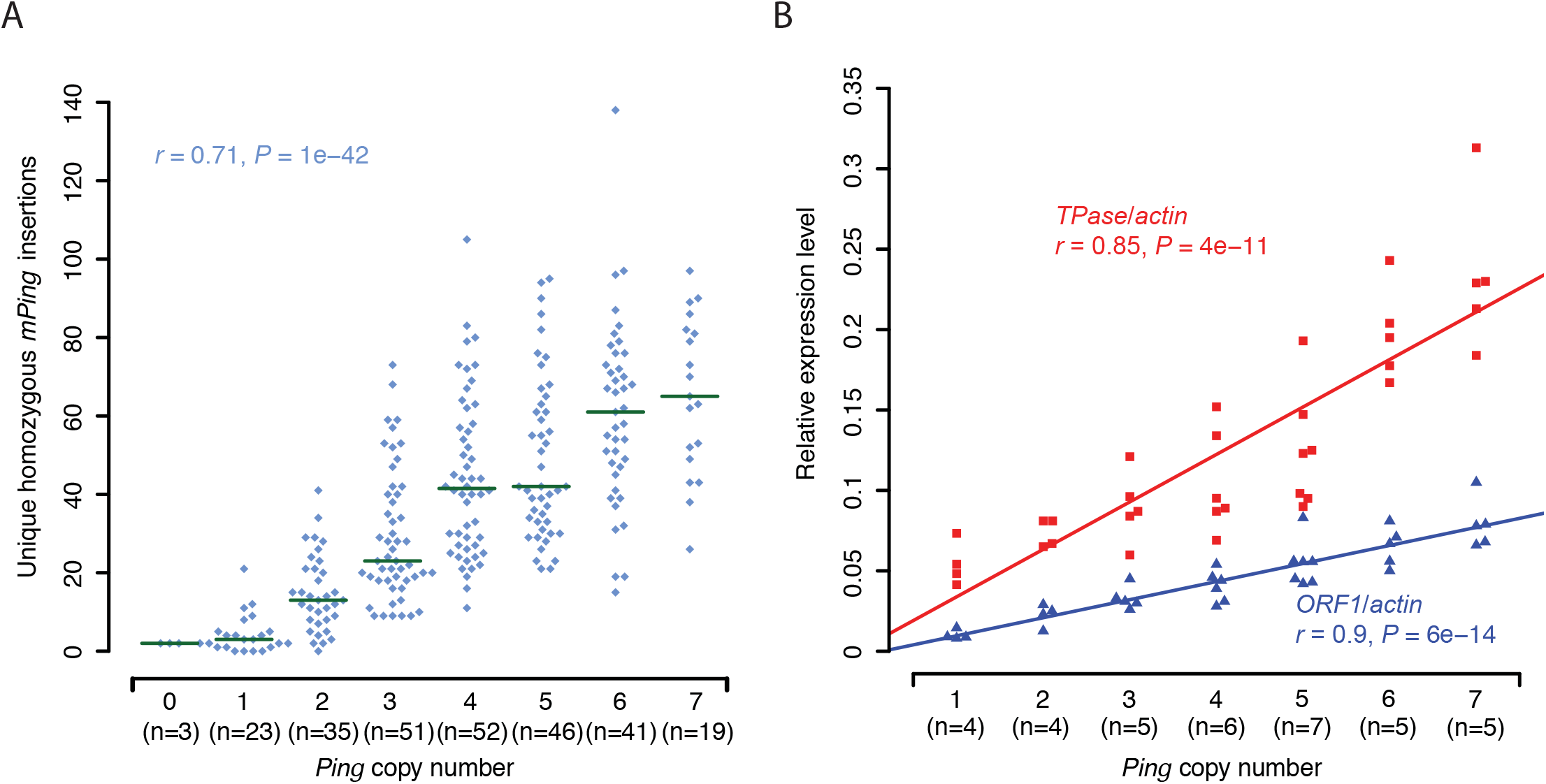
Accumulation of unique *mPing* insertions is dependent on *Ping* dosage. **A**, Correlation between unique *mPing* insertions and *Ping* copy number in the RILs. The 272 RILs were grouped by *Ping* copy number ranging from 0 to 7 and the scatterplot shows the number of unique *mPing* insertions in each group. The number of RILs in each group is in parentheses (n=). Green lines are the group median. The significance of correlation was tested by a two-tailed Pearson’s correlation test and indicated by *P* value. **B.** qRT-PCR analysis (3 replicates) of *Ping ORF1* and *TPase* transcription levels. RILs with *Ping* copy numbers of 1 to 7 were randomly selected. Relative transcription levels of *ORF1* (blue) and *TPases* (red) were normalized with rice *actin*. Colored lines are the best-fit line of the linear regression model. The significance of correlation was tested by a two-tailed Pearson’s correlation test and indicated by *P* values for both *ORF1* (blue) and *TPase* (red). Sampled RILs with 1 to 7 *Ping* copies used in Fig. 2B were: 1, (RIL12, 100, 166, 179); 2, (RIL16, 19, 36, 123); 3, (RIL5, 18, 22, 37, 111); 4, (RIL8, 13, 87, 119, 177, 219); 5, (RIL10, 15, 23, 58, 73, 92, 94); 6, (RIL30, 44, 54, 118, 134); 7, (RIL11, 21, 34, 69, 198).

### *Ping* activity at 8 different loci

The RIL population provided a unique opportunity to explore the activity of various combinations of the 8 segregating *Ping* loci present in the two parents. This was of particular interest because, although all *Ping* elements are identical, one *Ping* locus has been implicated in initiating the bursts (9, 15). Specifically, *PingA_Stow* on chromosome 1 (called *PingA* in this study) was shown previously to be the only *Ping* shared by the four bursting accessions (9) and found preferentially in the genomes of rice accessions with higher than background *mPing* copies (15). Taken together, these correlative data suggest that the *PingA* locus differs from the other *Ping* loci, perhaps by catalyzing more transposition.

The activity of individual *Ping* loci was quantified by first identifying RIL with one *Ping*. Among the RIL population are twenty-three lines with a single *Ping* that, collectively, harbor all parental *Pings* except *PingF. PingA* is present as a single *Ping* locus in RIL60. Because *mPing* rarely transposes in accessions with one *Ping* (like Nipponbare), two high-resolution independent methodologies, transposon display and deep sequencing, were employed to assess the number of new *mPing* insertions.

To identify new *mPing* insertions in individual progeny, Transposon Display (TD) was performed using DNA isolated from eight plants derived by self-pollination from each analyzed single-*Ping* RIL (Fig. 3A, Supplementary Figure 7 and Supplementary Table 5). The *mPing* transposition activity was estimated by counting new amplicons present in one individual but absent from its siblings. New *mPing* insertions were detected in all single *Ping* RIL analyzed except RIL179 (*PingB*) (Fig. 3A), however the progeny of RIL60 (with *PingA*), had significantly more (4-8 vs. 0-3 in the other single-*Ping* RIL) (Fig. 3A, one-way ANOVA with Tukey’s HSD test, *P* < 9.48e-8).

**Figure 3.**
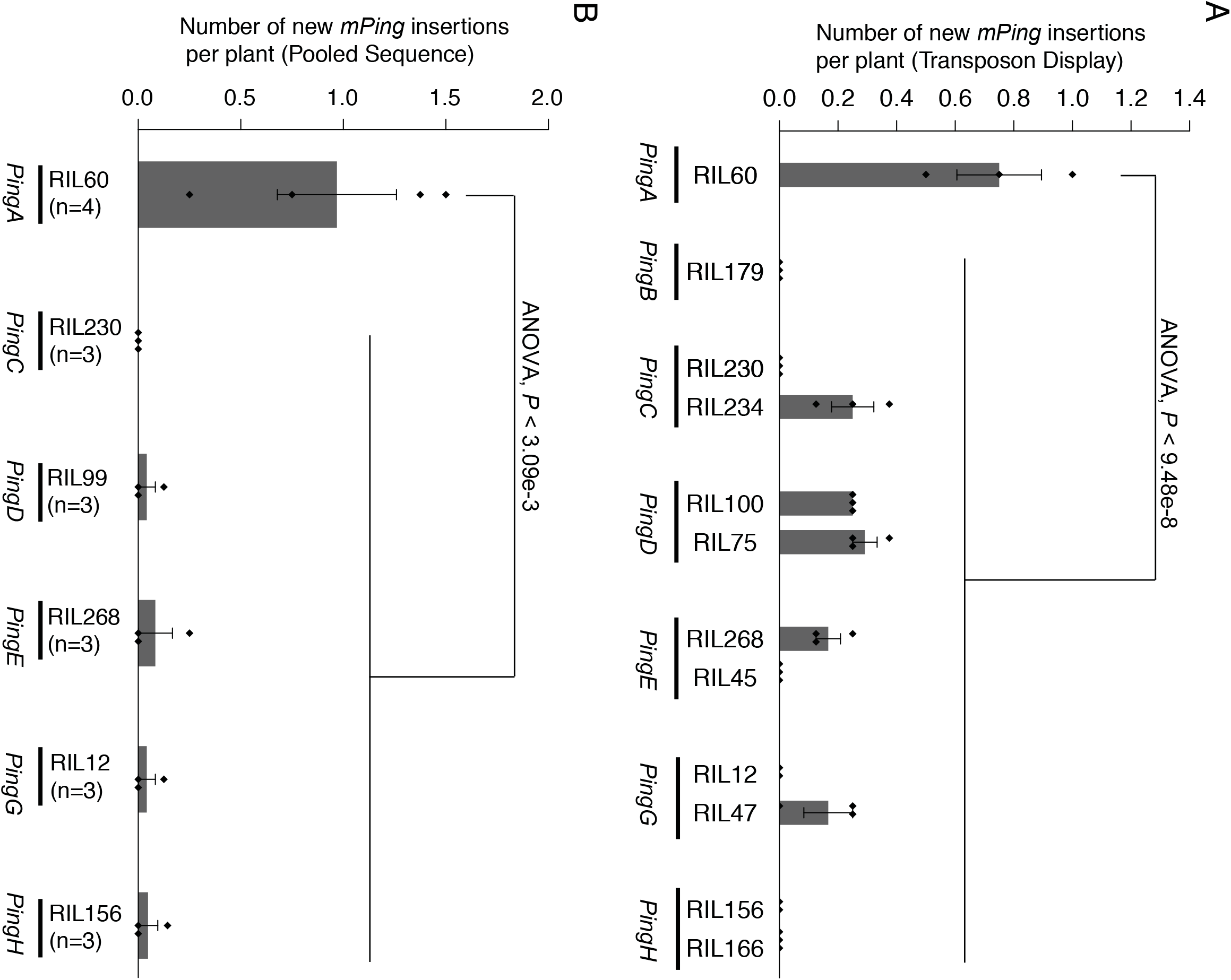
New *mPing* insertions in single-*Ping* RILs. **A**, New *mPing* insertions estimated by transposon display. New insertions were estimated by counting the number of new bands displayed by single plants. Eight sibling plants from a single-seed decent were used for each selected single-*Ping* RIL. Error bars show standard error of 2-3 independent biological replicates. **B**, New *mPing* insertions estimated by a pooled-sequencing approach with multiple progeny from a single-*Ping* RIL. RIL179 *(PingB)* was excluded due to contamination. Error bars show standard error of 3-4 independent biological replicates. Differences between *PingA* and other *Ping* loci were tested by a one-way ANOVA followed by Tukey’s HSD post hoc test.

In the second approach, new insertions were directly counted following deep sequencing of DNA samples isolated from 8-pooled progeny of each RIL. Detection of new *mPing* insertions for these RIL confirmed that all *Pings* tested are active thus providing a simple explanation for the *Ping* dosage series reported above. Similarly, the finding that *PingA* promotes more transposition than the other *Pings* (Fig. 3B and Supplementary Table 6, one-way ANOVA with Tukey’s HSD test, *P* < 3.09e-3) is consistent with the hypothesis that it initiated the bursts, but the underlying mechanism is beyond the scope of this study (see Discussion).

### Loss of parental *mPing* insertion loci in the RIL population

*Ping* and *mPing* elements, like other DNA transposons, can both insert into new loci and excise from existing sites. By tracking the fate of the 466 parental loci in the RIL population (44 from NB, 415 from HEG4, and 7 shared) (see Methods), we sought to determine whether some *mPing* loci were lost at a higher frequency than others. Loss of TEs over time has been attributed to many factors, the two most prominent being negative selection and structural instability (26, 27). To discriminate between these possibilities, we developed methodologies to identify all parental *mPing* excision events among the RIL population, determine excision frequencies at each locus, and analyze the genome context of loci that excised frequently. To this end we characterized empty (excised) *mPing* sites in RILs within haplotype blocks classified by parent of origin and expected to contain parental *mPing* loci (Fig. 4A). On average, 127 RILs (ranging from 71 to 159) were analyzed for each parental *mPing*.

**Figure 4.**
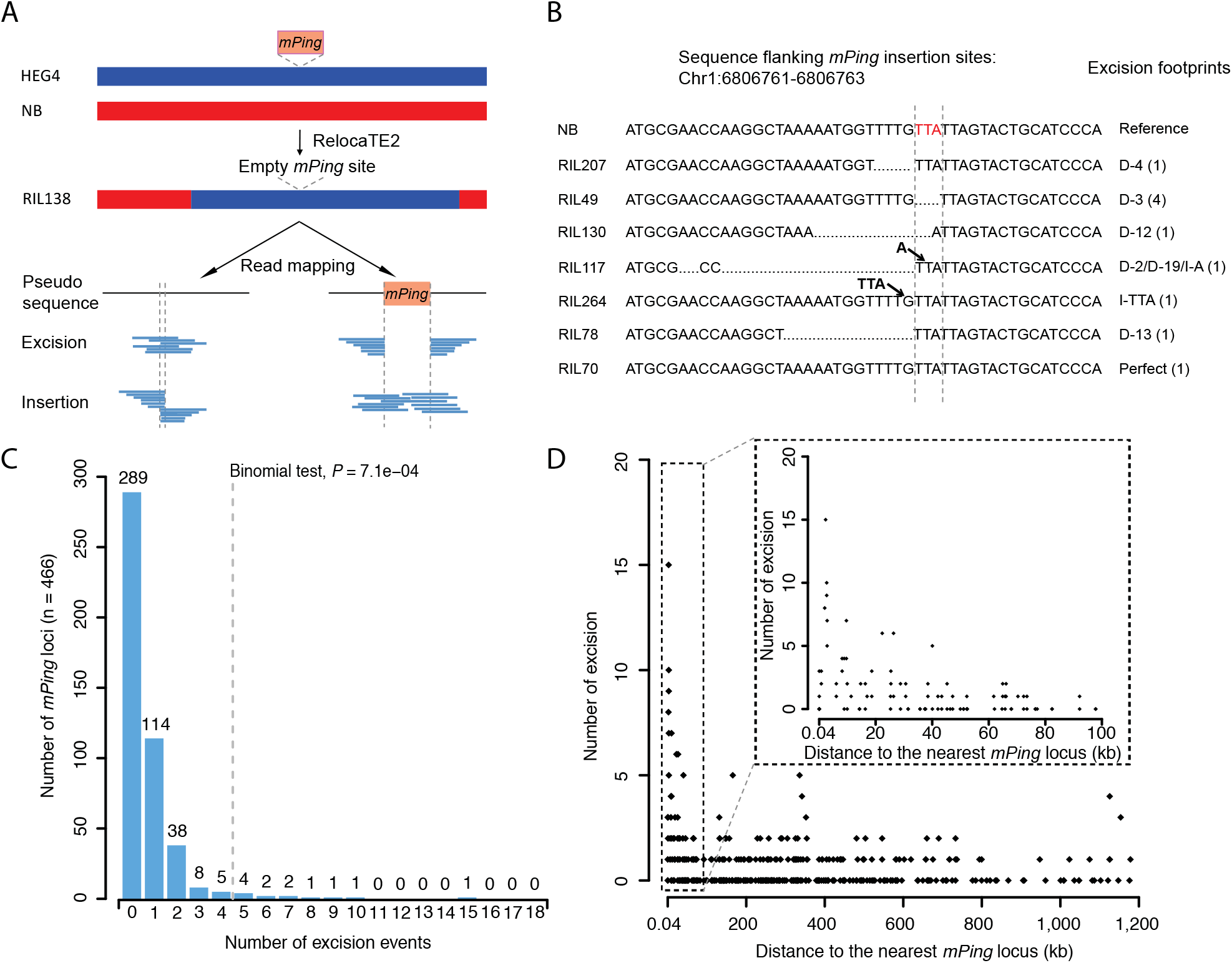
Identification of *mPing* loci with high excision frequencies. **A**, Workflow for identification of *mPing* excisions (See details in Methods). Red and blue colors represent Nipponbare (NB) and HEG4 genotypes in parental accessions. *mPing* insertions (orange boxes) are exclusively in HEG4 genotypes. Sequencing reads aligned to pseudoreference sequences with or without the *mPing* insertion are indicated by light blue bars. **B**, Example illustrating method used to detect independent excision events at *mPing* locus Chr1:6816415-6816417. Red letters in NB indicate the 3bp target site duplications (TSDs) upon *mPing* insertion. Footprints (deletions and insertions) are indicated by dots or arrows. CIGAR-like strings are used to record the footprints. Excision footprints are characterized as deletions (D) or insertions (A) with the length (e.g. D-X bp) and, in parenthesis, the number of RILs with this footprint. **C**, Distribution of parental *mPing* loci and number of independent excision events. The *P* value is based on a two-tailed binomial test. Dashed line is the cutoff for significant high frequency excision. The number of different *mPing* loci with that many excisions is at top of bar. **D**, *mPing* loci with more excision events are in close proximity to another *mPing*. Box shows a zoom of the 0.04 to 100 kb range.

In total, 742 excisions were detected from 177 parental *mPing* loci (17 NB, 160 HEG4), and no excisions were detected from the remaining 289 loci (34 NB, 255 HEG4). To validate the accuracy of the computer-assisted excision data, 30 loci were randomly selected and assayed by PCR with DNA isolated from the RIL where the excision was detected. Using this approach, 26 of 30 events were confirmed (Supplementary Figure 8). Next, to assess the excision frequency at each of the 177 parental loci, it was necessary to establish the independence of the excision events. For example, an excision event occurring early in the generation of the RIL population might be propagated in several RILs. Two approaches were used to eliminate dependent events: (1) a standard method where all excisions were analyzed after removal of early events, and (2) a conservative method where excision footprints were exploited to identify independent events. The latter is likely to underestimate independent events because perfect excision, which is common for *mPing*, will only be counted once per locus per RIL. Because both approaches produced very similar results, only the conservative method is described below. However, the results of both are found in the Methods.

For the conservative method, 367 of the 742 excisions from parental *mPing* loci have excision footprints (Supplementary Data 3). This reduced to 322 independent events following removal of excisions from the same locus with the same footprint (Fig. 4B and Supplementary Data 3). Of the 177 loci, 114 had only a single excision, 51 had fewer than five (Fig. 4C) and 12 loci experienced five or more independent excision events (Fig. 4C and Supplementary Table 7), which is significantly more than expected by chance (two-tailed Binomial test, *P* = 7.1e-4). Significantly, a majority of these loci (7 of 12) have another *mPing* insertion within 10kb. Of these 7 loci, 5 have a nearby *mPing* in HEG4, including two pairs of high frequency excision loci located close to each other (2.8 kb between Chr1:36267659 and Chr1:36270511; 2.6 kb between Chr5:15210313 and Chr5:15213006) and one (Chr1:6806761) located 9.6 kb from another parental *mPing* (Chr1:6816415) with 4 independent excisions (Fig. 4D and Supplementary Table 7). Two of the 7 loci have a nearby *mPing* (1.9 kb and 2.3 kb away) that is not a parental locus but rather present in 45% of the RILs (Supplementary Table 7). For the remaining 5 of 12 high excision *mPing* loci, the nearest *mPing* is from 22.3 to 335.8 kb (Supplementary Table 7). Taken together, these findings indicate that the loss of *mPing* among the RILs can be attributable to structural instability, not negative selection. In support of this claim is the finding that expression analysis of the protein-coding genes associated with the 12 high frequency excision loci identified only a single differentially expressed gene between the two parental accessions (LOC_Os01g07300) (Supplementary Table 8).

### *mPing* mediated sequence rearrangements

Another measure of the impact of high copy *mPings* on genome stability is the number of structural rearrangements in the RIL population and their proximity to *mPing* insertions. A prior study, limited to a single comparison between two accessions with high *mPing* copy numbers (HEG4 and EG4), identified a 120kb inversion with multiple *mPing* at the breakpoint (9).

The RIL population, with 97% having 200 or more *mPing* copies (Supplementary Data 2), provided an opportunity to investigate the impact, if any, of an active burst on the generation of structural rearrangements. A limitation of this analysis is that the use of short sequence reads precludes the identification of inversions mediated by *mPing* or other TEs; only deletions and small insertions (duplications) could be detected with confidence. Positions of rearrangements, irrespective of their proximity to *mPing* elements, were determined by aligning and comparing RIL sequences with the Nipponbare reference genome using a read depth strategy implemented in CNVnator (28). Rearrangements only in the RILs, were identified by first analyzing HEG4 using the same method to detect and eliminate from consideration rearrangements in the parental accession. In total, 16 rearrangements (15 deletions and 1 deletion plus duplication), ranging from ~700 bp to 56 kb, were resolved (Table 2 and Supplementary Table 9). Fourteen of the sixteen were only in a single RIL, while two are shared in several RILs and found to derive from a single F1 plant (#27, see Methods) that segregated in F2 generations (Supplementary Table 10). Of note, 12 of the 16 rearrangements have *mPings* in close proximity to the breakpoints with nine starting precisely at *mPing* sequences (Fig. 5, Supplementary Figures 9-10 and Table 2). Eleven *mPing* loci were identified at the breakpoints of these 9 rearrangements. Of these 11 loci, 5 were also characterized as high frequency excision loci (Supplementary Table 11). The 9 rearrangements with *mPing* sequences at the breakpoints were all validated by PCR (Supplementary Figure 11 and Table 2).

**Figure 5.**
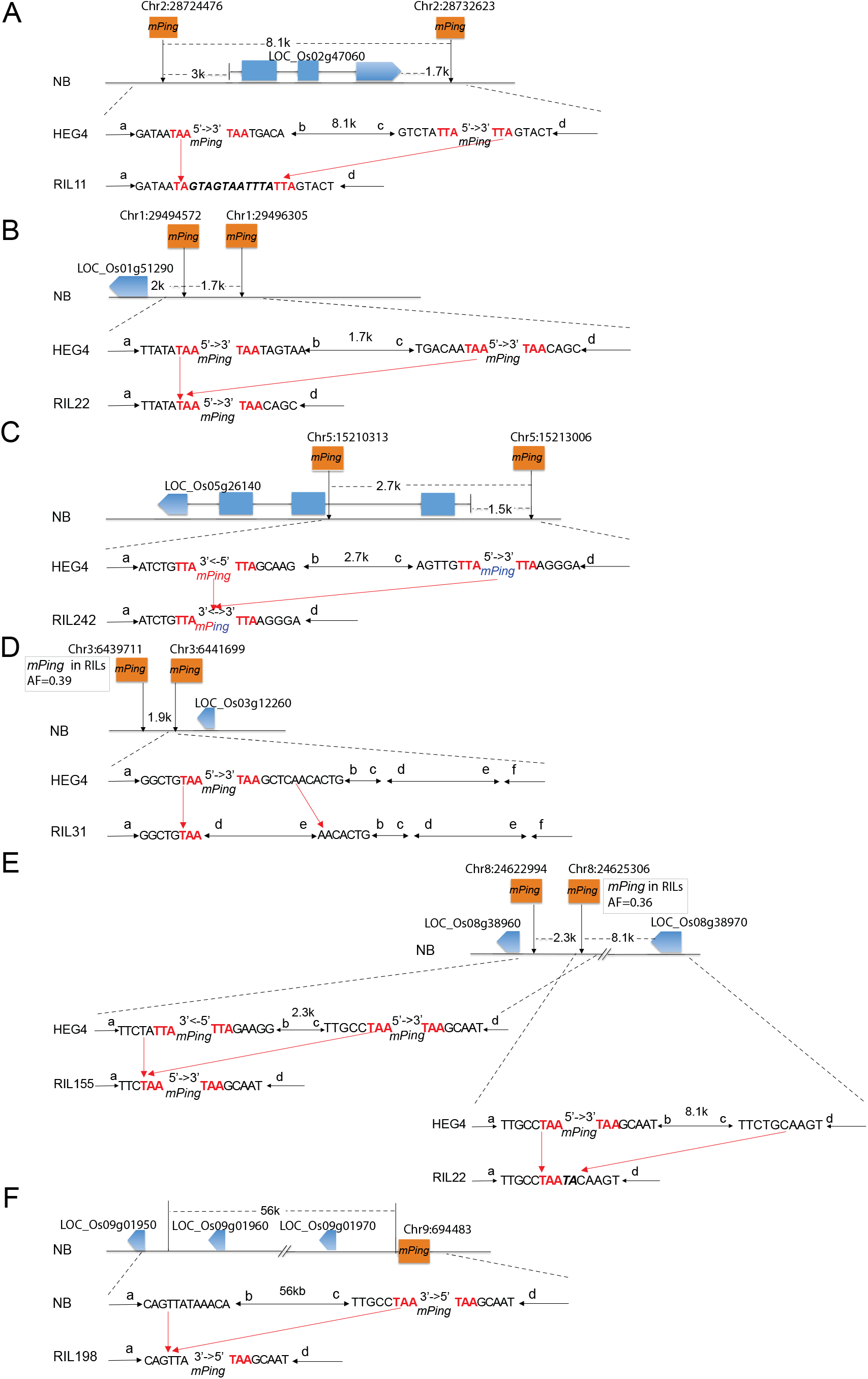
Structural variations mediated by *mPing* elements. **A**, SV2. **B**, SV1. **C**, SV6. **D**, SV4. **E**, SV7 and SV8. **F**, SV9. *mPing*-associated structural variations in select RILs were aligned with either Nipponbare (NB) or HEG4 based on the chromosome of origin. Lines indicate DNA fragments flanking *mPing* insertions or the breakpoint of structural variation. Letters (a-f) on lines label the DNA fragment end. Orange boxes indicate *mPing* insertions. Blue boxes indicate exons of protein-coding genes. Target site duplications (TSD) of *mPing* insertions are highlighted in red. Filler DNA at breakpoints is italicized. Red arrows indicate the breakpoints where structural variation occurred. Allele frequency (AF) is indicated if *mPing* is present in RILs but is absence in parental accessions.

**Table 2.**
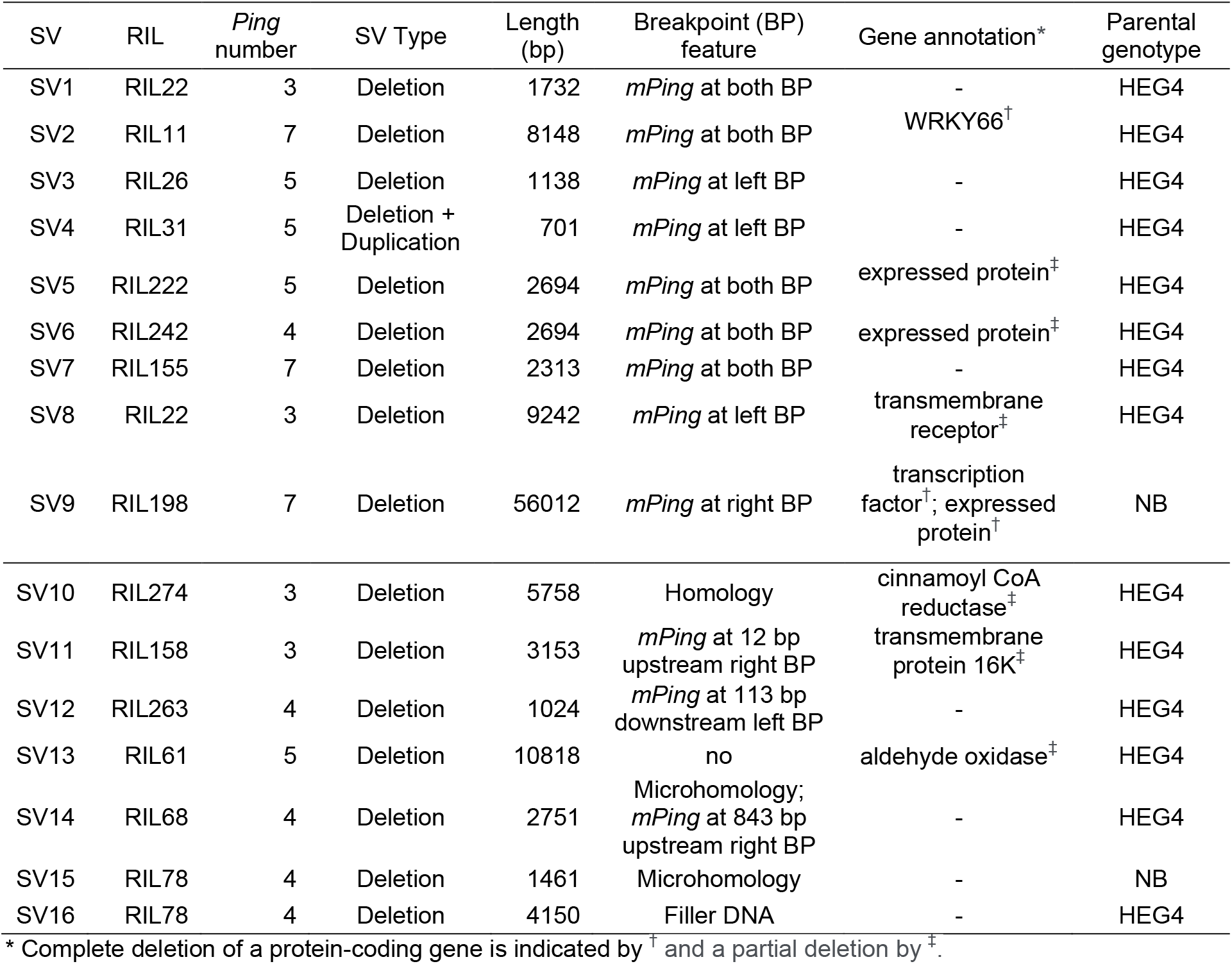
Features of structure variations in RILs

Close examination of these 9 rearrangements implicate aberrant transposition as the underlying mechanism. For example, transposase acting at the distal termini of *mPings* upstream and downstream of rice gene LOC_Os02g47060 (Fig. 5A and Supplementary Table 12), likely generated a megatransposon that includes both elements and 8.1 kb of intervening DNA. Similarly, termini of linked *mPings* appear to be involved in the deletion of potential regulatory sequences upstream of LOC_Os01g51290 (Fig. 5B and Supplementary Table 12) and intragenic sequences of LOC_Os05g26140 (Fig. 5C and Supplementary Table 12) leaving at the breakpoint, a single *mPing* in the former and a rearranged *mPing* in the latter. Deletion of sequences downstream of LOC_Os03g12260 (Fig. 5D and Supplementary Table 12) also correlates with the presence of tightly linked *mPings* although only one *mPing* terminus appears to be directly involved in the rearrangement. Tightly linked *mPings* are also involved in distinct intergenic rearrangements in RIL155 and RIL22, with the termini of both *mPings* implicated in the former, but only one of the two in the latter (Fig. 5E). The largest rearrangement with *mPing* at a breakpoint (a 56 kb deletion including 2 rice genes, Fig. 5F and Supplementary Table 12) involves only a single, unlinked *mPing* that is derived from the NB parent. Its participation in a rearrangement may be due to high transposase concentration – as RIL198 has 7 *Pings* (Table 2). In total, rearrangements were associated with deletion of four protein-coding genes and partial deletion of six protein-coding genes (Supplementary Table 12).

## Discussion

In this study recombinant inbred lines generated by crossing a bursting accession and the reference Nipponbare were exploited to characterize the spread of TEs through a small population and the resulting impact on genome diversity. To quantify family (transposase) activity, the number of new *mPing* insertions was used as a trait that varied across the RILs. In this way, the number of new *mPing* insertions per RIL served as a high-resolution proxy for *Ping* activity and led to a deeper understanding of the features of this TE family and its interaction with the host. To understand the impact on gene and genome diversity, all *Ping* and *mPing* insertions were localized and a determination made of their contribution to structural variation. Conclusions from these dual analyses are summarized below.

### Assessing *Ping* activity

#### Ping dosage

Prior studies of *Ping* activity were restricted to accessions, like NB, with a single *Ping* element, and the bursting accessions, HEG4, EG4, A119, and A123 with 7-10 *Pings* (9). Transposition of *mPing* was extremely rare in the former and robust in the latter with up to ~40 new insertions/plant/generation (7, 8). In this study, segregation of the 8 parental *Ping* loci generated lines with 0 to 8 *Pings*, often with many RILs for each *Ping* dosage (Supplementary Data 2). Variation in the number of new *mPing* insertions per RIL (0 to 138) led to the identification of two QTLs that each contain multiple linked *Ping* elements: on chromosomes 1 (*PingA*, *PingB*) and 9 (*PingE*, *PingF*, *PingG*) (Supplementary Figure 5). That these QTLs contain the only linked *Pings* raises the question of whether their contribution to *mPing* transposition is due to the effect of *Ping* dosage or something more subtle. We think the former for the following reasons. First, RILs with *PingC, PingD*, and *PingH* contain comparable numbers of new *mPing* insertions as RILs with *Ping* loci identified by QTL mapping (Supplementary Figure 6B-G). This suggests that single *Ping* loci also contribute to dosage effect on transposition activity. Second, two other bursting accessions (A119, A123) with high *Ping* copy numbers (7 and 10 respectively) only share the *PingA* locus (9). That is, their bursts are robust even though they do not have the 2 QTLs reported in this study. Thus, we conclude that variation in the number of new *mPing* insertions across the RILs reflects increasing *Ping* copy number (Fig. 2A) and *Ping* transcripts (Fig. 2B). These results indicate that incremental increases in *Ping* encoded products promote incremental increases in *Ping* activity. Just how high *Ping* dosage can be increased before activity plateaus is the goal of a future study to investigate new RIL populations where both parents are in the midst of a burst and many more *Pings* and *mPings* are segregating.

#### Individual Ping activity

Segregation of the seven (out of eight) *Ping* loci (containing identical *Ping* elements) into single-*Ping* containing RILs facilitated, for the first time, a comparative analysis of their ability to catalyze *mPing* transposition. To overcome the limitation of low germinal activity associated with single *Ping* containing lines, we exploited the higher frequency of somatic transposition coupled with deep sequencing. Consistent with prior studies where *PingA* was found to be the only locus shared by all bursting and high copy accessions (9), we found that *PingA* catalyzes more transposition than the other *Ping* loci (Fig. 3). Although these data could be interpreted to mean that the burst was initiated by higher *PingA* activity, future studies are needed to determine the underlying mechanism.

#### Ping is not silenced

The *Ping/mPing* family was shown previously to successfully amplify by eluding epigenetic silencing for decades of sib- or self-pollination of bursting inbred lines (9). In this study we extend that finding by showing that this TE family remains active after outcrossing (of a bursting accession to Nipponbare) and subsequent rounds of sib- or self-pollination of the RIL population. First, demonstration that there are no new insertions of *Pong*, which is active but silenced (9), in any RIL indicates that epigenetic regulation is maintained throughout the population (Table 1 and Supplementary Data 2). Despite robust epigenetic regulation, all RIL with *Ping* elements, especially lines with many *Pings*, have new *mPing* insertions indicating that *Ping* has not been silenced in any of these lines (Supplementary Data 2).

Data from this study provides important clues to why *Ping* activity has persisted in the population after outcrossing, and why, as the burst continues, it is increasingly unlikely to be silenced. A prior study demonstrated that although *mPing* is recognized by host epigenetic machinery, *Ping* is not silenced because *mPing* does not harbor any of *Ping*’s coding sequences. Thus, only *Ping* terminal sequences, that are shared with *mPing*, are methylated (9). These results suggested that for *Ping* to be silenced it has to transpose into a genomic region that generates siRNA. Here we show that *Ping* rarely transposes, making *Ping* silencing unlikely. Only 17 new *Ping* insertions were detected among the RIL population – compared to over 14,000 new *mPing* insertions (Supplementary Data 2). A close examination of the RILs with transposed *Pings* identified from 21 to 96 new *mPing* insertions including 4 to 60 heterozygous insertions that are likely to be new and provide evidence of recent activity (Supplementary Table 3). In addition, of the 17 new *Ping* insertion sites, 14 are into euchromatic loci and 3 into heterochromatin (Supplementary Table 3). Among this latter class (which is more likely to silence *Ping)*, new *mPing* insertions vary from 63 (21 are heterozygous) to 67 (15) to 111 (15), again inconsistent with the loss of *Ping* activity (Supplementary Data 2).

Taken together, these data suggest that as the burst continues, the ratio of *mPing* to *Ping* copies will increase – which is likely to further decrease the frequency of *Ping* transposition and its subsequent silencing. In addition, the finding that the single SNP adjacent to its terminal inverted repeats (TIRs) makes *mPing* a better substrate for transposase than *Ping* (15), will further reduce the chances of *Ping* silencing as the burst continues.

### Genome instability as the burst continues

If the burst is not likely to end with *Ping* silencing, how will it end? Data from this study strongly suggests that increases in *mPing* copy number destabilize the genome by elevating the chances of potentially deleterious structural variation (SV). TEs have long been known to generate SVs. The first TE discovered by McClintock, *Dissociation* (*Ds*), was initially detected as a site of chromosome breakage (29). These so-called “breaking” *Ds* elements, are a structurally distinct minority of characterized elements, often composed of tightly linked *Ds* copies (30, 31). Double-*Ds* elements form frequently because of the preference of the *Ac* transposase to catalyze local transposition (32, 33). Similar local hops and chromosome breakage are associated with TEs from all three kingdoms including the Drosophila *P* and the bacterial Tn*10* elements (34, 35).

In contrast, *Ping* does not promote local transposition of *mPing* as new insertions from a single integrated donor site in transgenic *Arabidopsis thaliana*, yeast, and soybean were not linked to that site (16, 36, 37). In operational terms, this means that *mPing* bursts scatter copies randomly throughout the genome with only a few near existing insertion sites. However, as the burst continues, the probability that new insertions will be linked to existing *mPing* elements increases. Whether linked *mPing* elements are more likely to generate SVs was investigated in the RIL population in two ways: (i) by quantifying the relative frequencies of parental *mPing* excisions, and (ii) by identifying SVs in the RIL population and determining whether any had *mPing* at one or both breakpoints. For the analysis of excisions, the RILs were scored for the frequency of excision among the 466 parental *mPing* loci. The fact that 7 of the 12 loci with 5 or more excisions had another *mPing* within 10kb suggests that tightly linked elements are unusually active with regard to transposition (Supplementary Table 7).

Whether higher transposition activity also contributes to the formation of a subset of structural variations was assessed by identifying deletions and small duplications among the RILs. As mentioned in the Results, a limitation of this analysis is that the use of short sequence reads precludes the identification of inversions and many larger SVs. Overall, sixteen SVs were detected, and 9 shown to have *mPing* at one (5/9) or both (4/9) breakpoints (Table 2). Furthermore, the positions of the breakpoints – at *mPing* termini – implicate *Ping* transposase activity in SV formation. This situation is reminiscent of the occurrence of *Ds* breakage only if *Ac* is in the genome (29), and that aberrant transposition is the likely mechanism (38, 39).

## Conclusions and implications

The results of this study suggest that there may be an evolutionary benefit to the host when a TE family bursts, scattering copies throughout the genome. Specifically, continued *mPing* amplification increases the number of tightly linked elements, which, in turn, increases the number of functionally relevant SVs upon which selection can act. Most of the SVs reported in this study are deletions that are unlikely to be of evolutionary benefit. However, this limitation probably reflects the use of short read sequences that have been reported to miss most of the SVs larger than 2kb including duplications and inversions (40). In fact, a prior study that compared the assembled genomes of two virtually identical rice accessions undergoing independent *mPing* bursts: HEG4 and EG4, detected an inversion of ~120kb with *mPings* at the breakpoints (9).

The significance of this finding is that it provides, for the first time, a TE-mediated mechanism that may generate much of the SVs represented by pan-genomes in plants and other organisms (41, 42). Specifically, it is generally accepted that a species, or even a population, cannot be adequately represented by a single reference genome (43). Rather, members of a species share core genes, but variation exists in the form of sequence rearrangements that alter so-called dispensable (or accessory) gene content. Associations of dispensable genomic regions with more TEs suggest that TEs are involved in generating observed presence, absence, and copy number variations (44). However, until this study, the only evidence was correlative; questions remained concerning when and how TEs generate the raw material for pan-genomes. Here we demonstrate that “when” is during an active TE burst, and “how” is through the scattering of tightly linked elements throughout the gene space, thereby seeding the genome with regions susceptible to transposase mediated SVs. One could imagine that in a sufficiently large population undergoing similar bursts, SVs that involve components of the core genome will be eliminated by negative selection while SVs that alter dispensable genes could be retained in the population.

## Materials and Methods

### RIL plants, DNA extractions and sequencing

Recombinant inbred lines (RILs) were developed through a single-seed descent approach from the F2 generation of a cross between Nipponbare (Maternal parent) and an *mPing* active accession HEG4 (Paternal parent) (See details in Fig. 1). RIL seeds were germinated at 37 °C and moved to soil 5 days after germination. Seedlings of 3-week-old plants were harvested to extract genomic DNA using the CTAB method (45). Purified genomic DNA was fragmented to 300 bp using a Covaris S220 Ultrasonicator (Covaris) and paired-end libraries were constructed using Illumina TruSeq DNA sample prep kit (Illumina) or KAPA LTP library preparation kit (Kapa Biosystems). Libraries were multiplexed and sequenced on HiSeq 2500 or NextSeq 500 (Illumina).

### RNA extraction and qRT-PCR

RILs were grown as described above. Seedlings of 3-week-old plants were used to extract total RNAs using the RNeasy Plant Mini Kit (Qiagen). Total RNAs were treated with amplification-grade RNase-free DNase I (Qiagen) to remove contaminating DNAs. The treated DNA-free total RNAs were reverse transcribed into cDNAs using SuperScript III first-strand supermix (Invitrogen). The resulting cDNAs were used as templates to perform qRT-PCR with iQ SYBR Green Supermix (Bio-Rad) on CFX96 system (Bio-Rad). Transcript levels of *ORF1* and *TPase* were normalized to rice *actin*. Primers for qRT-PCR are in Supplementary Data 4.

### Construction of recombination bin and linkage map

A total of 105,900 SNPs between parental accessions HEG4 and Nipponbare were identified and filtered using GATK (46) as described previously (9). High-quality SNPs were used to genotype the RILs and a Hidden Markov Model (HMM) algorithm was applied to construct recombination bins (22). Briefly, paired-end reads of each RIL were mapped to Nipponbare reference genome (MSUv7) by Burrows-Wheeler Aligner (BWA) aligner v0.6.2 (21, 47). Read alignments were sorted and PCR duplicates were removed using MarkDuplicate as implemented in PicardTools (http://broadinstitute.github.io/picard/). Potential SNP sites were identified by GATK toolkit following the best practice workflow of GATK (46). Only SNP sites matching parental SNPs were considered as informative sites. A genotyping error rate of 1.4% was estimated by counting the number of incorrectly genotyped SNPs from independently sequenced parental samples, Nipponbare and HEG4. Of these 105,900 SNPs, 1,311 were misgenotyped as HEG4 or heterozygous SNPs in the Nipponbare sample (864 HEG4 and 447 heterozygous, respectively), whereas 1,656 were misgenotyped as Nipponbare or heterozygous SNPs in the HEG4 sample (333 Nipponbare and 1,323 heterozygous, respectively). The resulting RIL SNP genotypes were imported into the R environment to construct a recombination bin map using the HMM approach as described in the Maximum Parsimony of Recombination (MPR) package (22). The expected proportion of genotypes for each locus was 49.75:0.50:49.75 with an error rate of 1.4%. Recombinant bins were used as markers and the missing data was imputed using an argmax method in the R/qtl package (48). A linkage map was built using the haldane function in the R/qtl package (48).

### Identification of *mPing* and *Ping* copies in the RILs

*mPing* copies, including elements present in the Nipponbare reference genome and elements present in the RILs or HEG4, were identified by RelocaTE2 (25). Briefly, RelocaTE2 identified paired-end read pairs that match the ends of *mPing* (junction reads) or internal sequences of *mPing* (supporting reads) using BLAT (49). After trimming *mPing* sequences, the remaining sequences from the original read pairs were searched against MSUv7 by BWA v0.6.2 to identify potential *mPing* insertions either present in the reference genome or in the RILs. Raw results from RelocaTE2 were filtered by removing *mPing* insertions with junction reads from only one end or *mPing* insertions without 3 bp target site duplications (TSDs), which is hallmark of *mPing* transposition. Heterozygous *mPing* insertions were characterized by analyzing BAM files of raw read pairs aligned to MSUv7 (50). These *mPing* insertions with reads aligned to both sides of the flanking sequences were characterized as heterozygous *mPing* insertions. *Ping* copy number in RILs was characterized as described for *mPing*. Read pairs from each *mPing/Ping* insertion site were analyzed to distinguish *mPing* and *Ping* sequence differences. All candidates *Ping* insertions were confirmed by PCR or Southern blot.

### Determination of *mPing* insertion sites in single-*Ping* RILs

Two experimental approaches (Transposon Display and Pooled DNA sequencing) were performed on multiple individuals to determine *de novo mPing* insertions. Because *mPing* transpositions are rare in single-*Ping* RILs, multiple individuals were pooled to maximize detection of new insertions catalyzed by each *Ping*.

Transposon Display: single-*Ping* RILs used in the experiment were *PingA:* RIL60, *PingB:* RIL179, *PingC:* RIL230, RIL234, RIL43, *PingD:* RIL100, RIL25, RIL75, *PingE:* RIL268, RIL45, *PingG:* RIL12, RIL47, *PingH:* RIL156, RIL166, RIL259. Genomic DNA was extracted using the CTAB method (45). Transposon display of *mPing* was performed on eight progeny of each RIL as described (51). Briefly, 50 ng genomic DNA was digested with *Mse*I and ligated with adapters overnight at 37°C. Preselective amplifications were performed with primers complementary to the adapters (*Mse*l + 0) and internal *mPing* sequences (*mPing* P1). Selective amplifications were performed with one selective base of the adapter primer (*Mse*I + T/A/G/C) and *mPing* P2 primers. PCR reactions were loaded on 6% denaturing acrylamide-bisacrylamide gels to visualize *mPing* transpositions. New bands that are present exclusively in one individual are counted as *de novo mPing* insertions. Transposition frequency of *mPing* was calculated using the number of new *mPing* insertions per individual from all the selective primer combinations. The primers for transposon display are in Supplementary Data 4.

Pooled DNA sequencing: single-*Ping* RILs used in the experiment were *PingA*: RIL60, *PingC*: RIL230, *PingD*: RIL99, *PingE*: RIL268, *PingG*: RIL12, *PingH*: RIL156. Genomic DNA from 7-9 plants per RIL were extracted, fragmented as described above and paired-end libraries constructed with KAPA LTP library preparation kit. Libraries were multiplexed and sequenced on HiSeq 2500 or NextSeq 500. Three or four biological replicates were performed for each experiment. RelocaTE2 was employed to identify *mPing* insertion sites in each sequence replicate. Only heterozygous *mPing* insertions that are exclusively present in one replicate were counted as *de novo* insertions.

### Analysis of *mPing* excision events

Parental *mPing* insertions segregating in the RILs were identified. First, 51 *mPing* insertions exclusively in Nipponbare or shared between HEG4 and Nipponbare were identified as parental *mPing* loci. These loci have allele frequencies of ~50% (35% to 55%) for 44 Nipponbare specific *mPing* loci or ~90% (89% to 95%) for 7 shared *mPing* loci. Second, 415 nonreference *mPing* insertions were identified from four independent sequenced HEG4 libraries (SRA accession number: SRR1619147, SRR833485, SRR833529, and SRR1619373). Determination of the allele frequency of the 415 *mPing* loci in the RILs revealed that they were present in more than 10% of the 272 RILs (ranging from 16% to 74%), suggesting these *mPing* loci were segregating in the population. Together, these 466 *mPing* loci were used as parental *mPing* loci in the analysis.

The recombination bin map and SNPs that distinguish the parental genotypes were used to phase the haplotypes of *mPing* loci in each RIL. Empty *mPing* loci within a Nipponbare or HEG4 haplotype were considered as potential excision sites and further confirmed by analyzing the alignments of reads covering the *mPing* insertion sites. Excisions were characterized only if there were two or more reads covering the 20 bp flanking an *mPing* insertion. Footprints of excision events were characterized by small insertion/deletions (indels) in the CIGAR alignments. Two approaches were applied to identify independent excision events: (*i*) a standard method that counts each excision as an independent event assuming that all RILs are independent; (*ii*) a conservative method that counts independent excisions from each locus according to the configuration of footprints in each RIL. Excisions that have the same footprint were counted as dependent events. A binomial test was used to test for significance of high excision frequency *mPing* insertions based on a frequency of 0.0051 excision events per locus per plant. Two methods identified similar sets of *mPing* loci with high frequency excision events (Supplementary Table 13). Excisions of high excision frequency *mPing* loci identified by both standard method and conservative method were manually inspected and confirmed by a PCR approach.

### Structural variation in the RILs and proximity to *mPing* elements

A bioinformatics approach was employed to detect potential sequence rearrangements in the RILs. Only deletions could be identified with confidence because of the limitations associated with the resolution of short reads (75 bp and 100 bp in this study) and small insertion size libraries (~200 bp in this study). CNVnator was used to analyze the read depth of bam alignment files in 272 RILs (28). A series of filters was used to obtain high-confident deletions from the raw results of CNVnator: (*i*) considering only these regions with normalized read depth less than 0.05 as candidate deletions; (*ii*) excluding centromeres, telomere chromosome ends and low coverage regions in Nipponbare and HEG4 genomes; (*iii*) any candidate regions should have at least one read supporting the junction of at least one end of deletions. Candidate structural variations were further analyzed by a PCR approach or sequencing with additional primers (Supplementary Data 4).

### Differential gene expression analysis

RNAseq reads from Nipponbare and HEG4 were obtained from SRA BioProject PRJNA264731. Each sample has three biological replicates with 51 nt single-end reads. Reads were aligned to MSUv7 with tophat v2.1.1 (52) using default parameters. Read counts of protein-coding genes were estimated by the htseq-count tool from HTSeq v0.7.2 (53). Differential gene expression analysis was performed with edgeR v2.99.8 (54) using the generalized linear model (GLM). Genes with FDR ≤ 0.05 were reported as differentially expressed genes.

### Statistical analysis

Pearson’s correlation test, Mann-Whitney test, one-way ANOVA and Tukey’s HSD test were performed in R with “cor.test”, “wilcox.test”, “aov”, and “TukeyHSD” functions. Samples sizes, statistical tests, and *P* values are described in figures or figure legends.

## Code availability

Scripts used in this study are available at https://github.com/stajichlab/Dynamic_rice_publications or http://doi.org/10.5281/zenodo.3662095.

## Data availability

The raw sequences have been deposited in the NCBI SRA project under the accession number SRP072364 or BioProject PRJNA316308.

## Acknowledgments

We thank Dr. Anna M. McClung from USDA for maintaining the RILs, and Dr. Venkateswari J. Chetty for preparation of plant materials and sequencing libraries. This work was supported by the National Science Foundation grant IOS-1027542 to S.R.W and J.E.S.

