## Supplementary information for "Genomic diversity generated by a transposable element burst in a rice recombinant inbred population"

Short title: *mPing* burst in a rice recombinant inbred population

Authors: Jinfeng Chen[a,b,1], Lu Lu[a,1], Sofia M.C. Robb[a,b], Matthew Collin[a,c], Yutaka Okumoto[d], Jason E. Stajich[b,c], and Susan R. Wessler[a,c,2]

Affiliations:

[a] Department of Botany and Plant Sciences, University of California, Riverside, CA 92521, USA.

[b] Department of Microbiology and Plant Pathology, University of California, Riverside, CA 92521, USA.

[c] Institute for Integrative Genome Biology, University of California, Riverside, CA 92521, USA.

[d] Graduate School of Agriculture, Kyoto University, Kitashirakawa-oiwake Sakyo, Kyoto 606-8502, Japan.

[1]J.C. and L.L. contributed equally to this work.

Keywords: *mPing*, active transposon, transposition, recombinant inbred lines, excision, sequence rearrangement, structural variation

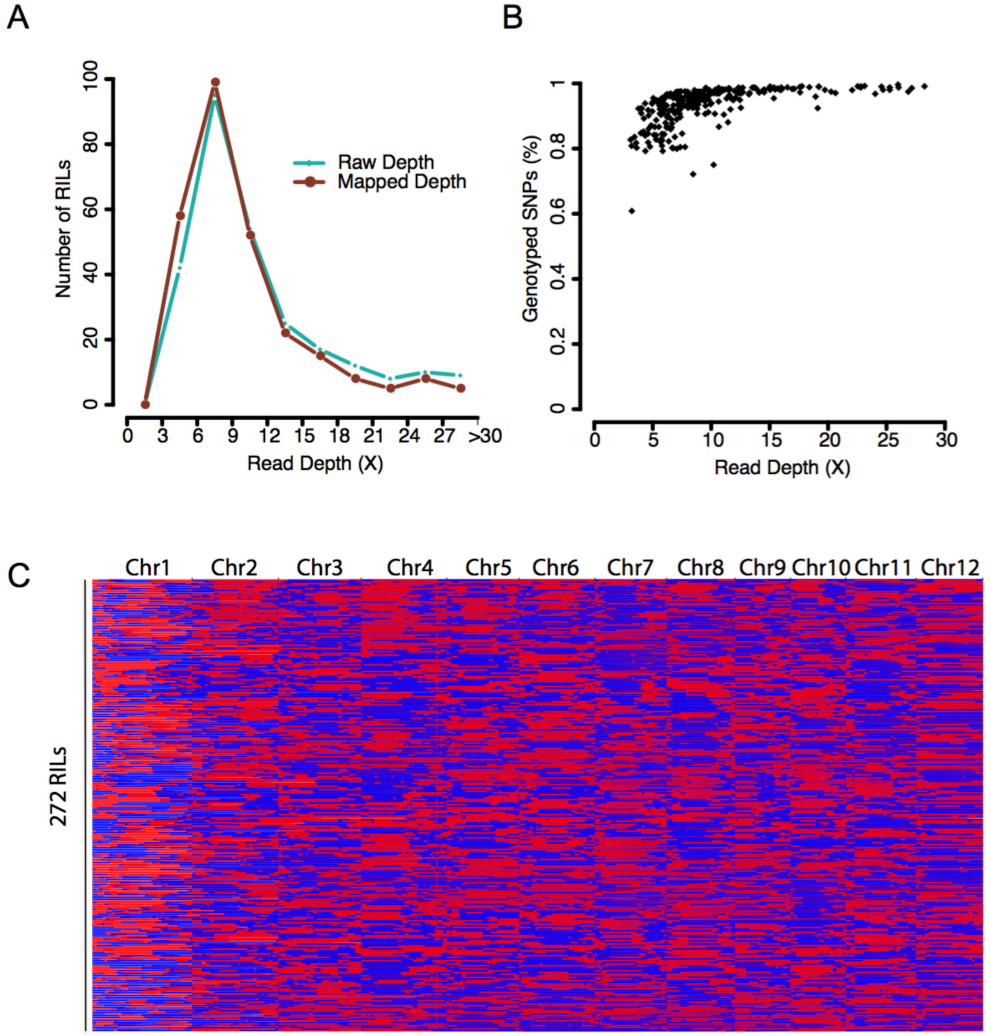


**Supplementary Figure 1. Sequencing features and genotypes of 272 RILs.** **A**, Sequence depth of 272 RILs. **B**, Genotyping rate of parental SNPs in the RILs. Reads were aligned to rice reference genome (MSUv7) by BWA and SNPs between parental strains were genotyped based on the alignments using GATK. **C**, Recombination map of RILs. Recombination bins were constructed using the high-quality genotype data in the RIL based on an HMM method. A comprehensive recombination map was constructed by imputation of missing makers and refining boundaries of the recombination bin using the R/qtl package. Blue and red represent parental accessions HEG4 and Nipponbare, respectively.

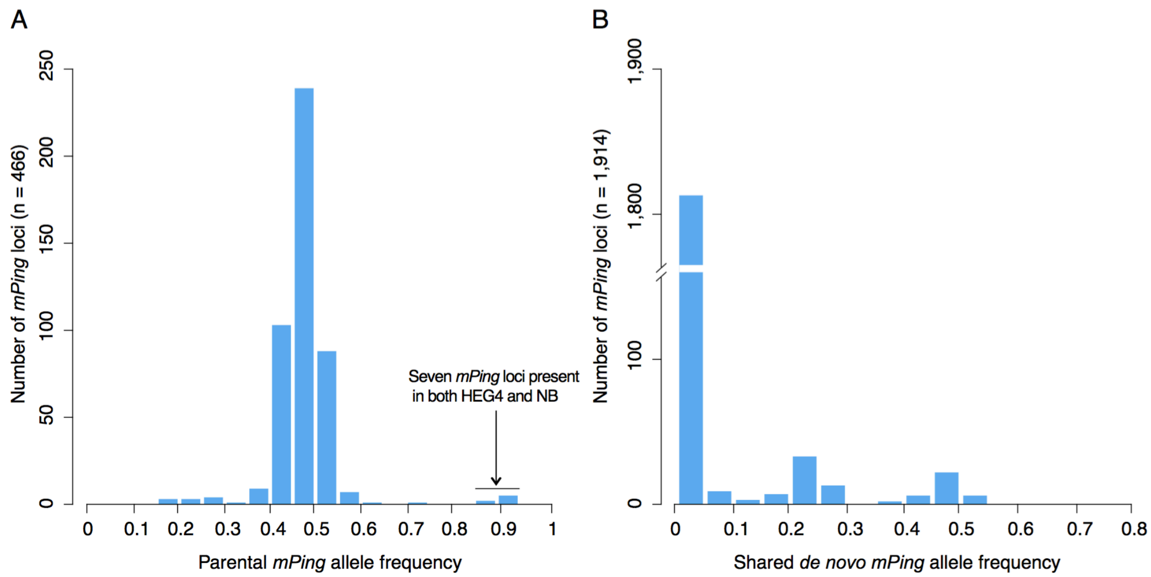


**Supplementary Figure 2. Allele frequency of parental and shared *mPing* loci. A**, Allele frequencies of parental *mPing* loci in 272 RILs. A total of 466 parental *mPing* loci (7 shared, 44 in NB, and 415 in HEG4) were used to calculate allele frequencies. **B**, Allele frequencies of shared *mPing* loci in 272 RILs. A total of 1,914 shared *de novo mPing* loci that are present in multiple RILs but not in HEG4 and NB were used to calculate the allele frequency.

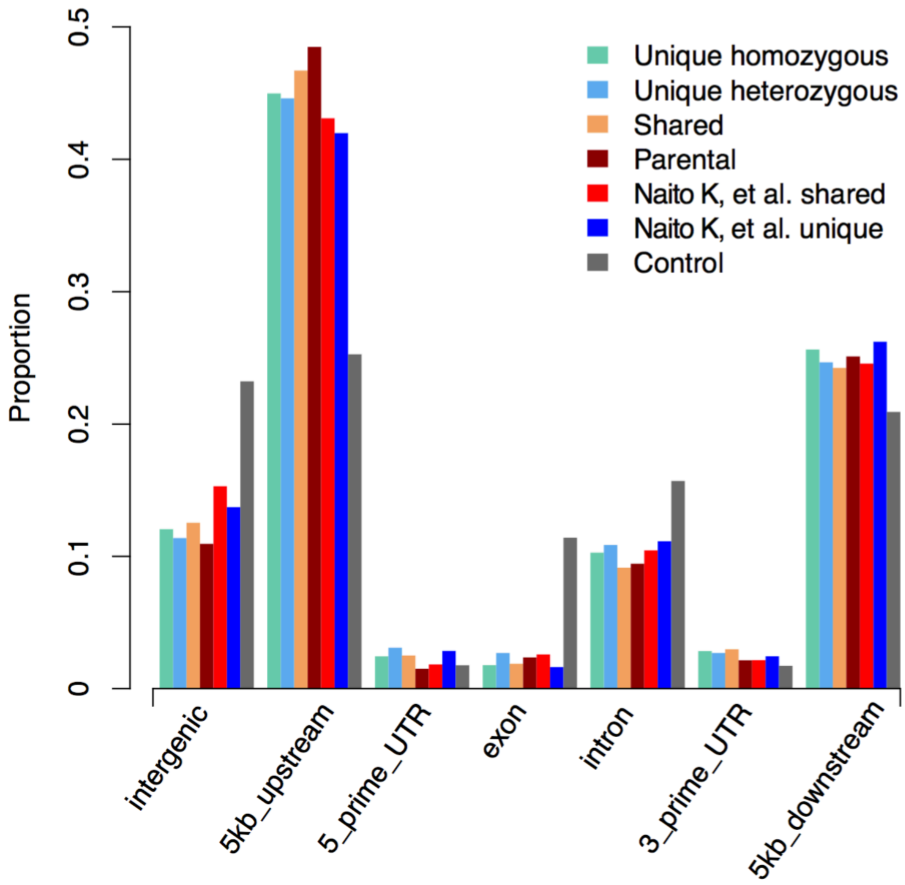


**Supplementary Figure 3. Distribution of *mPing* insertion sites in RILs compared to Naito K, et al, 2009.** *mPing* insertion sites are summarized according to their relative positions in and near protein-coding genes. Unique homozygous, unique heterozygous represent homozygous and heterozygous *mPing* insertion sites present exclusively in a single RIL. Shared represents insertion sites in multiple RILs but not in parents. Parental insertion sites are insertion sites shared in RILs and parents. Naito K, et al. shared are insertion sites in multiple individuals in Naito K, et al, 2009. Naito K, et al. unique are insertion sites in a single individual. Control is a random sampling from genomic sequence (n = 3,000).

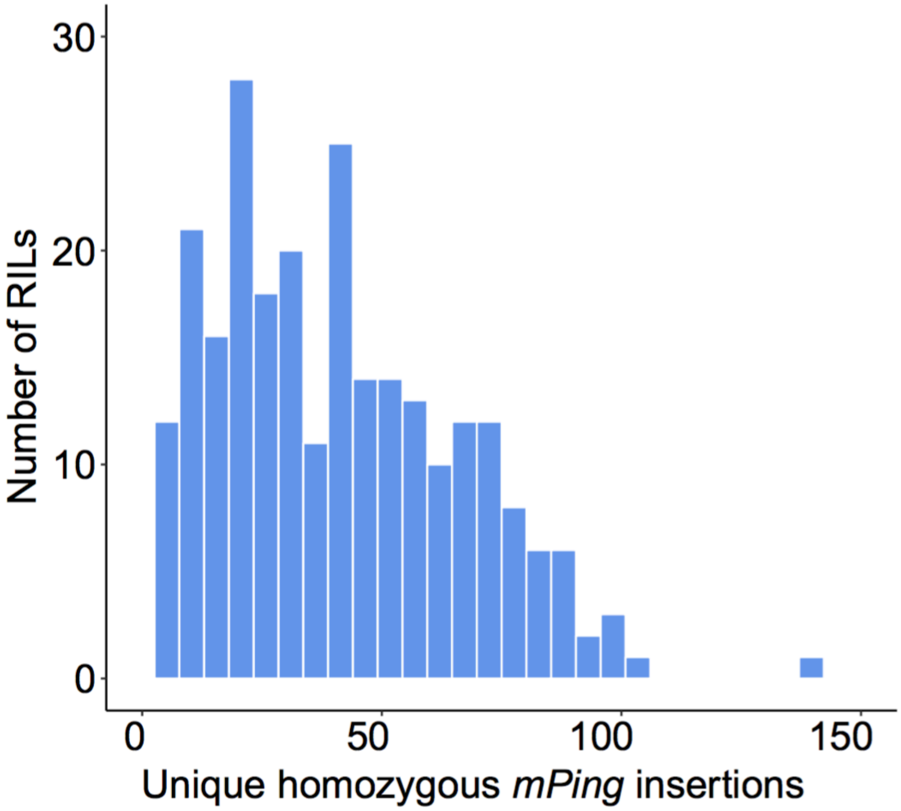


**Supplementary Figure 4. Distribution of the number of unique homozygous *mPing* insertions vs. the number of RILs with that number of new insertions.**

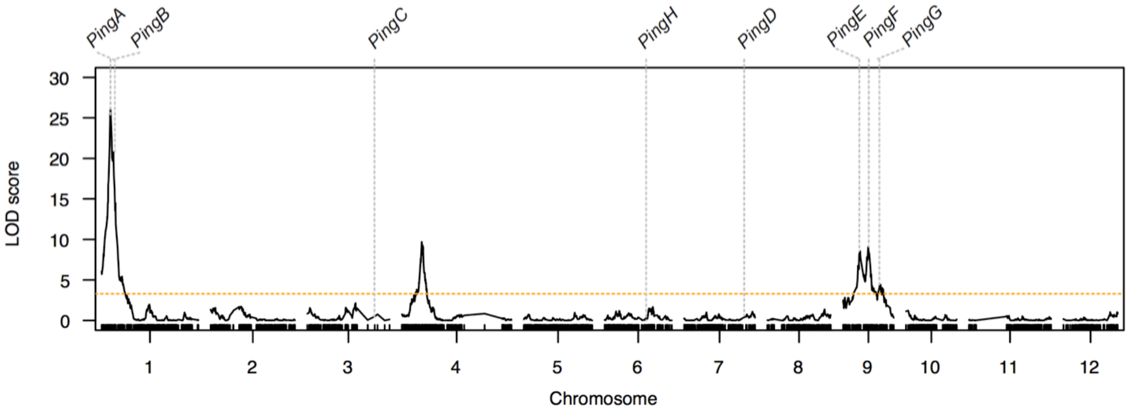


**Supplementary Figure 5. Mapped genetic loci impacting *mPing* transposition activity in the RILs.** Unique homozygous *mPing* insertions in RILs were used as the quantitative trait and recombination bins as markers. Markers along chromosomes are shown on the X axis and logarithm of the odds ratio (LOD score) of the QTL mapping along the Y axis. Orange dashed line indicates the threshold of the LOD score estimated by a permutation test (1,000 permutations, *P* = 0.05). *Ping* loci are shown (A -H).

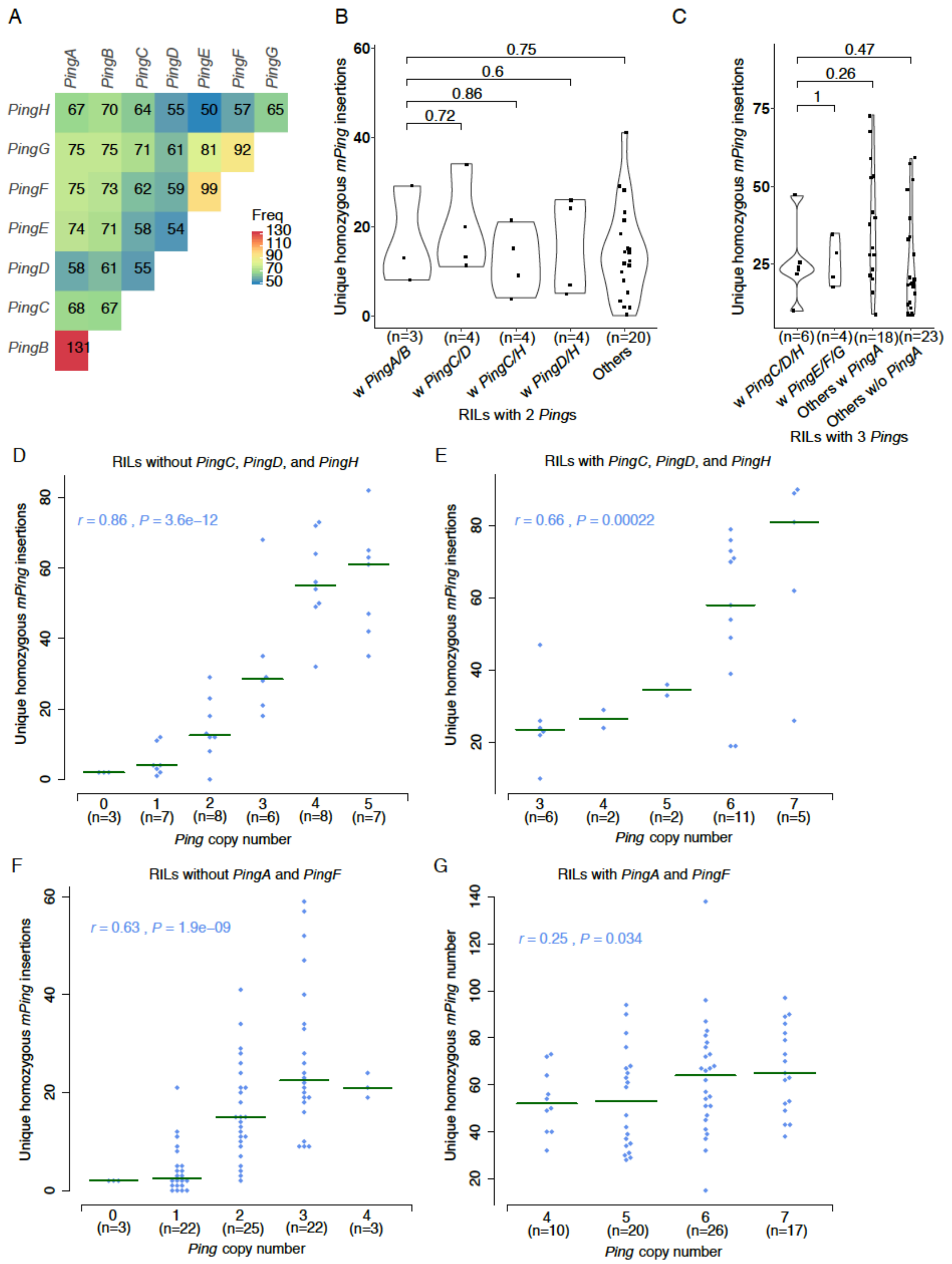


**Supplementary Figure 6. Comparison of unique homozygous *mPing* insertions in RILs with different *Ping* combinations. A,** Number of pairwise *Ping* combination in RILs. **B-C,** Comparison of dosage effect of *PingC*, *PingD*, *PingH* on *mPing* transposition in RILs with 2 *Ping*s (**B**) or 3 *Ping*s (**C**). The difference between groups was tested by Mann-Whitney test and indicated by *P* value. **D-E**, Correlation between unique homozygous *mPing* insertions and *Ping* copy number in the RILs without (**D**) or with *PingC*, *PingD*, and *PingH* (**E**). **F-G**, Correlation between unique homozygous *mPing* insertions and *Ping* copy number in the RILs without (**F**) or with *PingA* and *PingF* (**G**). The 272 RILs were grouped by *Ping* copy number ranging from 0 to 7 and the scatterplot shows the number of unique *mPing* insertions in each group. The number of RILs in each group is in parentheses (n=). Green lines are the group median. The significance of correlation was tested by a two-tailed Pearson's correlation test and indicated by *P* value.

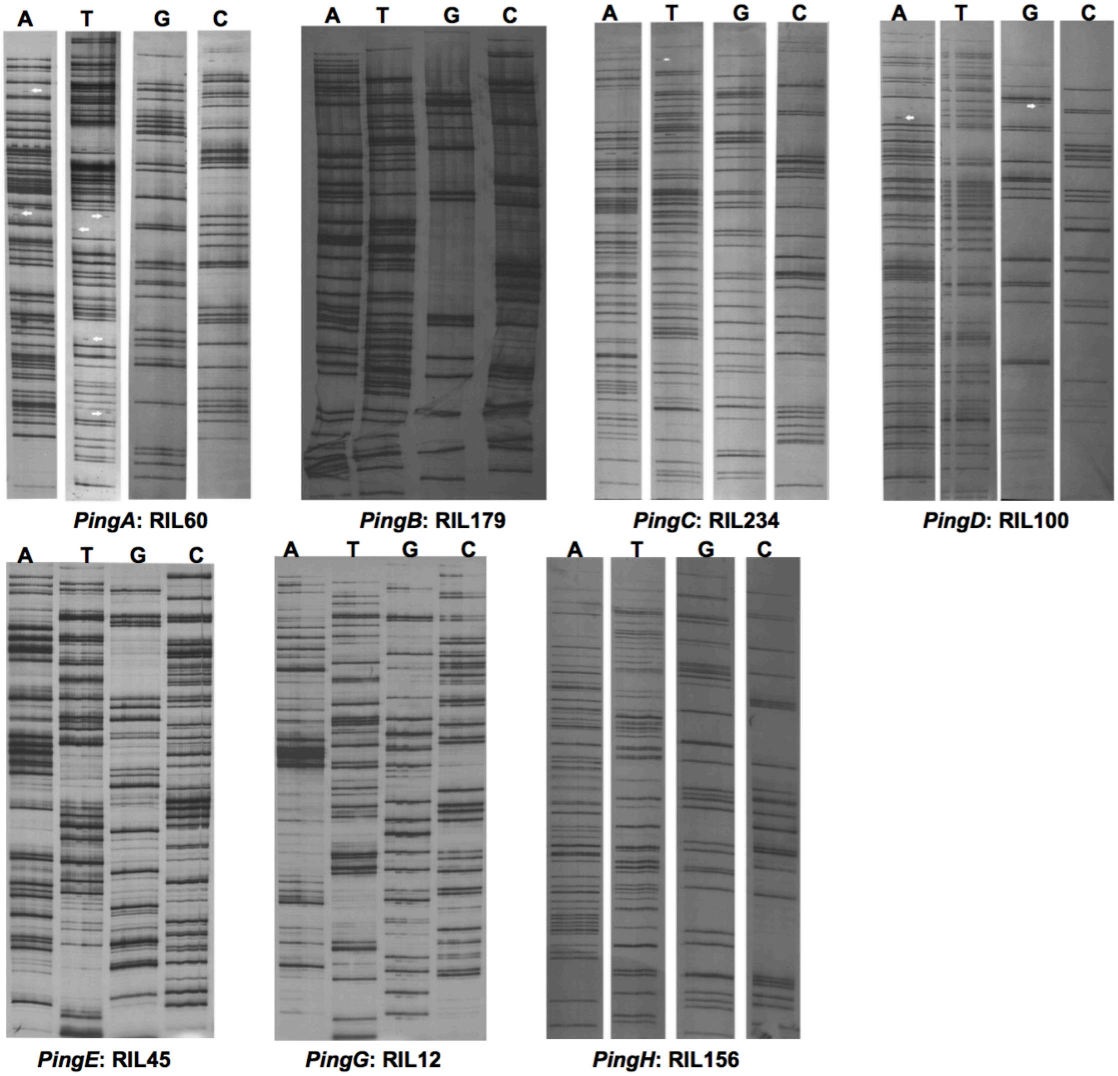


**Supplementary Figure 7. Transposon display analysis of *mPing* transposition in single-*Ping* RILs.** Genomic DNA of each plant was extracted and digested with *Mse*I, ligated with an adapter and two rounds of PCR amplification performed with preselective primers (*Mse*I + 0 and *mPing* P1) followed by four sets of selective primers (*Mse*I + A/T/G/C and *mPing* P2). Each panel contains 8 lanes – one for each plant analyzed. For each RIL, bands were compared within each selective primer set and new bands (indicated by white arrow) were counted if present in only one plant.

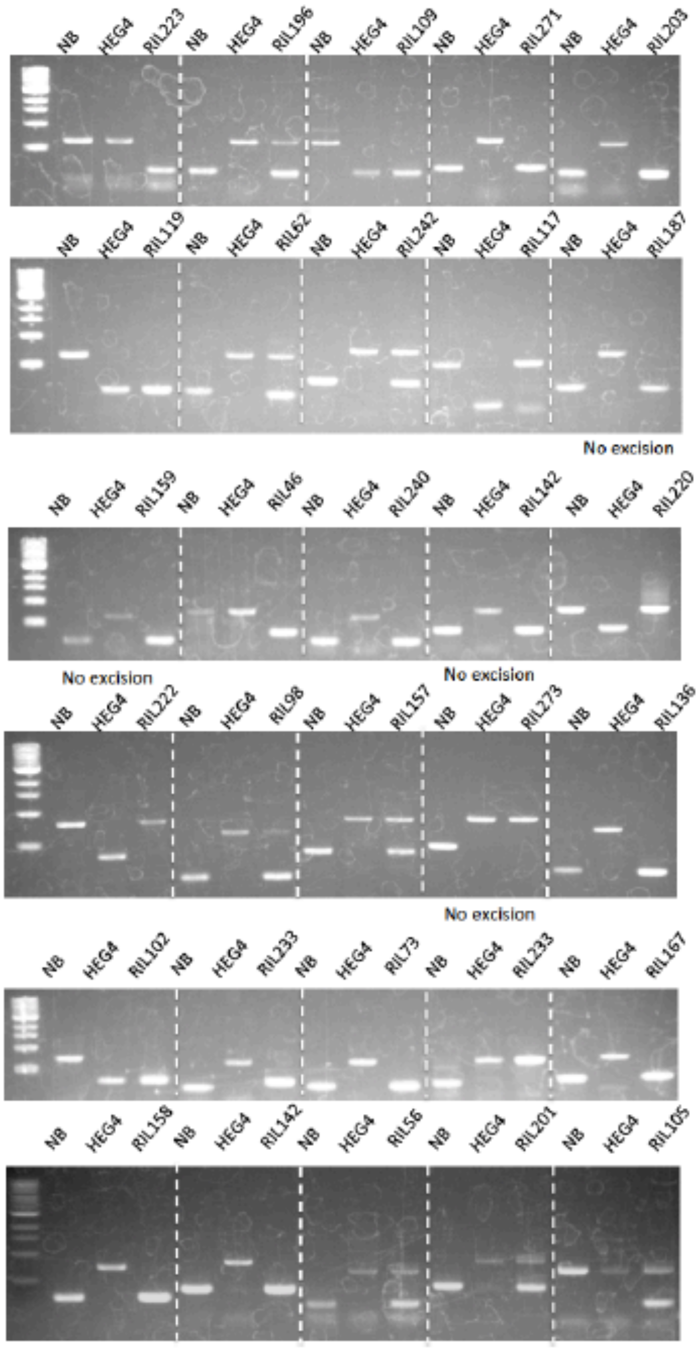


**Supplementary Figure 8.** PCR validation of 30 *mPing* excision events. Among excised *mPing* loci, 21 from HEG4 and 9 from NB (both specific and shared) were randomly selected to validate the accuracy of the excision identification method. Forward and reverse primers are in Supplementary Data 4.

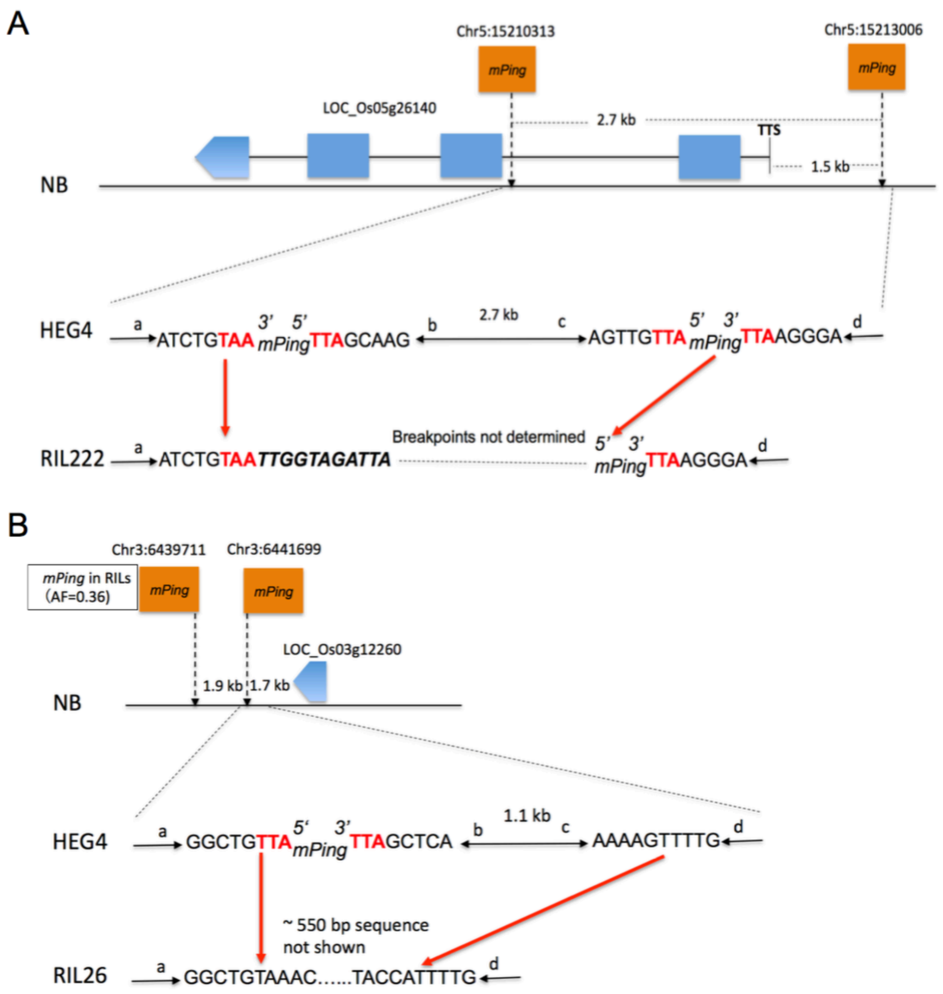


**Supplementary Figure 9. Structural variations mediated by *mPing* elements. A**, SV5. **B**, SV3. *mPing*-associated structural variations in select RILs were aligned with either Nipponbare (NB) or HEG4 based on the chromosome of origin. Lines indicate DNA fragments flanking *mPing* insertions or the breakpoint of structural variation. Letters (a-d) on lines label the DNA fragment end. Orange boxes indicate *mPing* insertions. Blue boxes indicate exons of protein-coding genes. Target site duplications (TSDs) of *mPing* insertions are highlighted in red. Filler DNA at breakpoints is italicized. Red arrows indicate the breakpoints where structural variation occurred. Allele frequency (AF) is indicated if *mPing* is present in RILs but absent in parental accessions.

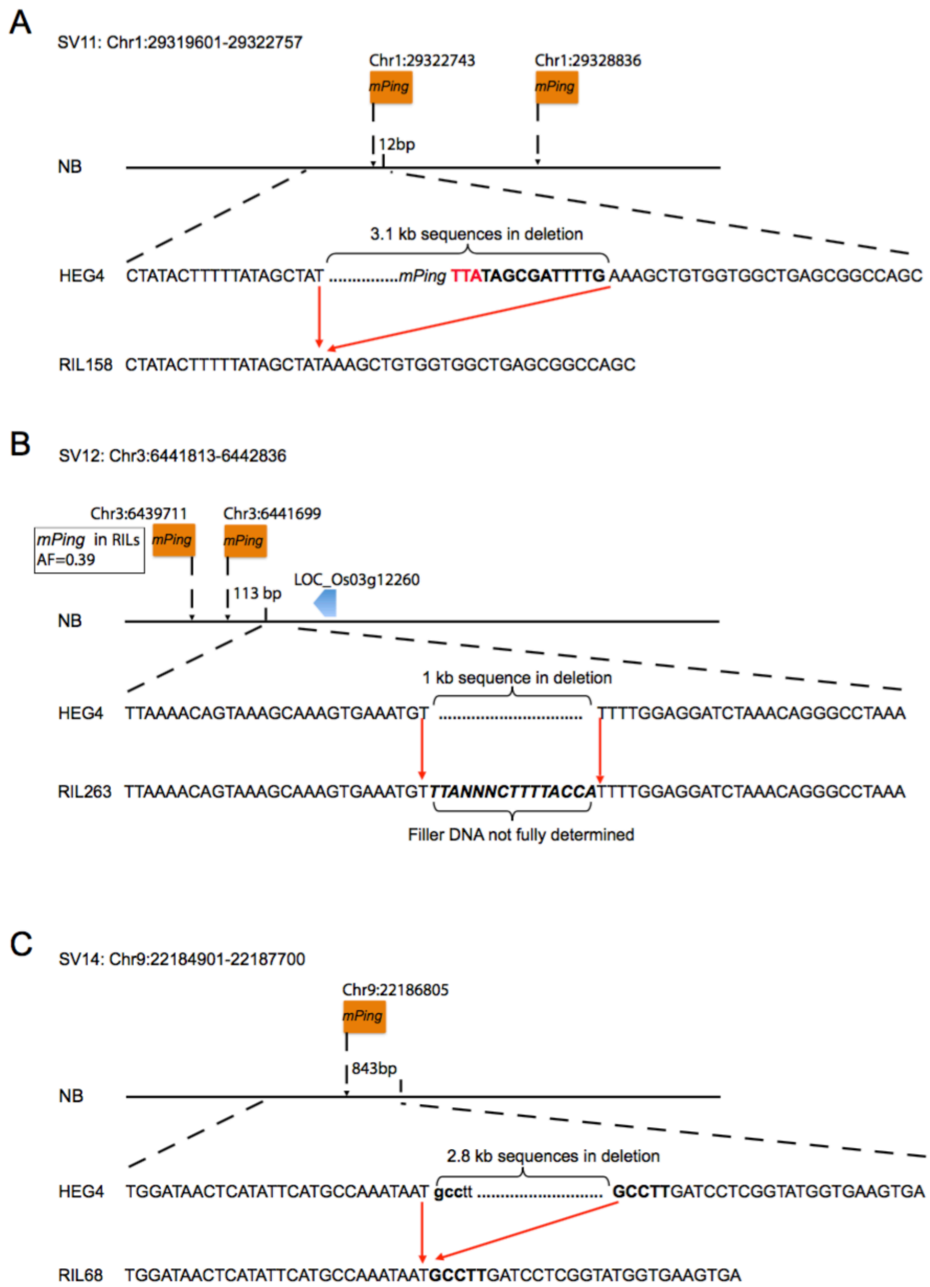


**Supplementary Figure 10. Structural variations with *mPing* elements nearby but not at breakpoints*.* A**, SV11. **B**, SV12. **C**, SV14. *mPing*-associated structural variations in select RILs were aligned with either Nipponbare (NB) or HEG4 based on the chromosome of origin. Orange boxes indicate *mPing* insertions. Blue boxes indicate exons of protein-coding genes. Target site duplications (TSDs) of *mPing* insertions are highlighted in red. Filler DNA at breakpoints are italicized. Red arrows indicate the breakpoints where structural variation occurred. Allele frequency (AF) is indicated if *mPing* is present in RILs but absent in parental accessions.

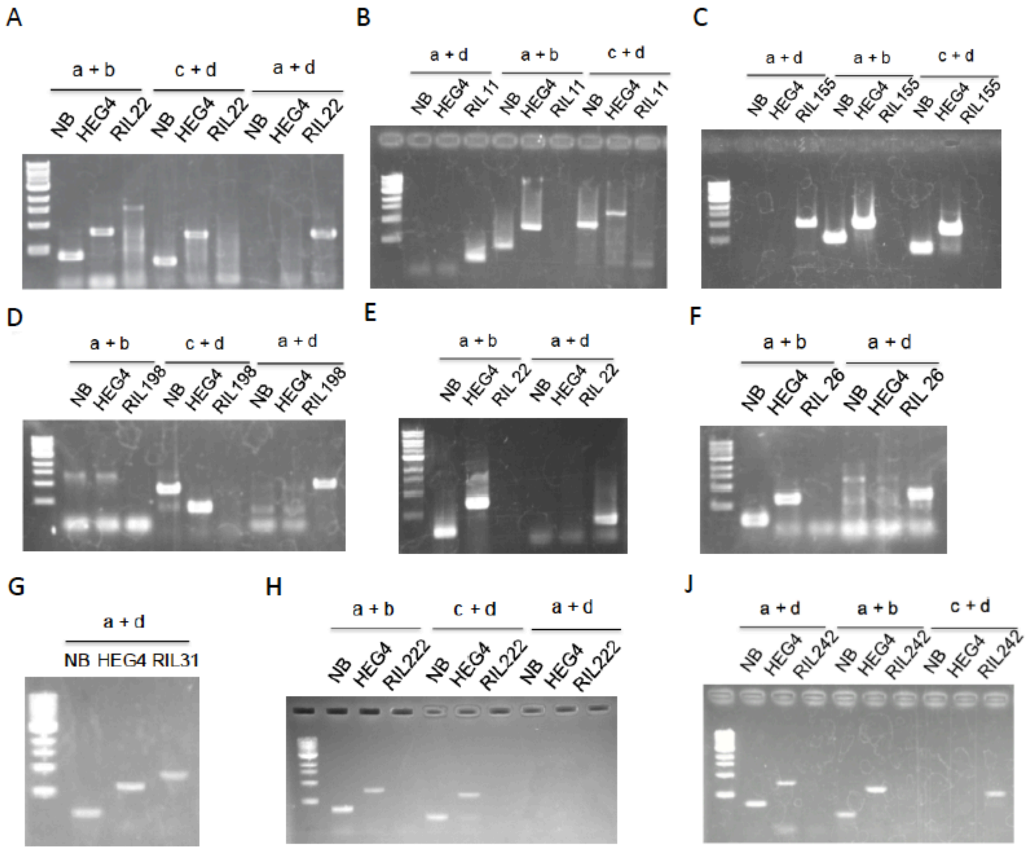


**Supplementary Figure 11.** PCR validation of *mPing*-associated sequence rearrangements shown in Figure 5 and Supplementary Figure 6. **A**, SV1. **B**, SV2. **C**, SV7. **D**, SV9. **E**, SV8. **F**, SV3. **G**, SV4. **H**, SV5. **J**, SV6. Forward and reverse primers are indicated by letters (a-d) as shown in Figure 5 and Supplementary Figure 6.

**Supplementary Table 1. Recombination bin map of 272 RILs**

| Chromosome | Number of bins | Average bin size (kb) | Average genetic distance (cM) | Total genetic distance (cM) | Physical length (Mb) | Recombination rate (cM/Mb) |
|---|---|---|---|---|---|---|
| Chr1 | 343 | 119.81 | 0.57 | 196.25 | 43.27 | 4.54 |
| Chr2 | 264 | 135.00 | 0.65 | 170.98 | 35.94 | 4.76 |
| Chr3 | 163 | 209.20 | 1.02 | 166.90 | 36.41 | 4.58 |
| Chr4 | 275 | 128.33 | 0.81 | 222.42 | 35.50 | 6.27 |
| Chr5 | 217 | 137.64 | 0.64 | 138.91 | 29.96 | 4.64 |
| Chr6 | 194 | 160.73 | 0.70 | 136.41 | 31.25 | 4.37 |
| Chr7 | 248 | 119.26 | 0.58 | 143.77 | 29.70 | 4.84 |
| Chr8 | 186 | 151.83 | 0.69 | 128.71 | 28.44 | 4.53 |
| Chr9 | 122 | 187.18 | 0.84 | 102.68 | 23.01 | 4.46 |
| Chr10 | 175 | 130.29 | 0.59 | 102.82 | 23.21 | 4.43 |
| Chr11 | 220 | 131.89 | 0.76 | 166.42 | 29.02 | 5.73 |
| Chr12 | 165 | 166.82 | 0.67 | 110.12 | 27.53 | 4.00 |
| Mean (Total) | 214 (2572) | 142.76 | 0.69 | 148.86 (1786) | 31.10 (373) | 4.79 |

**Supplementary Table 2. Genotypes of SNPs flanking *de novo mPing* insertion sites**

| Flanking SNP genotype[a] | Heterozygous *mPing*s | Homozygous *mPing*s |
|---|---|---|
| hom\|hom | 3129 | 8361 |
| -NB | 1312 | 3500 |
| -HEG4 | 1429 | 3700 |
| -NB+HEG4 | 123 | 315 |
| -Not determined | 265 | 846 |
| hom\|het | 89 | 262 |
| het\|het | 11 | 6 |
| Not determined | 778 | 1898 |
| Total | 4007 | 10527 |

[a], hom|hom is where SNPs flanking both ends of *mPing* are homozygous. hom+het is where SNPs flanking one end of *mPing* are homozygous while the other end is heterozygous. het|het is where SNPs flanking both ends of *mPing* are heterozygous.

**Supplementary Table 3.** ***de novo Ping*** **insertions**

| New *Ping* locus | RIL | New homozygous (heterozygous) *mPing* | Nearest protein-coding genes | Location relative to closest protein-coding gene | Chromatin environment | All *Ping* loci in RIL where N = new *Ping* |
|---|---|---|---|---|---|---|
| Chr3:865992 | 3 | 43 (4) | LOC_Os03g02430.1 | -3215 | Euchromatin | 7-NABCEFG |
| Chr1:34194379 | 14 | 48 (29) | LOC_Os01g59190.1 | 210 | Euchromatin | 6-NCEFGH |
| Chr8:18337647 | 26 | 42 (21) | LOC_Os08g29809.1 | -7734 (intergenic) | TE island | 5-NDEFG |
| Chr2:28343688 | 46 | 73 (46) | LOC_Os02g46490.1 | -2292 | Euchromatin | 7-NABEFGH |
| Chr2:1188028 | 51 | 67 (25) | LOC_Os02g03020.1 | -86 | Euchromatin | 6-NABDEF |
| Chr1:7258553 | 65 | 62 (18) | LOC_Os01g13040.1 | -1231 | Euchromatin | 6-NABDEF |
| Chr3:36131662 | 93 | 21 (9) | LOC_Os03g63940.1 | Intron | Euchromatin | 4-NBGH |
| Chr7:5440360 | 104 | 44 (7) | LOC_Os07g10130.1 | Intron | Euchromatin | 4-NABH |
| Chr10:13635022 | 149 | 52 (15) | LOC_Os10g26290.1 | -20820 (intergenic) | TE island | 3-NGH |
| Chr1:23216637 | 174 | 43 (15) | LOC_Os01g41000.1 | -1340 | Euchromatin | 7-NABDEFH |
| Chr5:4075298 | 175 | 81 (23) | LOC_Os05g07600.1 | -707 | Euchromatin | 7-NABCDGH |
| Chr9:18957569 | 189 | 38 (10) | LOC_Os09g31458.1 | 76 | Euchromatin | 8-NABCEFGH |
| Chr11:22264027 | 211 | 73 (13) | LOC_Os11g37680.1 | Intron | Euchromatin | 5-NABDH |
| Chr7:22688057 | 221 | 70 (60) | LOC_Os07g37830.1 | -841 | Euchromatin | 7-NABEFGH |
| Chr12:7386029 | 248 | 57 (20) | LOC_Os12g13280.1 | Intron | Euchromatin | 6-NABFGH |
| Chr11:4473183 | 270 | 96 (36) | LOC_Os11g08470.1 | 3436 | TE island | 6-NNEFGH |
| Chr11:5885534 | 270 | 96 (36) | LOC_Os11g10720.1 | -4005 | Euchromatin | 6-NNEFGH |

**Supplementary Table 4. QTL loci contributing to *mPing* activity**

| Chromosome | Genetic map position | Genetic map interval | Interval on chromosome | Fitted LOD | Variance explained by QTL | QTL effect | Standard deviation of QTL effect |
|---|---|---|---|---|---|---|---|
| Chr1 | 17.92 | 17.18-20.40 | 2496808-3672310 | 29.65 | 28.19 | 14.02 | 1.06 |
| Chr4 | 40.28 | 38.54-43.42 | 15520048-16724166 | 14.27 | 11.82 | 8.99 | 1.05 |
| Chr9 | 50.95 | 31.88-54.16 | 10571059-14581359 | 10.93 | 8.79 | 7.81 | 1.06 |

**Supplementary Table 5. *mPing* transposition estimated by the transposon display approach**

| *Ping* | RIL | Number of individuals | New *mPing* insertions | New *mPing* insertions per plant |
|---|---|---|---|---|
| *PingA* | RIL60 | 8 | 6 | 0.75 |
| *PingA* | RIL60 | 8 | 8 | 1 |
| *PingA* | RIL60 | 8 | 4 | 0.5 |
| *PingB* | RIL179 | 8 | 0 | 0 |
| *PingB* | RIL179 | 8 | 0 | 0 |
| *PingB* | RIL179 | 8 | 0 | 0 |
| *PingC* | RIL230 | 8 | 0 | 0 |
| *PingC* | RIL230 | 8 | 0 | 0 |
| *PingC* | RIL230 | 8 | 0 | 0 |
| *PingC* | RIl234 | 8 | 1 | 0.125 |
| *PingC* | RIl234 | 8 | 2 | 0.25 |
| *PingC* | RIl234 | 8 | 3 | 0.375 |
| *PingD* | RIL75 | 8 | 3 | 0.375 |
| *PingD* | RIL75 | 8 | 2 | 0.25 |
| *PingD* | RIL75 | 8 | 2 | 0.25 |
| *PingD* | RIL100 | 8 | 2 | 0.25 |
| *PingD* | RIL100 | 8 | 2 | 0.25 |
| *PingD* | RIL100 | 8 | 2 | 0.25 |
| *PingE* | RIL45 | 8 | 0 | 0 |
| *PingE* | RIL45 | 8 | 0 | 0 |
| *PingE* | RIL45 | 8 | 0 | 0 |
| *PingE* | RIL268 | 8 | 2 | 0.25 |
| *PingE* | RIL268 | 8 | 1 | 0.125 |

| | | | | |
|---|---|---|---|---|
| *PingE* | RIL268 | 8 | 1 | 0.125 |
| *PingG* | RII12 | 8 | 0 | 0 |
| *PingG* | RII12 | 8 | 0 | 0 |
| *PingG* | RII47 | 8 | 2 | 0.25 |
| *PingG* | RII47 | 8 | 2 | 0.25 |
| *PingG* | RII47 | 8 | 0 | 0 |
| *PingH* | RII156 | 8 | 0 | 0 |
| *PingH* | RII156 | 8 | 0 | 0 |
| *PingH* | RII166 | 8 | 0 | 0 |
| *PingH* | RII166 | 8 | 0 | 0 |
| *PingH* | RII166 | 8 | 0 | 0 |

**Supplementary Table 6. *mPing* transposition estimated by the pooled-sequencing approach**

| *Ping* | RIL | Replicate | Number of individuals | New *mPing* insertions | New *mPing* insertions per plant |
|---|---|---|---|---|---|
| *PingA* | RIL60 | Rep1 | 8 | 2 | 0.25 |
| *PingA* | RIL60 | Rep2 | 8 | 6 | 0.75 |
| *PingA* | RIL60 | Rep3 | 8 | 11 | 1.375 |
| *PingA* | RIL60 | Rep4 | 8 | 12 | 1.5 |
| *PingC* | RIL230 | Rep1 | 8 | 0 | 0 |
| *PingC* | RIL230 | Rep2 | 8 | 0 | 0 |
| *PingC* | RIL230 | Rep3 | 8 | 0 | 0 |
| *PingD* | RIL99 | Rep1 | 8 | 1 | 0.125 |
| *PingD* | RIL99 | Rep2 | 8 | 0 | 0 |
| *PingD* | RIL99 | Rep3 | 8 | 0 | 0 |
| *PingE* | RIL268 | Rep1 | 8 | 2 | 0.25 |
| *PingE* | RIL268 | Rep2 | 8 | 0 | 0 |
| *PingE* | RIL268 | Rep3 | 9 | 0 | 0 |
| *PingG* | RIL12 | Rep1 | 8 | 0 | 0 |
| *PingG* | RIL12 | Rep2 | 8 | 1 | 0.125 |
| *PingG* | RIL12 | Rep3 | 8 | 0 | 0 |
| *PingH* | RIL156 | Rep1 | 8 | 0 | 0 |
| *PingH* | RIL156 | Rep2 | 8 | 0 | 0 |
| *PingH* | RIL156 | Rep3 | 7 | 1 | 0.142857 |

**Supplementary Table 7. Distance from high frequency excised *mPing* to nearest *mPing* insertion**

| *mPing* locus | # Excision event | Nearest *mPing* locus in HEG4 (kb) | Nearest *mPing* locus in RILs (kb)[a] |
|---|---|---|---|
| Chr1:29494572 | 5 | 165.7 | 165.7 |
| Chr1:3444992 | 5 | 335.8 | 335.8 |
| Chr1:36267659 | 5 | 2.8 | 2.8 |
| Chr1:36270511 | 7 | 2.8 | 2.8 |
| Chr1:6806761 | 7 | 9.6 | 9.6 |
| Chr2:30934395 | 6 | 520.9 | 22.3 |
| Chr3:6441699 | 8 | 70.3 | 1.9 |
| Chr5:15210313 | 9 | 2.6 | 2.6 |
| Chr5:15213006 | 10 | 2.6 | 2.6 |
| Chr6.8316574 | 5 | 40 | 40 |
| Chr8:13076782 | 6 | 26.3 | 26.3 |
| Chr8:24622994 | 15 | 37.6 | 2.3 |

[a], 92 *mPing* loci are not parental but found in 10% or more of the RILs. These loci were also used to calculate the distance of *mPing* loci to their closest loci.

**Supplementary Table 8. Nearest protein-coding gene from high frequency excised *mPing***

| *mPing* | Nearest protein-coding gene | Location relative to closest protein-coding gene | Gene expression difference (FDR) |
|---|---|---|---|
| Chr1:29494572 | LOC_Os01g51290.1 | -2016 | 1 |
| Chr1:3444992 | LOC_Os01g07300.1 | -165 | 0 |
| Chr1:36267659 | LOC_Os01g62630.1 | 402 | 1 |
| Chr1:36270511 | LOC_Os01g62630.1 | -774 | 1 |
| Chr1:6806761 | LOC_Os01g12440.1 | -6951 | 0.95 |
| Chr2:30934395 | LOC_Os02g50640.1 | -104 | 0.75 |
| Chr3:6441699 | LOC_Os03g12260.1 | 1752 | 1 |
| Chr5:15210313 | LOC_Os05g26140.1 | Intron | 1 |
| Chr5:15213006 | LOC_Os05g26140.1 | -576 | 1 |
| Chr6:8316574 | LOC_Os06g16740.1 | Intron | 1 |
| Chr8:13076782 | LOC_Os08g21840.1 | 721 | 1 |
| Chr8:24622994 | LOC_Os08g38960.1 | -162 | 0.2 |

**Supplementary Table 9. Features of structural variations in RILs**

| SV | Chromosome | Start | End | RIL[a] | SV Type | Length (bp) |
|---|---|---|---|---|---|---|
| SV1 | Chr1 | 29494474 | 29496305 | 22 | Deletion | 1832 |
| SV2 | Chr2 | 28724476 | 28732623 | 11 | Deletion | 8148 |
| SV3 | Chr3 | 6441699 | 6442836 | 26 | Deletion | 1138 |
| SV4 | Chr3 | 6441699 | 6442399 | 31 | Deletion + Duplication | 701 |
| SV5 | Chr5 | 15210313 | 15213006 | 222 | Deletion | 2694 |
| SV6 | Chr5 | 15210313 | 15213006 | 242 | Deletion | 2694 |
| SV7 | Chr8 | 24622994 | 24625306 | 155 | Deletion | 2313 |
| SV8 | Chr8 | 24625306 | 24634547 | 22 | Deletion | 9242 |
| SV9 | Chr9 | 638901 | 694912 | 198 | Deletion | 56012 |
| SV10 | Chr1 | 10145801 | 10151500 | 274 | Deletion | 5700 |
| SV11 | Chr1 | 29319601 | 29322757 | 158 | Deletion | 3157 |
| SV12 | Chr3 | 6441813 | 6442836 | 263 | Deletion | 1024 |
| SV13 | Chr7 | 10749401 | 10760200 | 61 | Deletion | 10800 |
| SV14 | Chr9 | 22184901 | 22187700 | 68 | Deletion | 2800 |
| SV15 | Chr10 | 13066801 | 13068300 | 78 | Deletion | 1500 |
| SV16 | Chr10 | 17891753 | 17895900 | 78 | Deletion | 4148 |

[a], RILs where structural variations were characterized and validated

**Supplementary Table 10. Presence or absence of structural variations in F1 progeny**

| SV | Location | # of RILs | F1 #26 progeny (RIL1-139) | F1 #27 progeny (RIL140-280) |
|---|---|---|---|---|
| SV1 | Chr1:29494474-29496305 | 1 | 1 | 0 |
| SV2 | Chr2:28724476-28732623 | 1 | 1 | 0 |
| SV3 | Chr3:6441699-6442399 | 1 | 1 | 0 |
| SV4 | Chr3:6441699-6442836 | 1 | 1 | 0 |
| SV5 | Chr5:15210313-15213006 | 1 | 0 | 1 |
| SV6 | Chr5:15210313-15213006 | 1 | 0 | 1 |
| SV7 | Chr8:24622994-24625306 | 1 | 0 | 1 |
| SV8 | Chr8:24625306-24634547 | 1 | 1 | 0 |
| SV9 | Chr9:638901-694912 | 1 | 0 | 1 |
| SV10 | Chr1:10145801-10151500 | 67 | 0 | 67 |
| SV11 | Chr1:29319601-29322757 | 73 | 0 | 73 |
| SV12 | Chr3:6441813-6442836 | 1 | 0 | 1 |
| SV13 | Chr7:10749401-10760200 | 1 | 1 | 0 |
| SV14 | Chr9:22184901-22187700 | 1 | 1 | 0 |
| SV15 | Chr10:13066801-13068300 | 1 | 1 | 0 |
| SV16 | Chr10:17891753-17895900 | 1 | 1 | 0 |

**Supplementary Table 11. SVs that do/do not include high excision *mPings***

| *mPing* locus[a] | High frequency excision *mPing* | Structural variation-associated *mPing* |
|---|---|---|
| Chr1:29494572 (SV1) | + | + |
| Chr1:3444992 | + | - |
| Chr1:36267659 | + | - |
| Chr1:36270511 | + | - |
| Chr1:6806761 | + | - |
| Chr2:30934395 | + | - |
| Chr3:6441699 (SV3 & SV4) | + | + |
| Chr5:15210313 (SV5 & SV6) | + | + |
| Chr5:15213006 (SV5 & SV6) | + | + |
| Chr6:8316574 | + | - |
| Chr8:13076782 | + | - |
| Chr8:24622994 (SV7) | + | + |
| Chr8:24625306 (SV7 & SV8)[b] | - | + |
| Chr1:29496305 (SV1)[c] | - | + |
| Chr8:28724476 (SV2) | - | + |
| Chr8:28732623 (SV2) | - | + |
| Chr3:6439711 (SV3 & SV4)[d] | - | + |
| Chr9:694483 (SV9) | - | + |

[a], SVs in parentheses have *mPing* at breakpoints
[b], *mPing* locus Chr8:24625306 is in 97 RILs but not parents.
[c], *mPing* locus Chr1:29496305 is characterized by PCR (Supplementary Figure 7A) but is not detected by RelocaTE2.
[d], *mPing* locus Chr3:6439711 is in 107 RILs but not parents. It has 11 excision events and 10 of 11 with footprints.

**Supplementary Table 12. Protein-coding genes associated with SVs**

| Genes affected by structural variation | Gene annotation | Affected regions | Associated structural variation |
|---|---|---|---|
| LOC_Os01g18110 | cinnamoyl CoA reductase | exon and intron | SV10 |
| LOC_Os01g18120 | cinnamoyl CoA reductase | exon and intron | SV10 |
| LOC_Os01g51040 | transmembrane protein 16K | exon and intron | SV11 |
| LOC_Os02g47060 | WRKY66 | full gene | SV2 (*mPing* associated) |
| LOC_Os05g26140 | expressed protein | exon and intron | SV5 & SV6 (*mPing* associated) |
| LOC_Os07g18154 | aldehyde oxidase | exon and intron | SV13 |
| LOC_Os08g38970 | transmembrane receptor | exon | SV8 (*mPing* associated) |
| LOC_Os09g01960 | MYB family transcription factor | full gene | SV9 (*mPing* associated) |
| LOC_Os09g01970 | expressed protein | full gene | SV9 (*mPing* associated) |
| LOC_Os09g02000 | expressed protein | full gene | SV9 (*mPing* associated) |

**Supplementary Table 13. *mPing* excision events detected by conservative vs. standard methods**

| *mPing* | Conservative method | | Standard method | | Shared high frequency excision *mPing* |
|---|---|---|---|---|---|
| | Excision events | Event with footprint | Excision events | Event with footprint | |
| Chr8:24622994 | 15 | 14 | 28 | 14 | + |
| Chr5:15213006 | 10 | 9 | 14 | 9 | + |
| Chr5:15210313 | 9 | 8 | 18 | 9 | + |
| Chr3:6441699 | 8 | 7 | 19 | 8 | + |
| Chr1:36270511 | 7 | 6 | 19 | 7 | + |
| Chr1:6806761 | 7 | 6 | 10 | 9 | + |
| Chr2:30934395 | 6 | 5 | 8 | 5 | + |
| Chr8:13076782 | 6 | 5 | 6 | 5 | + |
| Chr1:36267659 | 5 | 4 | 18 | 4 | + |
| Chr1:3444992 | 5 | 4 | 8 | 4 | + |
| Chr1:29494572 | 5 | 4 | 7 | 4 | + |
| Chr6:8316574 | 5 | 5 | 6 | 6 | + |
| Chr2:14601853 | 4 | 3 | 11 | 3 | - |
| Chr2:28732623 | 4 | 3 | 6 | 5 | - |
| Chr7:29475470 | 4 | 3 | 6 | 4 | - |
| Chr2:30104937 | 3 | 2 | 6 | 3 | - |
| Chr5:23730117 | 3 | 2 | 6 | 2 | - |
| Chr9:7117395 | 3 | 2 | 6 | 3 | - |
| Chr7:25788368 | 2 | 1 | 10 | 1 | - |
| Chr5:23730902 | 2 | 1 | 6 | 1 | - |
| Chr4:33507343 | 2 | 1 | 6 | 3 | - |

Supplementary Data 1. Sequence and genotype quality of 272 RILs

| RIL ID | Number of reads | Read depth | Insert size (bp) | Genotyped SNPs (%) | Genome coverage (%) |
|---|---|---|---|---|---|
| 1 | 23383042 | 6.25 | 132 | 87.90 | 91.98 |
| 2 | 19649656 | 5.25 | 133 | 84.20 | 86.56 |
| 3 | 24519973 | 6.55 | 140 | 91.63 | 94.10 |
| 4 | 21499503 | 5.74 | 136 | 90.76 | 93.01 |
| 5 | 25126024 | 6.71 | 182 | 96.12 | 95.79 |
| 7 | 23157591 | 6.18 | 225 | 95.66 | 96.13 |
| 8 | 27594542 | 7.36 | 168 | 94.71 | 95.76 |
| 9 | 22707096 | 6.00 | 148 | 92.91 | 94.58 |
| 10 | 24009328 | 6.37 | 179 | 89.71 | 91.26 |
| 11 | 32377954 | 8.62 | 167 | 95.01 | 95.98 |
| 12 | 61108623 | 16.29 | 127 | 95.44 | 97.88 |
| 13 | 93999385 | 25.11 | 193 | 98.05 | 98.34 |
| 14 | 22645665 | 6.04 | 148 | 90.81 | 93.00 |
| 15 | 26750643 | 7.13 | 232 | 97.10 | 96.76 |
| 16 | 83610864 | 22.36 | 189 | 97.96 | 98.29 |
| 17 | 22677406 | 6.04 | 242 | 93.51 | 95.65 |
| 18 | 15322523 | 4.08 | 249 | 90.86 | 91.46 |
| 19 | 18847153 | 5.03 | 282 | 91.46 | 91.44 |
| 20 | 26666201 | 7.11 | 162 | 96.37 | 97.24 |
| 21 | 17508984 | 4.67 | 207 | 91.83 | 92.88 |
| 22 | 20275855 | 5.41 | 245 | 94.89 | 94.51 |
| 23 | 25951322 | 6.92 | 233 | 96.00 | 96.16 |
| 24 | 39005965 | 10.41 | 201 | 97.79 | 97.70 |
| 25 | 43441599 | 11.61 | 236 | 88.08 | 96.35 |
| 26 | 31456349 | 8.40 | 180 | 93.84 | 95.10 |
| 28 | 29421027 | 7.79 | 183 | 94.80 | 98.20 |
| 29 | 32682445 | 8.71 | 223 | 97.28 | 97.38 |
| 30 | 21871601 | 5.83 | 197 | 94.38 | 94.75 |
| 31 | 21698329 | 5.79 | 188 | 92.61 | 92.77 |
| 33 | 44575202 | 11.90 | 159 | 95.38 | 95.79 |
| 34 | 35150850 | 9.38 | 190 | 95.22 | 95.18 |
| 35 | 27468878 | 7.34 | 167 | 91.98 | 92.41 |
| 36 | 26072772 | 6.96 | 198 | 95.57 | 95.43 |
| 37 | 30617147 | 8.17 | 155 | 94.83 | 95.76 |
| 38 | 26079322 | 6.96 | 150 | 91.48 | 93.34 |
| 39 | 213561900 | 56.82 | 189 | 98.67 | 97.11 |
| 40 | 42973247 | 11.50 | 143 | 95.39 | 97.11 |
| 42 | 37841623 | 10.10 | 166 | 96.18 | 96.89 |
| 43 | 66951148 | 17.91 | 163 | 97.26 | 98.38 |

| | | | | | |
|---|---|---|---|---|---|
| 44 | 28406185 | 7.58 | 174 | 95.60 | 96.56 |
| 45 | 76868954 | 20.55 | 189 | 97.47 | 98.13 |
| 46 | 32165478 | 8.58 | 166 | 94.86 | 96.21 |
| 47 | 76637452 | 20.07 | 172 | 95.95 | 98.61 |
| 48 | 29784259 | 7.60 | 148 | 85.94 | 91.90 |
| 49 | 28659707 | 7.33 | 165 | 88.98 | 95.24 |
| 50 | 21456851 | 5.67 | 166 | 86.68 | 90.86 |
| 51 | 36269823 | 9.62 | 172 | 95.38 | 97.41 |
| 52 | 62261323 | 12.50 | 285 | 98.74 | 98.94 |
| 53 | 27295877 | 6.94 | 155 | 84.40 | 91.81 |
| 54 | 37373844 | 9.79 | 193 | 95.66 | 97.69 |
| 55 | 75974126 | 15.21 | 359 | 99.07 | 99.34 |
| 56 | 26394718 | 7.04 | 138 | 90.78 | 94.27 |
| 57 | 29682198 | 7.92 | 136 | 93.17 | 95.41 |
| 58 | 28436831 | 7.58 | 143 | 93.37 | 96.44 |
| 59 | 41356663 | 10.86 | 187 | 96.95 | 97.87 |
| 60 | 99703532 | 26.46 | 170 | 98.79 | 98.91 |
| 61 | 23932167 | 6.35 | 174 | 90.23 | 92.79 |
| 62 | 28110933 | 7.50 | 180 | 95.94 | 97.27 |
| 63 | 34190172 | 9.11 | 190 | 96.73 | 98.13 |
| 64 | 26400851 | 6.94 | 199 | 96.64 | 97.79 |
| 65 | 28549775 | 7.52 | 162 | 95.24 | 98.03 |
| 66 | 39729325 | 10.56 | 194 | 97.39 | 98.82 |
| 67 | 23763652 | 6.31 | 186 | 86.41 | 88.94 |
| 68 | 31098605 | 8.29 | 166 | 95.77 | 97.39 |
| 69 | 35833676 | 9.55 | 217 | 97.68 | 98.18 |
| 70 | 49843480 | 9.98 | 379 | 97.94 | 98.96 |
| 71 | 43276882 | 11.34 | 193 | 97.73 | 98.82 |
| 72 | 30513166 | 8.12 | 204 | 97.45 | 98.39 |
| 73 | 48059363 | 9.65 | 273 | 98.69 | 99.03 |
| 74 | 33101770 | 8.82 | 186 | 97.09 | 98.56 |
| 75 | 87991934 | 23.26 | 189 | 98.75 | 98.56 |
| 76 | 23262421 | 6.14 | 163 | 91.64 | 96.01 |
| 77 | 18102982 | 4.78 | 138 | 84.60 | 90.17 |
| 78 | 55845392 | 14.84 | 203 | 97.29 | 98.91 |
| 79 | 18152765 | 4.79 | 124 | 79.25 | 84.72 |
| 80 | 37519560 | 7.53 | 290 | 97.58 | 98.85 |
| 81 | 130949579 | 26.25 | 276 | 99.66 | 99.64 |
| 82 | 47660072 | 12.40 | 190 | 97.30 | 98.58 |
| 83 | 57229899 | 12.13 | 231 | 92.05 | 98.30 |
| 84 | 28838585 | 7.57 | 195 | 92.57 | 93.56 |
| 85 | 38269130 | 8.50 | 198 | 93.28 | 98.58 |

| | | | | | |
|---|---|---|---|---|---|
| 86 | 35582868 | 9.29 | 189 | 95.37 | 97.88 |
| 87 | 38459861 | 10.14 | 168 | 96.93 | 97.88 |
| 88 | 29336797 | 7.69 | 147 | 80.61 | 84.73 |
| 89 | 35671263 | 9.35 | 190 | 97.19 | 98.55 |
| 90 | 135772072 | 27.22 | 281 | 97.47 | 99.14 |
| 91 | 26782782 | 7.08 | 178 | 90.84 | 92.38 |
| 92 | 13667346 | 3.58 | 258 | 83.59 | 86.48 |
| 93 | 34419826 | 9.08 | 176 | 97.36 | 98.44 |
| 94 | 24833902 | 6.44 | 161 | 86.84 | 91.14 |
| 95 | 37113906 | 9.76 | 171 | 93.16 | 96.38 |
| 96 | 35001979 | 9.23 | 171 | 97.50 | 98.37 |
| 97 | 47737567 | 12.73 | 215 | 96.92 | 97.71 |
| 98 | 22765857 | 6.03 | 257 | 94.86 | 95.88 |
| 99 | 108250182 | 28.85 | 173 | 99.13 | 99.38 |
| 100 | 67700116 | 18.05 | 205 | 99.15 | 99.31 |
| 101 | 30323656 | 8.01 | 175 | 96.15 | 97.06 |
| 102 | 18224974 | 4.79 | 168 | 90.16 | 91.42 |
| 103 | 18232002 | 3.98 | 198 | 60.85 | 78.93 |
| 104 | 40963502 | 8.22 | 315 | 92.88 | 98.89 |
| 105 | 101828444 | 26.59 | 182 | 90.69 | 96.08 |
| 106 | 32429653 | 8.48 | 195 | 97.22 | 98.29 |
| 107 | 27792518 | 7.29 | 178 | 82.79 | 85.73 |
| 108 | 150092616 | 31.21 | 170 | 98.16 | 99.18 |
| 109 | 63629812 | 16.79 | 200 | 98.82 | 99.22 |
| 110 | 20892590 | 5.47 | 198 | 87.12 | 91.27 |
| 111 | 65263523 | 17.22 | 192 | 98.31 | 99.31 |
| 112 | 14453939 | 3.84 | 167 | 80.25 | 85.60 |
| 113 | 32373771 | 8.58 | 157 | 93.83 | 96.67 |
| 114 | 99296014 | 19.76 | 223 | 92.41 | 98.34 |
| 115 | 20481128 | 4.11 | 273 | 92.29 | 95.38 |
| 116 | 30640636 | 8.16 | 189 | 96.94 | 98.24 |
| 117 | 31272050 | 8.27 | 183 | 96.55 | 98.14 |
| 118 | 39308552 | 10.44 | 193 | 97.01 | 98.46 |
| 119 | 40099621 | 10.68 | 166 | 96.93 | 98.59 |
| 120 | 40916314 | 10.74 | 213 | 97.66 | 98.72 |
| 121 | 33081747 | 8.67 | 189 | 93.16 | 98.33 |
| 122 | 47346337 | 12.54 | 169 | 97.03 | 98.61 |
| 123 | 58596234 | 15.52 | 200 | 98.06 | 98.90 |
| 124 | 36484537 | 8.27 | 214 | 89.83 | 98.36 |
| 125 | 21837719 | 5.77 | 181 | 92.12 | 95.59 |
| 126 | 50553682 | 13.29 | 192 | 94.27 | 96.22 |
| 127 | 56502574 | 11.35 | 278 | 98.15 | 98.69 |

| | | | | | |
|---|---|---|---|---|---|
| 128 | 45337178 | 11.97 | 196 | 97.91 | 98.89 |
| 129 | 56819652 | 15.03 | 179 | 97.82 | 98.81 |
| 130 | 34315106 | 9.03 | 187 | 93.56 | 95.26 |
| 131 | 24875332 | 4.89 | 199 | 88.63 | 88.45 |
| 132 | 115347450 | 23.13 | 274 | 99.09 | 99.06 |
| 133 | 39141197 | 8.71 | 190 | 72.15 | 88.29 |
| 134 | 35144802 | 9.30 | 169 | 96.83 | 98.35 |
| 135 | 55150684 | 11.07 | 286 | 98.25 | 98.87 |
| 136 | 26893243 | 7.10 | 162 | 95.82 | 97.59 |
| 137 | 31624850 | 8.36 | 164 | 95.53 | 97.54 |
| 138 | 43939999 | 11.46 | 155 | 88.94 | 93.83 |
| 139 | 27085793 | 7.15 | 166 | 95.80 | 97.46 |
| 140 | 61930077 | 16.43 | 164 | 97.96 | 98.75 |
| 141 | 30137263 | 7.98 | 169 | 97.15 | 98.22 |
| 142 | 29439814 | 7.70 | 193 | 84.61 | 86.06 |
| 143 | 31020939 | 8.16 | 189 | 92.54 | 93.05 |
| 144 | 39331854 | 10.31 | 199 | 96.85 | 98.86 |
| 145 | 19352702 | 3.88 | 296 | 92.28 | 94.79 |
| 146 | 32568107 | 8.57 | 182 | 92.51 | 95.78 |
| 147 | 53529738 | 10.75 | 281 | 98.67 | 99.14 |
| 148 | 36252712 | 7.28 | 275 | 92.04 | 98.18 |
| 149 | 22639197 | 5.90 | 192 | 82.98 | 86.06 |
| 150 | 47411849 | 12.51 | 193 | 80.23 | 89.69 |
| 151 | 20854354 | 5.51 | 165 | 94.34 | 96.31 |
| 153 | 63862983 | 13.71 | 182 | 75.03 | 82.98 |
| 154 | 26702546 | 5.36 | 311 | 84.10 | 97.35 |
| 155 | 27643954 | 7.30 | 157 | 93.48 | 96.42 |
| 156 | 105291759 | 27.85 | 170 | 98.59 | 99.30 |
| 157 | 33463390 | 8.77 | 197 | 96.49 | 98.20 |
| 158 | 35287979 | 9.25 | 187 | 97.45 | 98.78 |
| 159 | 40340788 | 10.68 | 179 | 97.56 | 98.23 |
| 160 | 33340181 | 8.78 | 185 | 94.25 | 96.93 |
| 161 | 43348624 | 11.26 | 202 | 90.68 | 94.79 |
| 162 | 69416120 | 18.56 | 171 | 94.38 | 95.96 |
| 163 | 18894011 | 5.01 | 202 | 92.41 | 94.58 |
| 164 | 68030315 | 13.66 | 271 | 98.74 | 99.26 |
| 165 | 24573464 | 6.43 | 239 | 96.32 | 97.82 |
| 166 | 73402701 | 19.45 | 187 | 98.98 | 98.99 |
| 167 | 51338378 | 13.59 | 203 | 96.66 | 97.87 |
| 168 | 57539178 | 15.23 | 238 | 98.44 | 99.19 |
| 169 | 88933904 | 17.82 | 278 | 98.91 | 98.80 |
| 170 | 64007878 | 12.85 | 290 | 98.70 | 99.04 |

| | | | | | |
|---|---|---|---|---|---|
| 172 | 22520554 | 5.96 | 194 | 84.55 | 89.00 |
| 173 | 48172756 | 12.67 | 189 | 94.92 | 96.45 |
| 174 | 70466470 | 18.39 | 184 | 94.88 | 96.68 |
| 175 | 26465110 | 6.95 | 183 | 80.14 | 82.88 |
| 176 | 46950500 | 12.23 | 186 | 87.70 | 92.72 |
| 177 | 78760465 | 20.95 | 223 | 97.07 | 97.93 |
| 178 | 19785124 | 5.22 | 205 | 94.68 | 96.01 |
| 179 | 19188073 | 5.12 | 210 | 95.51 | 96.77 |
| 180 | 156030354 | 31.28 | 288 | 99.48 | 99.16 |
| 181 | 42745264 | 11.06 | 194 | 86.71 | 90.54 |
| 182 | 40862025 | 10.58 | 192 | 80.51 | 83.72 |
| 183 | 26448467 | 6.99 | 194 | 83.57 | 83.18 |
| 184 | 32929204 | 6.61 | 279 | 96.89 | 98.11 |
| 185 | 29689283 | 7.85 | 172 | 80.46 | 83.28 |
| 186 | 18390542 | 4.81 | 191 | 81.75 | 86.37 |
| 187 | 43806955 | 11.56 | 261 | 97.96 | 98.54 |
| 188 | 29004974 | 5.82 | 290 | 95.39 | 97.70 |
| 189 | 17735883 | 4.69 | 182 | 89.27 | 91.52 |
| 190 | 31285034 | 6.27 | 381 | 96.16 | 98.02 |
| 191 | 16746728 | 4.41 | 198 | 93.88 | 94.89 |
| 192 | 43191518 | 8.67 | 279 | 97.93 | 98.99 |
| 193 | 20183774 | 5.21 | 210 | 86.10 | 92.70 |
| 194 | 131726771 | 26.41 | 266 | 98.76 | 98.95 |
| 195 | 34338281 | 8.98 | 177 | 86.51 | 88.90 |
| 196 | 44872044 | 9.01 | 282 | 97.29 | 98.93 |
| 197 | 50478900 | 13.32 | 183 | 97.20 | 98.05 |
| 198 | 34917371 | 9.24 | 181 | 97.09 | 98.35 |
| 199 | 54282739 | 14.38 | 188 | 97.20 | 98.15 |
| 200 | 22975778 | 6.05 | 184 | 79.26 | 81.08 |
| 201 | 39087788 | 7.85 | 278 | 97.55 | 98.59 |
| 203 | 34036807 | 9.02 | 189 | 90.72 | 94.58 |
| 204 | 19091866 | 5.01 | 197 | 93.80 | 95.31 |
| 205 | 37126800 | 9.86 | 185 | 98.01 | 98.61 |
| 206 | 44748927 | 11.80 | 269 | 98.47 | 98.97 |
| 207 | 45718101 | 12.15 | 193 | 97.72 | 98.91 |
| 208 | 13896308 | 3.50 | 256 | 82.73 | 88.20 |
| 209 | 69461358 | 13.95 | 308 | 98.65 | 98.81 |
| 210 | 24036211 | 6.32 | 204 | 96.75 | 97.34 |
| 211 | 35380498 | 9.31 | 189 | 92.23 | 94.29 |
| 212 | 93023734 | 19.73 | 218 | 97.10 | 99.12 |
| 213 | 23203247 | 6.13 | 171 | 93.58 | 95.16 |
| 214 | 21635749 | 5.72 | 199 | 93.28 | 95.50 |

| | | | | | |
|---|---|---|---|---|---|
| 215 | 23810036 | 6.27 | 199 | 91.94 | 95.10 |
| 216 | 97717902 | 26.01 | 206 | 95.70 | 98.00 |
| 217 | 24242436 | 6.26 | 172 | 90.39 | 94.33 |
| 218 | 37325947 | 9.87 | 169 | 90.65 | 92.65 |
| 219 | 39216524 | 10.28 | 181 | 95.11 | 97.66 |
| 220 | 35666562 | 7.16 | 330 | 79.94 | 98.40 |
| 221 | 39362569 | 10.40 | 245 | 97.46 | 98.56 |
| 222 | 58183878 | 12.90 | 184 | 97.32 | 98.63 |
| 223 | 116416542 | 23.38 | 290 | 99.24 | 99.17 |
| 224 | 34256204 | 8.92 | 187 | 82.54 | 85.35 |
| 225 | 99788611 | 26.23 | 221 | 90.43 | 92.72 |
| 226 | 33673524 | 8.85 | 199 | 97.40 | 98.00 |
| 227 | 26964409 | 7.17 | 187 | 96.99 | 97.87 |
| 228 | 69982302 | 14.05 | 291 | 98.37 | 98.76 |
| 229 | 32073645 | 8.47 | 179 | 95.77 | 98.46 |
| 230 | 62384302 | 12.52 | 286 | 98.64 | 98.94 |
| 231 | 80771086 | 16.21 | 295 | 98.78 | 98.97 |
| 232 | 81102867 | 16.29 | 267 | 98.25 | 98.82 |
| 233 | 83562405 | 22.08 | 184 | 94.46 | 96.28 |
| 234 | 73428554 | 19.47 | 202 | 99.06 | 99.03 |
| 235 | 31506873 | 8.33 | 193 | 84.76 | 86.43 |
| 236 | 90866634 | 19.34 | 230 | 98.13 | 98.97 |
| 237 | 93911578 | 24.92 | 162 | 99.02 | 99.29 |
| 238 | 43254548 | 8.68 | 285 | 96.64 | 98.46 |
| 239 | 46700347 | 12.36 | 193 | 96.43 | 97.50 |
| 240 | 113833688 | 22.80 | 282 | 99.14 | 99.20 |
| 241 | 44190816 | 11.66 | 174 | 98.48 | 98.60 |
| 242 | 77344062 | 15.53 | 286 | 98.77 | 99.18 |
| 244 | 79107401 | 15.88 | 279 | 98.43 | 99.29 |
| 245 | 56549727 | 14.96 | 177 | 96.27 | 98.62 |
| 246 | 85056700 | 22.52 | 186 | 98.05 | 98.99 |
| 247 | 37941521 | 10.03 | 174 | 96.52 | 98.39 |
| 248 | 36349018 | 7.29 | 291 | 97.45 | 98.31 |
| 249 | 23975384 | 6.36 | 151 | 93.24 | 96.80 |
| 250 | 31455139 | 6.31 | 318 | 96.40 | 98.19 |
| 251 | 28102454 | 7.45 | 181 | 97.42 | 98.44 |
| 252 | 122206896 | 32.37 | 183 | 98.03 | 99.26 |
| 253 | 95174019 | 25.07 | 187 | 92.53 | 98.33 |
| 254 | 97603758 | 25.75 | 155 | 98.03 | 99.02 |
| 255 | 31158156 | 8.26 | 174 | 97.42 | 98.31 |
| 256 | 41501020 | 10.57 | 215 | 95.64 | 98.28 |
| 257 | 16892124 | 4.47 | 156 | 85.22 | 89.97 |

| | | | | | |
|---|---|---|---|---|---|
| 258 | 16653675 | 4.42 | 177 | 80.85 | 84.17 |
| 259 | 65572096 | 17.40 | 180 | 98.39 | 98.86 |
| 260 | 45829502 | 12.20 | 173 | 95.24 | 97.57 |
| 261 | 78556264 | 16.69 | 193 | 93.91 | 97.96 |
| 262 | 20682120 | 5.46 | 199 | 88.74 | 93.63 |
| 263 | 24724949 | 6.53 | 189 | 82.24 | 83.32 |
| 264 | 16894657 | 4.47 | 188 | 84.01 | 89.05 |
| 265 | 25682293 | 6.48 | 234 | 93.67 | 97.12 |
| 266 | 52930674 | 14.06 | 166 | 98.72 | 99.16 |
| 267 | 22090084 | 5.83 | 179 | 82.05 | 84.65 |
| 268 | 25680448 | 10.18 | 206 | 94.56 | 98.63 |
| 269 | 57990233 | 15.23 | 208 | 86.65 | 88.98 |
| 270 | 34207974 | 9.06 | 156 | 95.99 | 98.29 |
| 271 | 28432635 | 7.54 | 180 | 89.70 | 94.75 |
| 272 | 50939281 | 11.08 | 208 | 91.96 | 97.93 |
| 273 | 106479958 | 28.15 | 163 | 96.83 | 99.00 |
| 274 | 19922818 | 5.22 | 208 | 94.28 | 96.23 |
| 275 | 17847361 | 4.72 | 166 | 81.90 | 87.97 |
| 276 | 24062940 | 6.14 | 155 | 80.86 | 84.70 |
| 277 | 150041504 | 39.31 | 191 | 98.41 | 98.59 |
| 278 | 41017018 | 10.87 | 211 | 93.67 | 98.47 |
| 279 | 99205070 | 21.70 | 211 | 91.17 | 96.77 |
| 280 | 92958817 | 18.65 | 285 | 98.85 | 98.86 |

RIL IDs are assigned using plant numbers (1 to 280) of the F2 generation. Five lines have been lost during the process of breeding and three lines have been removed because they are duplicates of other RILs. The lines that has been lost during breeding are 27, 32, 41, 171, and 202. The lines that are duplicates and have been removed are 6 (duplicate of 5), 152 (duplicate of 145), and 243 (duplicate of 278).

Supplemenary Data 2. List of *mPing* and *Ping* copy numbers in the RILs

| RIL ID | Total number of *mPing* | Number of shared parental *mPing* | Number of other shared *mPing* | Number of unique *mPing* | Number of unique homozygous *mPing* | Number of unique heterozgyous *mPing* | *Ping* copy number | *Pong* copy number | *Ping* code |
|---|---|---|---|---|---|---|---|---|---|
| 1 | 253 | 176 | 49 | 28 | 19 | 9 | 6 | 1 | CDEFGH |
| 2 | 163 | 131 | 20 | 12 | 10 | 2 | 3 | 4 | CDH |
| 3 | 303 | 206 | 50 | 47 | 43 | 4 | 7 | 3 | *ABCEFG |
| 4 | 234 | 167 | 41 | 26 | 23 | 3 | 5 | 4 | CDEFG |
| 5 | 211 | 156 | 32 | 23 | 18 | 5 | 3 | 4 | FGH |
| 7 | 247 | 203 | 35 | 9 | 9 | 0 | 3 | 3 | FGH |
| 8 | 274 | 190 | 34 | 50 | 40 | 10 | 4 | 3 | ABDF |
| 9 | 214 | 160 | 29 | 25 | 16 | 9 | 4 | 3 | CEFH |
| 10 | 344 | 238 | 56 | 50 | 39 | 11 | 5 | 4 | ABCDF |
| 11 | 435 | 257 | 66 | 112 | 97 | 15 | 7 | 3 | ABDEFGH |
| 12 | 264 | 231 | 32 | 1 | 1 | 0 | 1 | 4 | G |
| 13 | 464 | 223 | 66 | 175 | 105 | 70 | 4 | 4 | ABGH |
| 14 | 325 | 198 | 50 | 77 | 48 | 29 | 6 | 3 | *CEFGH |
| 15 | 375 | 210 | 56 | 109 | 75 | 34 | 5 | 4 | DEFGH |
| 16 | 283 | 247 | 35 | 1 | 0 | 1 | 2 | 3 | EF |
| 17 | 229 | 183 | 26 | 20 | 14 | 6 | 2 | 4 | DG |
| 18 | 260 | 214 | 30 | 16 | 13 | 3 | 3 | 4 | DFG |
| 19 | 236 | 175 | 40 | 21 | 18 | 3 | 2 | 4 | EG |
| 20 | 226 | 176 | 34 | 16 | 15 | 1 | 2 | 4 | BD |
| 21 | 309 | 207 | 45 | 57 | 38 | 19 | 7 | 3 | ABCEFGH |
| 22 | 309 | 229 | 52 | 28 | 21 | 7 | 3 | 3 | ABC |
| 23 | 292 | 219 | 38 | 35 | 28 | 7 | 5 | 3 | ABDEF |
| 24 | 414 | 248 | 60 | 106 | 73 | 33 | 6 | 4 | ABCDFH |
| 25 | 261 | 227 | 33 | 1 | 1 | 0 | 1 | 3 | D |
| 26 | 371 | 256 | 52 | 63 | 42 | 21 | 5 | 2 | *DEFG |

| | | | | | | | | | |
|---|---|---|---|---|---|---|---|---|---|
| 28 | 217 | 169 | 32 | 16 | 15 | 1 | 2 | 4 | CH |
| 29 | 286 | 199 | 43 | 44 | 22 | 22 | 3 | 4 | CDH |
| 30 | 306 | 203 | 45 | 58 | 55 | 3 | 6 | 3 | ABEFGH |
| 31 | 263 | 201 | 39 | 23 | 21 | 2 | 5 | 3 | CEFGH |
| 33 | 273 | 212 | 47 | 14 | 11 | 3 | 2 | 3 | CD |
| 34 | 473 | 279 | 84 | 110 | 82 | 28 | 7 | 4 | ABCDEFG |
| 35 | 258 | 209 | 41 | 8 | 8 | 0 | 2 | 3 | FH |
| 36 | 311 | 254 | 45 | 12 | 8 | 4 | 2 | 3 | AB |
| 37 | 230 | 176 | 29 | 25 | 16 | 9 | 3 | 3 | ABH |
| 38 | 281 | 222 | 36 | 23 | 19 | 4 | 6 | 3 | CDEFGH |
| 39 | 234 | 190 | 42 | 2 | 2 | 0 | 0 | 3 | No *Ping* |
| 40 | 291 | 238 | 48 | 5 | 5 | 0 | 1 | 4 | D |
| 42 | 307 | 246 | 45 | 16 | 12 | 4 | 3 | 4 | DEF |
| 43 | 294 | 234 | 38 | 22 | 21 | 1 | 1 | 4 | C |
| 44 | 354 | 210 | 59 | 85 | 76 | 9 | 6 | 3 | ABCEFG |
| 45 | 237 | 199 | 34 | 4 | 4 | 0 | 1 | 3 | E |
| 46 | 385 | 212 | 54 | 119 | 73 | 46 | 7 | 3 | *ABEFGH |
| 47 | 252 | 224 | 25 | 3 | 3 | 0 | 1 | 4 | G |
| 48 | 288 | 173 | 48 | 67 | 54 | 13 | 4 | 1 | ABFG |
| 49 | 338 | 209 | 47 | 82 | 63 | 19 | 4 | 3 | ABEH |
| 50 | 295 | 212 | 42 | 41 | 39 | 2 | 6 | 3 | ABCDGH |
| 51 | 377 | 222 | 63 | 92 | 67 | 25 | 6 | 3 | *ABDEF |
| 52 | 338 | 273 | 42 | 23 | 16 | 7 | 3 | 4 | CDE |
| 53 | 249 | 179 | 41 | 29 | 28 | 1 | 3 | 4 | ABD |
| 54 | 423 | 255 | 57 | 111 | 71 | 40 | 6 | 4 | ABCDGH |
| 55 | 278 | 205 | 39 | 34 | 21 | 13 | 4 | 3 | DFGH |
| 56 | 375 | 201 | 57 | 117 | 78 | 39 | 6 | 4 | ABDEFG |
| 57 | 415 | 256 | 74 | 85 | 54 | 31 | 6 | 3 | CDEFGH |
| 58 | 309 | 216 | 51 | 42 | 29 | 13 | 5 | 3 | CDEFG |
| 59 | 371 | 284 | 55 | 32 | 21 | 11 | 3 | 4 | CDG |

| | | | | | | | | | |
|---|---|---|---|---|---|---|---|---|---|
| 60 | 241 | 202 | 35 | 4 | 4 | 0 | 1 | 3 | A |
| 61 | 288 | 188 | 42 | 58 | 56 | 2 | 5 | 2 | CDEFG |
| 62 | 392 | 233 | 49 | 110 | 76 | 34 | 5 | 4 | ACEFG |
| 63 | 388 | 253 | 60 | 75 | 53 | 22 | 3 | 4 | ABC |
| 64 | 346 | 218 | 56 | 72 | 53 | 19 | 7 | 4 | ABCEFGH |
| 65 | 352 | 223 | 49 | 80 | 62 | 18 | 6 | 4 | *ABDEF |
| 66 | 467 | 276 | 71 | 120 | 87 | 33 | 6 | 3 | ABDEFG |
| 67 | 200 | 166 | 29 | 5 | 2 | 3 | 2 | 3 | CG |
| 68 | 349 | 217 | 54 | 78 | 56 | 22 | 4 | 3 | AEFG |
| 69 | 423 | 257 | 68 | 98 | 79 | 19 | 7 | 3 | ABCDEFG |
| 70 | 368 | 270 | 56 | 42 | 27 | 15 | 4 | 4 | ABDG |
| 71 | 358 | 229 | 61 | 68 | 44 | 24 | 4 | 4 | ABCE |
| 72 | 407 | 241 | 69 | 97 | 68 | 29 | 6 | 4 | ABCEFG |
| 73 | 280 | 194 | 48 | 38 | 30 | 8 | 5 | 4 | ABCFH |
| 74 | 307 | 224 | 55 | 28 | 23 | 5 | 3 | 4 | ADH |
| 75 | 321 | 274 | 47 | 0 | 0 | 0 | 1 | 3 | D |
| 76 | 283 | 211 | 45 | 27 | 19 | 8 | 4 | 3 | BCGH |
| 77 | 315 | 185 | 56 | 74 | 52 | 22 | 4 | 1 | ABCD |
| 78 | 411 | 215 | 74 | 122 | 73 | 49 | 4 | 3 | ABCH |
| 79 | 166 | 137 | 21 | 8 | 7 | 1 | 2 | 1 | DH |
| 80 | 312 | 208 | 51 | 53 | 41 | 12 | 2 | 4 | BC |
| 81 | 235 | 189 | 32 | 14 | 4 | 10 | 2 | 4 | CH |
| 82 | 259 | 199 | 37 | 23 | 18 | 5 | 3 | 4 | EFG |
| 83 | 399 | 237 | 66 | 96 | 67 | 29 | 5 | 4 | ABCEF |
| 84 | 305 | 245 | 45 | 15 | 11 | 4 | 4 | 3 | CEFH |
| 85 | 217 | 155 | 29 | 33 | 29 | 4 | 3 | 3 | EFG |
| 86 | 353 | 257 | 54 | 42 | 26 | 16 | 5 | 4 | BCEFG |
| 87 | 375 | 244 | 52 | 79 | 58 | 21 | 4 | 4 | ABCG |
| 88 | 330 | 229 | 48 | 53 | 51 | 2 | 5 | 3 | ABDEH |
| 89 | 290 | 183 | 42 | 65 | 47 | 18 | 4 | 3 | EFGH |

| | | | | | | | | | |
|---|---|---|---|---|---|---|---|---|---|
| 90 | 395 | 247 | 57 | 91 | 58 | 33 | 6 | 3 | CDEFGH |
| 91 | 332 | 265 | 43 | 24 | 20 | 4 | 3 | 4 | CGH |
| 92 | 341 | 187 | 64 | 90 | 61 | 29 | 5 | 4 | ABEFG |
| 93 | 288 | 214 | 44 | 30 | 21 | 9 | 4 | 4 | *BGH |
| 94 | 341 | 218 | 51 | 72 | 63 | 9 | 5 | 4 | ABEFG |
| 95 | 341 | 258 | 52 | 31 | 31 | 0 | 5 | 4 | ABFGH |
| 96 | 380 | 214 | 54 | 112 | 89 | 23 | 7 | 3 | ABCDEFH |
| 97 | 373 | 275 | 59 | 39 | 28 | 11 | 3 | 4 | ABG |
| 98 | 382 | 265 | 61 | 56 | 47 | 9 | 3 | 3 | CDH |
| 99 | 303 | 259 | 41 | 3 | 2 | 1 | 1 | 4 | D |
| 100 | 215 | 192 | 22 | 1 | 0 | 1 | 1 | 3 | D |
| 101 | 262 | 174 | 45 | 43 | 33 | 10 | 5 | 3 | CDFGH |
| 102 | 165 | 110 | 31 | 24 | 21 | 3 | 2 | 4 | CH |
| 103 | 216 | 155 | 39 | 22 | 20 | 2 | 3 | 2 | ABH |
| 104 | 317 | 212 | 54 | 51 | 44 | 7 | 4 | 4 | *ABH |
| 105 | 356 | 202 | 52 | 102 | 83 | 19 | 6 | 3 | ABEFGH |
| 106 | 283 | 239 | 41 | 3 | 2 | 1 | 0 | 4 | No *Ping* |
| 107 | 260 | 169 | 38 | 53 | 44 | 9 | 4 | 1 | ABDH |
| 108 | 298 | 231 | 37 | 30 | 26 | 4 | 3 | 3 | CDH |
| 109 | 278 | 203 | 42 | 33 | 23 | 10 | 3 | 4 | CDH |
| 110 | 366 | 288 | 55 | 23 | 20 | 3 | 3 | 3 | ABC |
| 111 | 305 | 204 | 57 | 44 | 34 | 10 | 3 | 4 | EGH |
| 112 | 254 | 155 | 48 | 51 | 34 | 17 | 5 | 3 | ABEFH |
| 113 | 345 | 227 | 64 | 54 | 38 | 16 | 3 | 4 | ABC |
| 114 | 245 | 190 | 31 | 24 | 23 | 1 | 2 | 4 | EF |
| 115 | 233 | 176 | 27 | 30 | 28 | 2 | 3 | 3 | BDG |
| 116 | 281 | 201 | 41 | 39 | 24 | 15 | 3 | 3 | CDH |
| 117 | 384 | 197 | 71 | 116 | 82 | 34 | 5 | 4 | ABEFG |
| 118 | 375 | 223 | 60 | 92 | 70 | 22 | 6 | 3 | CDEFGH |
| 119 | 357 | 212 | 59 | 86 | 42 | 44 | 4 | 3 | ABCH |

| | | | | | | | | | |
|---|---|---|---|---|---|---|---|---|---|
| 120 | 485 | 235 | 72 | 178 | 96 | 82 | 6 | 3 | ABCEFG |
| 121 | 359 | 188 | 50 | 121 | 86 | 35 | 5 | 4 | ABCGH |
| 122 | 373 | 213 | 65 | 95 | 68 | 27 | 5 | 3 | ACEFG |
| 123 | 261 | 202 | 37 | 22 | 13 | 9 | 2 | 4 | AB |
| 124 | 270 | 191 | 58 | 21 | 15 | 6 | 6 | 4 | ABEFGH |
| 125 | 274 | 211 | 42 | 21 | 18 | 3 | 3 | 3 | BCD |
| 126 | 354 | 231 | 51 | 72 | 53 | 19 | 5 | 3 | ABCGH |
| 127 | 406 | 252 | 56 | 98 | 69 | 29 | 6 | 3 | ABCDEG |
| 128 | 358 | 241 | 63 | 54 | 42 | 12 | 3 | 4 | ABC |
| 129 | 240 | 212 | 26 | 2 | 2 | 0 | 0 | 4 | No *Ping* |
| 130 | 331 | 220 | 55 | 56 | 40 | 16 | 5 | 3 | ABCGH |
| 131 | 328 | 239 | 48 | 41 | 29 | 12 | 5 | 3 | ABDFG |
| 132 | 368 | 267 | 54 | 47 | 19 | 28 | 3 | 3 | CDF |
| 133 | 404 | 263 | 72 | 69 | 65 | 4 | 7 | 3 | ABCDEFG |
| 134 | 427 | 257 | 59 | 111 | 76 | 35 | 6 | 3 | ABCDEH |
| 135 | 386 | 232 | 63 | 91 | 51 | 40 | 6 | 3 | ABCEFG |
| 136 | 466 | 271 | 75 | 120 | 90 | 30 | 5 | 4 | ABDEF |
| 137 | 416 | 262 | 75 | 79 | 59 | 20 | 3 | 3 | ABH |
| 138 | 287 | 208 | 47 | 32 | 30 | 2 | 5 | 4 | ABCGH |
| 139 | 239 | 195 | 32 | 12 | 11 | 1 | 2 | 4 | CG |
| 140 | 275 | 239 | 27 | 9 | 9 | 0 | 1 | 3 | H |
| 141 | 289 | 222 | 31 | 36 | 28 | 8 | 2 | 4 | DG |
| 142 | 380 | 216 | 62 | 102 | 72 | 30 | 6 | 2 | ABDEFH |
| 143 | 373 | 299 | 48 | 26 | 21 | 5 | 5 | 3 | CDEFG |
| 144 | 462 | 201 | 65 | 196 | 138 | 58 | 6 | 4 | ABDEFG |
| 145 | 204 | 147 | 26 | 31 | 27 | 4 | 4 | 3 | ABEH |
| 146 | 415 | 239 | 70 | 106 | 83 | 23 | 4 | 4 | ABDE |
| 147 | 291 | 214 | 37 | 40 | 19 | 21 | 3 | 3 | DEG |
| 148 | 374 | 242 | 55 | 77 | 49 | 28 | 3 | 4 | CFG |
| 149 | 303 | 182 | 54 | 67 | 52 | 15 | 3 | 4 | *GH |

| | | | | | | | | | |
|---|---|---|---|---|---|---|---|---|---|
| 150 | 358 | 231 | 77 | 50 | 40 | 10 | 3 | 2 | ABH |
| 151 | 339 | 229 | 35 | 75 | 45 | 30 | 4 | 3 | ABCG |
| 153 | 354 | 219 | 54 | 81 | 67 | 14 | 6 | 4 | ABDFGH |
| 154 | 338 | 250 | 64 | 24 | 9 | 15 | 3 | 3 | DGH |
| 155 | 376 | 269 | 40 | 67 | 49 | 18 | 7 | 3 | ABDEFGH |
| 156 | 261 | 228 | 25 | 8 | 8 | 0 | 1 | 3 | H |
| 157 | 382 | 276 | 60 | 46 | 41 | 5 | 4 | 3 | CFGH |
| 158 | 374 | 224 | 62 | 88 | 73 | 15 | 3 | 3 | ACH |
| 159 | 485 | 251 | 93 | 141 | 95 | 46 | 5 | 4 | ABCDG |
| 160 | 378 | 285 | 42 | 51 | 41 | 10 | 6 | 3 | ABCEFH |
| 161 | 258 | 204 | 32 | 22 | 20 | 2 | 2 | 3 | CD |
| 162 | 319 | 238 | 38 | 43 | 35 | 8 | 3 | 4 | EFG |
| 163 | 269 | 202 | 32 | 35 | 32 | 3 | 6 | 4 | ABCEFG |
| 164 | 306 | 216 | 40 | 50 | 32 | 18 | 4 | 4 | AEFG |
| 165 | 289 | 227 | 36 | 26 | 21 | 5 | 3 | 4 | EFG |
| 166 | 237 | 213 | 24 | 0 | 0 | 0 | 1 | 3 | H |
| 167 | 455 | 280 | 57 | 118 | 94 | 24 | 5 | 4 | ABDFG |
| 168 | 266 | 206 | 39 | 21 | 9 | 12 | 3 | 4 | CGH |
| 169 | 309 | 223 | 40 | 46 | 29 | 17 | 4 | 4 | ACDH |
| 170 | 351 | 262 | 44 | 45 | 33 | 12 | 5 | 3 | CDEFG |
| 172 | 307 | 236 | 38 | 33 | 24 | 9 | 4 | 4 | DEFG |
| 173 | 283 | 222 | 32 | 29 | 24 | 5 | 2 | 3 | DH |
| 174 | 233 | 138 | 37 | 58 | 43 | 15 | 7 | 4 | *ABDEFH |
| 175 | 432 | 269 | 59 | 104 | 81 | 23 | 7 | 1 | *ABCDGH |
| 176 | 448 | 290 | 59 | 99 | 79 | 20 | 4 | 3 | ABCE |
| 177 | 270 | 196 | 36 | 38 | 22 | 16 | 4 | 4 | CEFG |
| 178 | 336 | 228 | 49 | 59 | 42 | 17 | 5 | 3 | CDEFG |
| 179 | 178 | 147 | 29 | 2 | 2 | 0 | 1 | 4 | B |
| 180 | 368 | 271 | 43 | 54 | 33 | 21 | 4 | 3 | ABCH |
| 181 | 335 | 227 | 43 | 65 | 53 | 12 | 3 | 3 | ABH |

| | | | | | | | | | |
|---|---|---|---|---|---|---|---|---|---|
| 182 | 294 | 220 | 35 | 39 | 33 | 6 | 3 | 3 | EGH |
| 183 | 281 | 197 | 35 | 49 | 40 | 9 | 4 | 2 | ABFH |
| 184 | 309 | 200 | 38 | 71 | 55 | 16 | 5 | 3 | ABCGH |
| 185 | 321 | 192 | 50 | 79 | 39 | 40 | 6 | 4 | ABDFGH |
| 186 | 316 | 215 | 45 | 56 | 45 | 11 | 6 | 2 | ABCFGH |
| 187 | 335 | 216 | 46 | 73 | 40 | 33 | 3 | 4 | BCH |
| 188 | 283 | 236 | 33 | 14 | 9 | 5 | 3 | 3 | DGH |
| 189 | 309 | 223 | 38 | 48 | 38 | 10 | 8 | 3 | *ABCEFGH |
| 190 | 312 | 231 | 35 | 46 | 30 | 16 | 4 | 4 | ABDH |
| 191 | 303 | 234 | 39 | 30 | 23 | 7 | 5 | 3 | CEFGH |
| 192 | 341 | 250 | 33 | 58 | 40 | 18 | 4 | 3 | ABCE |
| 193 | 262 | 224 | 33 | 5 | 5 | 0 | 2 | 3 | CF |
| 194 | 417 | 272 | 51 | 94 | 66 | 28 | 6 | 4 | ACDEFG |
| 195 | 261 | 209 | 32 | 20 | 15 | 5 | 2 | 3 | CG |
| 196 | 392 | 234 | 56 | 102 | 65 | 37 | 5 | 3 | ABEFG |
| 197 | 359 | 228 | 50 | 81 | 54 | 27 | 6 | 3 | ACDEFG |
| 198 | 426 | 291 | 58 | 77 | 63 | 14 | 7 | 4 | ABCDEFG |
| 199 | 227 | 186 | 24 | 17 | 12 | 5 | 2 | 3 | EG |
| 200 | 296 | 230 | 36 | 30 | 25 | 5 | 4 | 3 | BDEF |
| 201 | 421 | 249 | 60 | 112 | 81 | 31 | 6 | 3 | ABCEFG |
| 203 | 315 | 223 | 39 | 53 | 41 | 12 | 5 | 4 | BDEFG |
| 204 | 325 | 271 | 38 | 16 | 11 | 5 | 3 | 3 | CDF |
| 205 | 343 | 237 | 44 | 62 | 39 | 23 | 5 | 4 | DEFGH |
| 206 | 232 | 194 | 24 | 14 | 14 | 0 | 2 | 4 | CF |
| 207 | 498 | 284 | 72 | 142 | 86 | 56 | 7 | 3 | ABDEFGH |
| 208 | 163 | 117 | 23 | 23 | 19 | 4 | 3 | 2 | BDH |
| 209 | 448 | 280 | 50 | 118 | 72 | 46 | 4 | 3 | ABFG |
| 210 | 189 | 159 | 23 | 7 | 5 | 2 | 2 | 4 | DH |
| 211 | 360 | 210 | 64 | 86 | 73 | 13 | 5 | 4 | *ABDH |
| 212 | 349 | 229 | 43 | 77 | 59 | 18 | 3 | 4 | BDH |

| | | | | | | | | | |
|---|---|---|---|---|---|---|---|---|---|
| 213 | 281 | 213 | 28 | 40 | 30 | 10 | 4 | 3 | ABCD |
| 214 | 312 | 211 | 44 | 57 | 49 | 8 | 4 | 4 | ABEF |
| 215 | 299 | 232 | 35 | 32 | 24 | 8 | 4 | 4 | CDGH |
| 216 | 439 | 241 | 102 | 96 | 79 | 17 | 6 | 3 | ABCDGH |
| 217 | 372 | 241 | 40 | 91 | 61 | 30 | 5 | 3 | ABCEG |
| 218 | 371 | 242 | 54 | 75 | 62 | 13 | 4 | 3 | ABCH |
| 219 | 298 | 196 | 41 | 61 | 50 | 11 | 4 | 4 | ABEF |
| 220 | 440 | 261 | 110 | 69 | 26 | 43 | 7 | 4 | ABCDEGH |
| 221 | 429 | 235 | 64 | 130 | 70 | 60 | 7 | 3 | *ABEFGH |
| 222 | 346 | 209 | 55 | 82 | 37 | 45 | 5 | 4 | ABFGH |
| 223 | 360 | 195 | 56 | 109 | 49 | 60 | 6 | 3 | CDEFGH |
| 224 | 318 | 209 | 40 | 69 | 57 | 12 | 3 | 4 | DGH |
| 225 | 317 | 264 | 40 | 13 | 13 | 0 | 2 | 3 | CD |
| 226 | 312 | 254 | 34 | 24 | 9 | 15 | 3 | 3 | ABD |
| 227 | 216 | 178 | 26 | 12 | 10 | 2 | 2 | 4 | DE |
| 228 | 357 | 262 | 49 | 46 | 31 | 15 | 6 | 4 | BDEFGH |
| 229 | 392 | 276 | 53 | 63 | 41 | 22 | 4 | 4 | ABCG |
| 230 | 239 | 206 | 31 | 2 | 1 | 1 | 1 | 3 | C |
| 231 | 189 | 162 | 24 | 3 | 3 | 0 | 1 | 4 | H |
| 232 | 417 | 278 | 51 | 88 | 64 | 24 | 4 | 3 | ABEF |
| 233 | 374 | 245 | 42 | 87 | 68 | 19 | 3 | 4 | ABE |
| 234 | 284 | 241 | 33 | 10 | 4 | 6 | 1 | 4 | C |
| 235 | 359 | 257 | 49 | 53 | 42 | 11 | 5 | 3 | ABEFG |
| 236 | 332 | 225 | 51 | 56 | 47 | 9 | 5 | 3 | ABEFG |
| 237 | 216 | 185 | 28 | 3 | 2 | 1 | 1 | 3 | H |
| 238 | 302 | 256 | 34 | 12 | 12 | 0 | 1 | 4 | G |
| 239 | 299 | 174 | 50 | 75 | 52 | 23 | 7 | 3 | ABDEFGH |
| 240 | 297 | 225 | 38 | 34 | 29 | 5 | 4 | 4 | CDFG |
| 241 | 271 | 170 | 36 | 65 | 41 | 24 | 5 | 3 | CEFGH |
| 242 | 436 | 237 | 65 | 134 | 69 | 65 | 4 | 4 | ABCH |

| | | | | | | | | | |
|---|---|---|---|---|---|---|---|---|---|
| 244 | 310 | 236 | 37 | 37 | 29 | 8 | 2 | 3 | AB |
| 245 | 379 | 264 | 42 | 73 | 42 | 31 | 4 | 4 | CEFH |
| 246 | 285 | 220 | 31 | 34 | 26 | 8 | 4 | 4 | EFGH |
| 247 | 381 | 247 | 52 | 82 | 59 | 23 | 5 | 4 | ABDEF |
| 248 | 350 | 214 | 59 | 77 | 57 | 20 | 6 | 4 | *ABFGH |
| 249 | 312 | 246 | 42 | 24 | 21 | 3 | 2 | 3 | DE |
| 250 | 345 | 205 | 48 | 92 | 67 | 25 | 4 | 4 | ABDH |
| 251 | 257 | 216 | 28 | 13 | 12 | 1 | 2 | 3 | EF |
| 252 | 347 | 246 | 46 | 55 | 36 | 19 | 5 | 3 | CDFGH |
| 253 | 351 | 210 | 65 | 76 | 55 | 21 | 5 | 4 | ABCEH |
| 254 | 261 | 226 | 30 | 5 | 5 | 0 | 1 | 4 | D |
| 255 | 324 | 212 | 53 | 59 | 51 | 8 | 6 | 4 | ABCDEF |
| 256 | 284 | 220 | 32 | 32 | 26 | 6 | 2 | 4 | DH |
| 257 | 276 | 191 | 41 | 44 | 35 | 9 | 5 | 1 | ABEFG |
| 258 | 252 | 201 | 26 | 25 | 23 | 2 | 4 | 3 | ABDG |
| 259 | 217 | 184 | 32 | 1 | 0 | 1 | 1 | 3 | H |
| 260 | 339 | 230 | 52 | 57 | 47 | 10 | 6 | 3 | ABCEFH |
| 261 | 409 | 267 | 64 | 78 | 62 | 16 | 7 | 4 | ABCDFGH |
| 262 | 311 | 239 | 37 | 35 | 29 | 6 | 2 | 4 | BD |
| 263 | 324 | 236 | 44 | 44 | 38 | 6 | 4 | 3 | ABDH |
| 264 | 333 | 250 | 40 | 43 | 37 | 6 | 6 | 4 | ABCEFG |
| 265 | 208 | 187 | 18 | 3 | 2 | 1 | 1 | 4 | H |
| 266 | 257 | 202 | 39 | 16 | 9 | 7 | 2 | 4 | CH |
| 267 | 368 | 250 | 49 | 69 | 42 | 27 | 3 | 3 | ADE |
| 268 | 240 | 205 | 24 | 11 | 11 | 0 | 1 | 3 | E |
| 269 | 351 | 272 | 43 | 36 | 34 | 2 | 2 | 3 | CD |
| 270 | 411 | 197 | 82 | 132 | 96 | 36 | 6 | 4 | **EFGH |
| 271 | 397 | 222 | 63 | 112 | 90 | 22 | 7 | 3 | ABCDEFH |
| 272 | 266 | 228 | 31 | 7 | 3 | 4 | 2 | 4 | EH |
| 273 | 369 | 209 | 69 | 91 | 73 | 18 | 4 | 4 | ABEF |

| | | | | | | | | | |
|---|---|---|---|---|---|---|---|---|---|
| 274 | 287 | 211 | 30 | 46 | 30 | 16 | 3 | 3 | ABH |
| 275 | 294 | 225 | 38 | 31 | 25 | 6 | 4 | 3 | ABDG |
| 276 | 370 | 200 | 61 | 109 | 80 | 29 | 4 | 1 | CEFH |
| 277 | 282 | 243 | 34 | 5 | 2 | 3 | 2 | 3 | AD |
| 278 | 444 | 239 | 81 | 124 | 86 | 38 | 8 | 4 | ABCDEFGH |
| 279 | 362 | 224 | 53 | 85 | 61 | 24 | 6 | 3 | ABCEGH |
| 280 | 432 | 291 | 68 | 73 | 57 | 16 | 4 | 4 | ABCG |

*Ping* code: Chr1:2640500..2640502 (*PingA*); Chr1:4220010..4220012 (*PingB*); Chr3:28019800..28019802 (*PingC*); Chr7:26460307..26460309 (*PingD*); Chr9:10863118..10863120 (*PingE*); Chr9:13736141..13736143 (*PingF*); Chr9:16690612..16690614 (*PingG*); Chr6:23521641..23526981 (*PingH*, Nipponbare *Ping*); New *Ping* (*)

Supplementary Data 3. List of excisions and footprints at parental *mPing* loci

| *mPing* loci | Number of excisions | Number of excisions with footprints | Number of RILs with excisions | Number of RILs with footprints | Footprints (number of RILs) |
|---|---|---|---|---|---|
| Chr1:10903901 | 0 | 0 | 0 | 0 | NA |
| Chr1:11226135 | 1 | 0 | 1 | 0 | Perfect(1) |
| Chr1:1132975 | 0 | 0 | 0 | 0 | NA |
| Chr1:12249228 | 1 | 0 | 1 | 0 | Perfect(1) |
| Chr1:13935275 | 0 | 0 | 0 | 0 | NA |
| Chr1:14047376 | 0 | 0 | 0 | 0 | NA |
| Chr1:1715117 | 1 | 0 | 1 | 0 | Perfect(1) |
| Chr1:17513834 | 1 | 1 | 1 | 1 | I_TGTGACACAG(1) |
| Chr1:18989742 | 2 | 2 | 2 | 2 | D_4-D_2(1);I_GACC(1) |
| Chr1:21351851 | 0 | 0 | 0 | 0 | NA |
| Chr1:21565700 | 0 | 0 | 0 | 0 | NA |
| Chr1:22415370 | 0 | 0 | 0 | 0 | NA |
| Chr1:23332547 | 2 | 1 | 2 | 1 | D_22(1);Perfect(1) |
| Chr1:24082898 | 0 | 0 | 0 | 0 | NA |
| Chr1:24610175 | 0 | 0 | 0 | 0 | NA |
| Chr1:24779771 | 2 | 1 | 2 | 1 | I_A(1);Perfect(1) |
| Chr1:25256701 | 1 | 1 | 1 | 1 | D_2(1) |
| Chr1:25261112 | 0 | 0 | 0 | 0 | NA |
| Chr1:25680936 | 0 | 0 | 0 | 0 | NA |
| Chr1:28356229 | 0 | 0 | 0 | 0 | NA |
| Chr1:28924285 | 0 | 0 | 0 | 0 | NA |
| Chr1:29276379 | 0 | 0 | 0 | 0 | NA |
| Chr1:29322743 | 1 | 1 | 1 | 1 | I_AAAACGAGGAT(1) |
| Chr1:29328836 | 2 | 1 | 2 | 1 | D_7(1);Perfect(1) |

| | | | | | |
|---|---|---|---|---|---|
| Chr1:29494572 | 5 | 4 | 7 | 4 | D_17(1);D_6(1);I_A(1);I_AA(1);Perfect(3) |
| Chr1:29926801 | 1 | 0 | 1 | 0 | Perfect(1) |
| Chr1:29931517 | 2 | 1 | 3 | 2 | D_9(2);Perfect(1) |
| Chr1:30895565 | 0 | 0 | 0 | 0 | NA |
| Chr1:31556970 | 0 | 0 | 0 | 0 | NA |
| Chr1:32293360 | 1 | 0 | 1 | 0 | Perfect(1) |
| Chr1:34086312 | 1 | 0 | 1 | 0 | Perfect(1) |
| Chr1:34281436 | 0 | 0 | 0 | 0 | NA |
| Chr1:3444992 | 5 | 4 | 8 | 4 | D_8(1);D_5(1);I_TTTTC-I_TC-I_CTTG(1);D_2(1);Perfect(4) |
| Chr1:34897489 | 0 | 0 | 0 | 0 | NA |
| Chr1:35465445 | 0 | 0 | 0 | 0 | NA |
| Chr1:35607599 | 1 | 1 | 1 | 1 | D_2(1) |
| Chr1:36114224 | 0 | 0 | 0 | 0 | NA |
| Chr1:36149918 | 0 | 0 | 0 | 0 | NA |
| Chr1:36267659 | 5 | 4 | 57 | 43 | D_5(40);D_12(1);D_2(1);D_9(1);Perfect(14) |
| Chr1:36270511 | 7 | 6 | 50 | 38 | D_31(1);D_20(32);D_23(2);D_16(1);D_4(1);I_T(1);Perfect(12) |
| Chr1:3780849 | 0 | 0 | 0 | 0 | NA |
| Chr1:37828361 | 0 | 0 | 0 | 0 | NA |
| Chr1:38092038 | 0 | 0 | 0 | 0 | NA |
| Chr1:38674212 | 0 | 0 | 0 | 0 | NA |
| Chr1:38859876 | 2 | 1 | 2 | 1 | D_5(1);Perfect(1) |
| Chr1:38898352 | 1 | 1 | 1 | 1 | D_17(1) |
| Chr1:39147394 | 1 | 0 | 1 | 0 | Perfect(1) |
| Chr1:39212320 | 2 | 1 | 2 | 1 | D_8(1);Perfect(1) |
| Chr1:3929765 | 2 | 2 | 2 | 2 | D_5(1);I_AC(1) |
| Chr1:39543338 | 2 | 1 | 2 | 1 | D_2-D_9(1);Perfect(1) |

| | | | | | |
|---|---|---|---|---|---|
| Chr1:39957257 | 0 | 0 | 0 | 0 | NA |
| Chr1:40469772 | 0 | 0 | 0 | 0 | NA |
| Chr1:40641754 | 0 | 0 | 0 | 0 | NA |
| Chr1:4076464 | 3 | 2 | 3 | 2 | D_4(1);D_2(1);Perfect(1) |
| Chr1:40818575 | 0 | 0 | 0 | 0 | NA |
| Chr1:40832862 | 0 | 0 | 0 | 0 | NA |
| Chr1:4101860 | 2 | 2 | 2 | 2 | D_19(1);I_A(1) |
| Chr1:41148511 | 0 | 0 | 0 | 0 | NA |
| Chr1:41818259 | 0 | 0 | 0 | 0 | NA |
| Chr1:42156581 | 0 | 0 | 0 | 0 | NA |
| Chr1:42599829 | 0 | 0 | 0 | 0 | NA |
| Chr1:42930250 | 0 | 0 | 0 | 0 | NA |
| Chr1:6172233 | 2 | 1 | 2 | 1 | D_16(1);Perfect(1) |
| Chr1:6806761 | 7 | 6 | 10 | 9 | D_2-D_19-I_A(1);D_13(1);D_12(1);D_4(1);D_3(4);I_TTA(1);Perfect(1) |
| Chr1:6816415 | 4 | 3 | 4 | 3 | D_10(1);D_2(1);D_3(1);Perfect(1) |
| Chr1:6846279 | 1 | 0 | 2 | 0 | Perfect(2) |
| Chr1:6864904 | 3 | 2 | 3 | 2 | D_9(1);D_20(1);Perfect(1) |
| Chr1:7514412 | 1 | 0 | 1 | 0 | Perfect(1) |
| Chr1:8479732 | 0 | 0 | 0 | 0 | NA |
| Chr1:8750744 | 0 | 0 | 0 | 0 | NA |
| Chr1:9142465 | 0 | 0 | 0 | 0 | NA |
| Chr1:9581605 | 1 | 0 | 1 | 0 | Perfect(1) |
| Chr10:11955070 | 0 | 0 | 0 | 0 | NA |
| Chr10:13102744 | 0 | 0 | 0 | 0 | NA |
| Chr10:13370541 | 0 | 0 | 0 | 0 | NA |
| Chr10:14020306 | 0 | 0 | 0 | 0 | NA |
| Chr10:16851282 | 2 | 1 | 4 | 1 | D_1-D_1-D_3-D_1(1);Perfect(3) |
| Chr10:17496871 | 0 | 0 | 0 | 0 | NA |

| | | | | | |
|---|---|---|---|---|---|
| Chr10:18044214 | 0 | 0 | 0 | 0 | NA |
| Chr10:18765103 | 1 | 1 | 1 | 1 | I_T(1) |
| Chr10:21163118 | 0 | 0 | 0 | 0 | NA |
| Chr10:21361206 | 0 | 0 | 0 | 0 | NA |
| Chr10:21716391 | 1 | 1 | 1 | 1 | D_1(1) |
| Chr10:22510407 | 0 | 0 | 0 | 0 | NA |
| Chr10:22578379 | 0 | 0 | 0 | 0 | NA |
| Chr10:2734624 | 0 | 0 | 0 | 0 | NA |
| Chr10:2854153 | 0 | 0 | 0 | 0 | NA |
| Chr10:2854243 | 0 | 0 | 0 | 0 | NA |
| Chr10:4007946 | 3 | 2 | 3 | 2 | I_TAAG(1);I_AA(1);Perfect(1) |
| Chr10:4320421 | 0 | 0 | 0 | 0 | NA |
| Chr10:6566243 | 0 | 0 | 0 | 0 | NA |
| Chr10:6896566 | 0 | 0 | 0 | 0 | NA |
| Chr10:9169250 | 1 | 1 | 1 | 1 | D_7-D_4(1) |
| Chr11:10797353 | 0 | 0 | 0 | 0 | NA |
| Chr11:10863129 | 1 | 1 | 1 | 1 | D_11(1) |
| Chr11:14340125 | 0 | 0 | 0 | 0 | NA |
| Chr11:14578665 | 0 | 0 | 0 | 0 | NA |
| Chr11:14670786 | 1 | 1 | 1 | 1 | D_2(1) |
| Chr11:16397045 | 1 | 1 | 1 | 1 | D_7(1) |
| Chr11:16894772 | 0 | 0 | 0 | 0 | NA |
| Chr11:19084257 | 0 | 0 | 0 | 0 | NA |
| Chr11:19939294 | 1 | 0 | 1 | 0 | Perfect(1) |
| Chr11:20490425 | 0 | 0 | 0 | 0 | NA |
| Chr11:21176364 | 0 | 0 | 0 | 0 | NA |
| Chr11:21459184 | 0 | 0 | 0 | 0 | NA |
| Chr11:22474609 | 0 | 0 | 0 | 0 | NA |
| Chr11:22938098 | 0 | 0 | 0 | 0 | NA |
| Chr11:23200105 | 0 | 0 | 0 | 0 | NA |

| | | | | | |
|---|---|---|---|---|---|
| Chr11:24682792 | 0 | 0 | 0 | 0 | NA |
| Chr11:25523876 | 0 | 0 | 0 | 0 | NA |
| Chr11:26077404 | 1 | 1 | 1 | 1 | D_13(1) |
| Chr11:28625682 | 0 | 0 | 0 | 0 | NA |
| Chr11:3851785 | 0 | 0 | 0 | 0 | NA |
| Chr11:393598 | 0 | 0 | 0 | 0 | NA |
| Chr11:4179066 | 1 | 0 | 2 | 0 | Perfect(2) |
| Chr11:6969696 | 2 | 1 | 4 | 1 | D_1-D_2-I_A(1);Perfect(3) |
| Chr11:714790 | 0 | 0 | 0 | 0 | NA |
| Chr11:7473250 | 1 | 1 | 1 | 1 | D_2(1) |
| Chr11:7503886 | 2 | 1 | 3 | 1 | D_2(1);Perfect(2) |
| Chr12:1045463 | 1 | 0 | 1 | 0 | Perfect(1) |
| Chr12:10612331 | 0 | 0 | 0 | 0 | NA |
| Chr12:10822439 | 0 | 0 | 0 | 0 | NA |
| Chr12:19177973 | 0 | 0 | 0 | 0 | NA |
| Chr12:23877483 | 1 | 0 | 1 | 0 | Perfect(1) |
| Chr12:24595103 | 1 | 1 | 1 | 1 | D_11(1) |
| Chr12:26261130 | 0 | 0 | 0 | 0 | NA |
| Chr12:26642870 | 0 | 0 | 0 | 0 | NA |
| Chr12:2734541 | 0 | 0 | 0 | 0 | NA |
| Chr12:3119686 | 1 | 1 | 1 | 1 | I_CCTCC-I_CT(1) |
| Chr12:3285787 | 0 | 0 | 0 | 0 | NA |
| Chr12:332029 | 0 | 0 | 0 | 0 | NA |
| Chr12:3373257 | 0 | 0 | 0 | 0 | NA |
| Chr12:3616590 | 0 | 0 | 0 | 0 | NA |
| Chr12:4097962 | 0 | 0 | 0 | 0 | NA |
| Chr12:4965478 | 0 | 0 | 0 | 0 | NA |
| Chr12:6625424 | 0 | 0 | 0 | 0 | NA |
| Chr12:7210157 | 0 | 0 | 0 | 0 | NA |
| Chr12:839604 | 1 | 1 | 1 | 1 | I_TTA(1) |

| | | | | | |
|---|---|---|---|---|---|
| Chr12:8852564 | 0 | 0 | 0 | 0 | NA |
| Chr12:9350665 | 1 | 0 | 2 | 0 | Perfect(2) |
| Chr12:9951106 | 0 | 0 | 0 | 0 | NA |
| Chr2:10076667 | 1 | 1 | 1 | 1 | D_11(1) |
| Chr2:1161811 | 0 | 0 | 0 | 0 | NA |
| Chr2:12467700 | 2 | 1 | 3 | 1 | D_3-D_5(1);Perfect(2) |
| Chr2:13161938 | 0 | 0 | 0 | 0 | NA |
| Chr2:13200714 | 1 | 0 | 2 | 0 | Perfect(2) |
| Chr2:14601853 | 4 | 3 | 51 | 43 | D_6(1);D_6(41);D_1-D_5(1);Perfect(8) |
| Chr2:15727562 | 1 | 0 | 56 | 0 | Perfect(56) |
| Chr2:1583547 | 1 | 0 | 1 | 0 | Perfect(1) |
| Chr2:18534787 | 1 | 0 | 1 | 0 | Perfect(1) |
| Chr2:18569683 | 1 | 1 | 2 | 2 | D_8(2) |
| Chr2:20000327 | 1 | 0 | 1 | 0 | Perfect(1) |
| Chr2:20144823 | 0 | 0 | 0 | 0 | NA |
| Chr2:2025376 | 1 | 1 | 1 | 1 | D_12(1) |
| Chr2:20589717 | 2 | 1 | 2 | 1 | D_28(1);Perfect(1) |
| Chr2:20722279 | 0 | 0 | 0 | 0 | NA |
| Chr2:21001168 | 0 | 0 | 0 | 0 | NA |
| Chr2:21319934 | 1 | 0 | 1 | 0 | Perfect(1) |
| Chr2:21443360 | 0 | 0 | 0 | 0 | NA |
| Chr2:214437 | 0 | 0 | 0 | 0 | NA |
| Chr2:223440 | 2 | 1 | 2 | 1 | D_2(1);Perfect(1) |
| Chr2:22478977 | 0 | 0 | 0 | 0 | NA |
| Chr2:22549115 | 0 | 0 | 0 | 0 | NA |
| Chr2:23882557 | 0 | 0 | 0 | 0 | NA |
| Chr2:24014618 | 2 | 1 | 2 | 1 | D_1(1);Perfect(1) |
| Chr2:25548875 | 0 | 0 | 0 | 0 | NA |
| Chr2:25969392 | 0 | 0 | 0 | 0 | NA |

| | | | | | |
|---|---|---|---|---|---|
| Chr2:26510355 | 0 | 0 | 0 | 0 | NA |
| Chr2:27162235 | 0 | 0 | 0 | 0 | NA |
| Chr2:27244610 | 0 | 0 | 0 | 0 | NA |
| Chr2:27685185 | 1 | 1 | 1 | 1 | D_3(1) |
| Chr2:28008341 | 1 | 0 | 1 | 0 | Perfect(1) |
| Chr2:28376797 | 0 | 0 | 0 | 0 | NA |
| Chr2:28662452 | 0 | 0 | 0 | 0 | NA |
| Chr2:28724476 | 3 | 3 | 4 | 4 | D_10(1);D_3(2);I_A(1) |
| Chr2:28732623 | 4 | 3 | 6 | 5 | D_14(1);D_12(2);D_9(2);Perfect(1) |
| Chr2:29244327 | 0 | 0 | 0 | 0 | NA |
| Chr2:29429219 | 0 | 0 | 0 | 0 | NA |
| Chr2:29547915 | 2 | 2 | 2 | 2 | D_22(1);D_2-D_1(1) |
| Chr2:29576700 | 0 | 0 | 0 | 0 | NA |
| Chr2:30104937 | 3 | 2 | 6 | 3 | D_3(2);I_TT(1);Perfect(3) |
| Chr2:30104997 | 1 | 1 | 1 | 1 | D_1-D_6(1) |
| Chr2:30934395 | 6 | 5 | 8 | 5 | D_1(1);D_3(1);I_TT(1);I_TTA(1);I_TTAAC(1);Perfect(3) |
| Chr2:31455301 | 1 | 0 | 1 | 0 | Perfect(1) |
| Chr2:31636377 | 0 | 0 | 0 | 0 | NA |
| Chr2:32397174 | 1 | 0 | 2 | 0 | Perfect(2) |
| Chr2:32572258 | 0 | 0 | 0 | 0 | NA |
| Chr2:32858536 | 0 | 0 | 0 | 0 | NA |
| Chr2:32895944 | 0 | 0 | 0 | 0 | NA |
| Chr2:33871069 | 0 | 0 | 0 | 0 | NA |
| Chr2:34670530 | 1 | 1 | 1 | 1 | D_21(1) |
| Chr2:3502594 | 0 | 0 | 0 | 0 | NA |
| Chr2:3639031 | 0 | 0 | 0 | 0 | NA |
| Chr2:4314535 | 1 | 1 | 1 | 1 | D_16(1) |
| Chr2:617949 | 0 | 0 | 0 | 0 | NA |
| Chr2:7113420 | 1 | 0 | 1 | 0 | Perfect(1) |

| | | | | | |
|---|---|---|---|---|---|
| Chr2:8187917 | 0 | 0 | 0 | 0 | NA |
| Chr2:8898160 | 0 | 0 | 0 | 0 | NA |
| Chr2:913613 | 0 | 0 | 0 | 0 | NA |
| Chr3:10593111 | 0 | 0 | 0 | 0 | NA |
| Chr3:10990369 | 0 | 0 | 0 | 0 | NA |
| Chr3:11329080 | 1 | 0 | 1 | 0 | Perfect(1) |
| Chr3:11781493 | 0 | 0 | 0 | 0 | NA |
| Chr3:11922190 | 1 | 0 | 1 | 0 | Perfect(1) |
| Chr3:12036120 | 0 | 0 | 0 | 0 | NA |
| Chr3:12067613 | 0 | 0 | 0 | 0 | NA |
| Chr3:12409837 | 1 | 0 | 2 | 0 | Perfect(2) |
| Chr3:1247288 | 0 | 0 | 0 | 0 | NA |
| Chr3:12735756 | 2 | 2 | 2 | 2 | D_12(1);I_TTA(1) |
| Chr3:1299651 | 0 | 0 | 0 | 0 | NA |
| Chr3:13027116 | 1 | 1 | 1 | 1 | D_15(1) |
| Chr3:13281744 | 1 | 0 | 1 | 0 | Perfect(1) |
| Chr3:13683979 | 0 | 0 | 0 | 0 | NA |
| Chr3:14597027 | 0 | 0 | 0 | 0 | NA |
| Chr3:15487801 | 1 | 0 | 1 | 0 | Perfect(1) |
| Chr3:15698486 | 0 | 0 | 0 | 0 | NA |
| Chr3:16222140 | 0 | 0 | 0 | 0 | NA |
| Chr3:17195960 | 1 | 0 | 1 | 0 | Perfect(1) |
| Chr3:17575717 | 0 | 0 | 0 | 0 | NA |
| Chr3:17642391 | 0 | 0 | 0 | 0 | NA |
| Chr3:198572 | 0 | 0 | 0 | 0 | NA |
| Chr3:21026572 | 1 | 0 | 1 | 0 | Perfect(1) |
| Chr3:21360120 | 0 | 0 | 0 | 0 | NA |
| Chr3:22701631 | 2 | 1 | 17 | 16 | D_25(16);Perfect(1) |
| Chr3:24436164 | 1 | 0 | 1 | 0 | Perfect(1) |
| Chr3:24855529 | 2 | 1 | 2 | 1 | D_7(1);Perfect(1) |

| | | | | | |
|---|---|---|---|---|---|
| Chr3:24920217 | 0 | 0 | 0 | 0 | NA |
| Chr3:26028786 | 0 | 0 | 0 | 0 | NA |
| Chr3:26753248 | 0 | 0 | 0 | 0 | NA |
| Chr3:2675912 | 1 | 0 | 36 | 0 | Perfect(36) |
| Chr3:28088459 | 1 | 0 | 3 | 0 | Perfect(3) |
| Chr3:28365657 | 0 | 0 | 0 | 0 | NA |
| Chr3:29404858 | 0 | 0 | 0 | 0 | NA |
| Chr3:29404901 | 0 | 0 | 0 | 0 | NA |
| Chr3:30784687 | 0 | 0 | 0 | 0 | NA |
| Chr3:31364757 | 0 | 0 | 0 | 0 | NA |
| Chr3:31683285 | 1 | 1 | 1 | 1 | D_1(1) |
| Chr3:31890843 | 1 | 1 | 1 | 1 | I_TTC(1) |
| Chr3:33362073 | 0 | 0 | 0 | 0 | NA |
| Chr3:34077806 | 0 | 0 | 0 | 0 | NA |
| Chr3:34592545 | 1 | 1 | 1 | 1 | D_3(1) |
| Chr3:35176773 | 0 | 0 | 0 | 0 | NA |
| Chr3:35894895 | 0 | 0 | 0 | 0 | NA |
| Chr3:35945979 | 0 | 0 | 0 | 0 | NA |
| Chr3:3623527 | 0 | 0 | 0 | 0 | NA |
| Chr3:3804370 | 0 | 0 | 0 | 0 | NA |
| Chr3:399055 | 1 | 0 | 1 | 0 | Perfect(1) |
| Chr3:4377252 | 0 | 0 | 0 | 0 | NA |
| Chr3:511329 | 0 | 0 | 0 | 0 | NA |
| Chr3:5504299 | 0 | 0 | 0 | 0 | NA |
| Chr3:6154020 | 0 | 0 | 0 | 0 | NA |
| Chr3:6328408 | 1 | 0 | 1 | 0 | Perfect(1) |
| Chr3:6441699 | 8 | 7 | 19 | 8 | D_18(2);D_6(1);D_2(1);D_7(1);I_TTAT(1);I_AAATACA(1);D_26(1);Perfect(11) |
| Chr3:6512009 | 1 | 0 | 1 | 0 | Perfect(1) |

| | | | | | |
|---|---|---|---|---|---|
| Chr3:6513589 | 0 | 0 | 0 | 0 | NA |
| Chr3:7296367 | 0 | 0 | 0 | 0 | NA |
| Chr3:9014023 | 2 | 1 | 2 | 1 | D_1-D_11-D_10(1);Perfect(1) |
| Chr3:9162954 | 0 | 0 | 0 | 0 | NA |
| Chr3:9240074 | 1 | 0 | 1 | 0 | Perfect(1) |
| Chr3:9427120 | 0 | 0 | 0 | 0 | NA |
| Chr3:9568670 | 0 | 0 | 0 | 0 | NA |
| Chr3:982231 | 0 | 0 | 0 | 0 | NA |
| Chr4:13471891 | 0 | 0 | 0 | 0 | NA |
| Chr4:15070775 | 0 | 0 | 0 | 0 | NA |
| Chr4:18566524 | 0 | 0 | 0 | 0 | NA |
| Chr4:1877633 | 0 | 0 | 0 | 0 | NA |
| Chr4:19021060 | 0 | 0 | 0 | 0 | NA |
| Chr4:19471782 | 0 | 0 | 0 | 0 | NA |
| Chr4:19922773 | 0 | 0 | 0 | 0 | NA |
| Chr4:19999252 | 0 | 0 | 0 | 0 | NA |
| Chr4:20249932 | 0 | 0 | 0 | 0 | NA |
| Chr4:21600764 | 1 | 1 | 1 | 1 | I_A(1) |
| Chr4:21673101 | 0 | 0 | 0 | 0 | NA |
| Chr4:21998764 | 2 | 2 | 2 | 2 | D_23(1);I_TTTAAGATCTAATC(1) |
| Chr4:23199000 | 0 | 0 | 0 | 0 | NA |
| Chr4:23439376 | 1 | 0 | 1 | 0 | Perfect(1) |
| Chr4:23571130 | 0 | 0 | 0 | 0 | NA |
| Chr4:24208721 | 0 | 0 | 0 | 0 | NA |
| Chr4:25029910 | 0 | 0 | 0 | 0 | NA |
| Chr4:26020117 | 0 | 0 | 0 | 0 | NA |
| Chr4:26982348 | 0 | 0 | 0 | 0 | NA |
| Chr4:27475745 | 0 | 0 | 0 | 0 | NA |
| Chr4:28289923 | 0 | 0 | 0 | 0 | NA |
| Chr4:28739797 | 0 | 0 | 0 | 0 | NA |

| | | | | | |
|---|---|---|---|---|---|
| Chr4:28903396 | 0 | 0 | 0 | 0 | NA |
| Chr4:29218325 | 0 | 0 | 0 | 0 | NA |
| Chr4:29603863 | 1 | 1 | 1 | 1 | D_1-D_4-D_8(1) |
| Chr4:30320688 | 1 | 0 | 1 | 0 | Perfect(1) |
| Chr4:30513997 | 0 | 0 | 0 | 0 | NA |
| Chr4:31677002 | 0 | 0 | 0 | 0 | NA |
| Chr4:31997420 | 1 | 1 | 1 | 1 | D_8(1) |
| Chr4:32326935 | 1 | 0 | 1 | 0 | Perfect(1) |
| Chr4:33017281 | 0 | 0 | 0 | 0 | NA |
| Chr4:33021868 | 0 | 0 | 0 | 0 | NA |
| Chr4:33507343 | 2 | 1 | 6 | 3 | D_8(3);Perfect(3) |
| Chr4:33861550 | 1 | 0 | 1 | 0 | Perfect(1) |
| Chr4:34302847 | 0 | 0 | 0 | 0 | NA |
| Chr4:34688306 | 0 | 0 | 0 | 0 | NA |
| Chr4:35244614 | 0 | 0 | 0 | 0 | NA |
| Chr4:35421806 | 0 | 0 | 0 | 0 | NA |
| Chr4:35422419 | 0 | 0 | 0 | 0 | NA |
| Chr5:14154610 | 1 | 0 | 1 | 0 | Perfect(1) |
| Chr5:1441039 | 0 | 0 | 0 | 0 | NA |
| Chr5:14654628 | 0 | 0 | 0 | 0 | NA |
| Chr5:14878712 | 1 | 0 | 1 | 0 | Perfect(1) |
| Chr5:15210313 | 9 | 8 | 18 | 9 | D_25(1);D_12(1);D_15(1);I_A-D_3(1);D_17(2);I_CAATCAAATT(1);I_TT(1);I_A(1);Perfect(9) |
| Chr5:15213006 | 10 | 9 | 14 | 9 | D_11-D_23(1);D_23(1);D_15(1);D_9(1);D_7(1);D_9(1);D_12(1);I_TTA(1);D_1(1);Perfect(5) |
| Chr5:15488460 | 0 | 0 | 0 | 0 | NA |
| Chr5:15586258 | 0 | 0 | 0 | 0 | NA |

| | | | | | |
|---|---|---|---|---|---|
| Chr5:15698449 | 0 | 0 | 0 | 0 | NA |
| Chr5:16364136 | 0 | 0 | 0 | 0 | NA |
| Chr5:16615709 | 0 | 0 | 0 | 0 | NA |
| Chr5:18747498 | 0 | 0 | 0 | 0 | NA |
| Chr5:19143339 | 1 | 0 | 1 | 0 | Perfect(1) |
| Chr5:19328618 | 1 | 0 | 1 | 0 | Perfect(1) |
| Chr5:19961220 | 2 | 1 | 2 | 1 | D_6(1);Perfect(1) |
| Chr5:20117985 | 1 | 0 | 1 | 0 | Perfect(1) |
| Chr5:21633282 | 0 | 0 | 0 | 0 | NA |
| Chr5:22056644 | 0 | 0 | 0 | 0 | NA |
| Chr5:22235594 | 1 | 0 | 1 | 0 | Perfect(1) |
| Chr5:22646609 | 1 | 0 | 1 | 0 | Perfect(1) |
| Chr5:226600 | 1 | 0 | 1 | 0 | Perfect(1) |
| Chr5:23257994 | 0 | 0 | 0 | 0 | NA |
| Chr5:23730117 | 3 | 2 | 6 | 2 | I_A(1);I_ATTG(1);Perfect(4) |
| Chr5:23730902 | 2 | 1 | 6 | 1 | D_29(1);Perfect(5) |
| Chr5:24878619 | 0 | 0 | 0 | 0 | NA |
| Chr5:25474861 | 1 | 1 | 1 | 1 | D_1(1) |
| Chr5:25536621 | 0 | 0 | 0 | 0 | NA |
| Chr5:25546566 | 2 | 1 | 2 | 1 | D_3-D_1-I_A-I_C(1);Perfect(1) |
| Chr5:26247575 | 0 | 0 | 0 | 0 | NA |
| Chr5:28151164 | 0 | 0 | 0 | 0 | NA |
| Chr5:28927631 | 0 | 0 | 0 | 0 | NA |
| Chr5:4249078 | 0 | 0 | 0 | 0 | NA |
| Chr5:5370456 | 1 | 0 | 1 | 0 | Perfect(1) |
| Chr5:6164428 | 2 | 1 | 2 | 1 | D_4(1);Perfect(1) |
| Chr5:772694 | 2 | 1 | 2 | 1 | I_TTATCTCTGAATGT(1);Perfect(1) |
| Chr6:10360943 | 0 | 0 | 0 | 0 | NA |
| Chr6:13737618 | 0 | 0 | 0 | 0 | NA |
| Chr6:14533008 | 2 | 1 | 3 | 1 | I_G(1);Perfect(2) |

| | | | | | |
|---|---|---|---|---|---|
| Chr6:14790333 | 1 | 0 | 1 | 0 | Perfect(1) |
| Chr6:17106306 | 1 | 0 | 1 | 0 | Perfect(1) |
| Chr6:18136422 | 0 | 0 | 0 | 0 | NA |
| Chr6:18886270 | 0 | 0 | 0 | 0 | NA |
| Chr6:2035335 | 2 | 2 | 2 | 2 | D_11(1);I_AATCAGTGTCTG(1) |
| Chr6:20717707 | 0 | 0 | 0 | 0 | NA |
| Chr6:21010525 | 0 | 0 | 0 | 0 | NA |
| Chr6:22646015 | 0 | 0 | 0 | 0 | NA |
| Chr6:23126474 | 0 | 0 | 0 | 0 | NA |
| Chr6:23535331 | 0 | 0 | 0 | 0 | NA |
| Chr6:24594639 | 0 | 0 | 0 | 0 | NA |
| Chr6:25002506 | 0 | 0 | 0 | 0 | NA |
| Chr6:26083883 | 0 | 0 | 0 | 0 | NA |
| Chr6:27840591 | 0 | 0 | 0 | 0 | NA |
| Chr6:2856376 | 1 | 0 | 1 | 0 | Perfect(1) |
| Chr6:29048879 | 0 | 0 | 0 | 0 | NA |
| Chr6:30049593 | 0 | 0 | 0 | 0 | NA |
| Chr6:30099538 | 2 | 2 | 2 | 2 | D_5(1);D_2(1) |
| Chr6:30293899 | 0 | 0 | 0 | 0 | NA |
| Chr6:30450783 | 0 | 0 | 0 | 0 | NA |
| Chr6:4093156 | 0 | 0 | 0 | 0 | NA |
| Chr6:4729737 | 0 | 0 | 0 | 0 | NA |
| Chr6:5711665 | 0 | 0 | 0 | 0 | NA |
| Chr6:6151042 | 1 | 1 | 1 | 1 | D_14(1) |
| Chr6:6356702 | 0 | 0 | 0 | 0 | NA |
| Chr6:6521725 | 0 | 0 | 0 | 0 | NA |
| Chr6:6848916 | 0 | 0 | 0 | 0 | NA |
| Chr6:7690782 | 0 | 0 | 0 | 0 | NA |
| Chr6:8042613 | 1 | 1 | 1 | 1 | I_TT(1) |

| | | | | | |
|---|---|---|---|---|---|
| Chr6:8316574 | 5 | 5 | 6 | 6 | D_14(1);D_21(2);I_TTA(1);D_10(1);D_5(1) |
| Chr6:8356631 | 0 | 0 | 0 | 0 | NA |
| Chr6:844779 | 0 | 0 | 0 | 0 | NA |
| Chr6:986514 | 0 | 0 | 0 | 0 | NA |
| Chr7:11852825 | 0 | 0 | 0 | 0 | NA |
| Chr7:13481675 | 1 | 1 | 1 | 1 | D_19(1) |
| Chr7:14975641 | 0 | 0 | 0 | 0 | NA |
| Chr7:16023232 | 0 | 0 | 0 | 0 | NA |
| Chr7:17125850 | 0 | 0 | 0 | 0 | NA |
| Chr7:18920532 | 0 | 0 | 0 | 0 | NA |
| Chr7:20152299 | 1 | 1 | 1 | 1 | D_2(1) |
| Chr7:20204538 | 0 | 0 | 0 | 0 | NA |
| Chr7:20615708 | 1 | 1 | 1 | 1 | D_9(1) |
| Chr7:21376225 | 0 | 0 | 0 | 0 | NA |
| Chr7:22060085 | 1 | 0 | 1 | 0 | Perfect(1) |
| Chr7:22679547 | 0 | 0 | 0 | 0 | NA |
| Chr7:23009797 | 1 | 1 | 1 | 1 | I_T(1) |
| Chr7:23272150 | 0 | 0 | 0 | 0 | NA |
| Chr7:23752427 | 0 | 0 | 0 | 0 | NA |
| Chr7:23852999 | 2 | 2 | 2 | 2 | D_13(1);I_TAGTA(1) |
| Chr7:23898253 | 0 | 0 | 0 | 0 | NA |
| Chr7:24937108 | 0 | 0 | 0 | 0 | NA |
| Chr7:25772025 | 0 | 0 | 0 | 0 | NA |
| Chr7:25788368 | 2 | 1 | 33 | 24 | D_16(24);Perfect(9) |
| Chr7:26090554 | 2 | 1 | 2 | 1 | D_20(1);Perfect(1) |
| Chr7:26156682 | 1 | 0 | 1 | 0 | Perfect(1) |
| Chr7:27443190 | 0 | 0 | 0 | 0 | NA |
| Chr7:28151483 | 0 | 0 | 0 | 0 | NA |
| Chr7:28821663 | 0 | 0 | 0 | 0 | NA |

| | | | | | |
|---|---|---|---|---|---|
| Chr7:29015072 | 0 | 0 | 0 | 0 | NA |
| Chr7:29475470 | 4 | 3 | 6 | 4 | I_TA(1);D_25(2);I_AA(1);Perfect(2) |
| Chr7:29484349 | 0 | 0 | 0 | 0 | NA |
| Chr7:3667745 | 3 | 2 | 4 | 2 | D_17(1);D_2(1);Perfect(2) |
| Chr7:4019952 | 0 | 0 | 0 | 0 | NA |
| Chr7:4560975 | 0 | 0 | 0 | 0 | NA |
| Chr7:7314674 | 1 | 0 | 1 | 0 | Perfect(1) |
| Chr7:7972038 | 1 | 0 | 1 | 0 | Perfect(1) |
| Chr7:8879928 | 0 | 0 | 0 | 0 | NA |
| Chr8:1019672 | 0 | 0 | 0 | 0 | NA |
| Chr8:11403145 | 0 | 0 | 0 | 0 | NA |
| Chr8:13076782 | 6 | 5 | 6 | 5 | D_15(1);D_14(1);I_C(1);I_AA(1);I_TGAAG(1);Perfect(1) |
| Chr8:13103108 | 1 | 0 | 1 | 0 | Perfect(1) |
| Chr8:14721482 | 0 | 0 | 0 | 0 | NA |
| Chr8:15387727 | 1 | 0 | 1 | 0 | Perfect(1) |
| Chr8:16683588 | 0 | 0 | 0 | 0 | NA |
| Chr8:19197271 | 0 | 0 | 0 | 0 | NA |
| Chr8:19452248 | 0 | 0 | 0 | 0 | NA |
| Chr8:20442416 | 0 | 0 | 0 | 0 | NA |
| Chr8:20628372 | 0 | 0 | 0 | 0 | NA |
| Chr8:20674237 | 0 | 0 | 0 | 0 | NA |
| Chr8:22948360 | 4 | 4 | 4 | 4 | D_13(1);D_37(1);I_A(1);I_AGAATAGAGGG(1) |
| Chr8:23290289 | 0 | 0 | 0 | 0 | NA |
| Chr8:23802328 | 0 | 0 | 0 | 0 | NA |
| Chr8:24585356 | 0 | 0 | 0 | 0 | NA |

| | | | | | |
|---|---|---|---|---|---|
| Chr8:24622994 | 15 | 14 | 28 | 14 | D_9(1);D_8(1);D_20(1);D_15(1);D_8(1);D_13(1);D_11(1);D_3(1);D_4-D_29(1);I_T(1);I_TT(1);D_14-D_4-I_C(1);D_2(1);D_1(1);Perfect(14) |
| Chr8:24830426 | 0 | 0 | 0 | 0 | NA |
| Chr8:24965704 | 0 | 0 | 0 | 0 | NA |
| Chr8:25236840 | 1 | 0 | 1 | 0 | Perfect(1) |
| Chr8:25278859 | 0 | 0 | 0 | 0 | NA |
| Chr8:25881178 | 1 | 1 | 1 | 1 | I_TAA(1) |
| Chr8:27109715 | 0 | 0 | 0 | 0 | NA |
| Chr8:27153071 | 1 | 0 | 1 | 0 | Perfect(1) |
| Chr8:28186241 | 0 | 0 | 0 | 0 | NA |
| Chr8:458415 | 0 | 0 | 0 | 0 | NA |
| Chr8:4712970 | 1 | 1 | 1 | 1 | D_4(1) |
| Chr8:737334 | 0 | 0 | 0 | 0 | NA |
| Chr8:8581520 | 1 | 0 | 1 | 0 | Perfect(1) |
| Chr8:8655022 | 0 | 0 | 0 | 0 | NA |
| Chr8:9385127 | 1 | 0 | 1 | 0 | Perfect(1) |
| Chr8:973880 | 0 | 0 | 0 | 0 | NA |
| Chr8:9778572 | 1 | 1 | 1 | 1 | D_10(1) |
| Chr9:14118723 | 0 | 0 | 0 | 0 | NA |
| Chr9:15394515 | 0 | 0 | 0 | 0 | NA |
| Chr9:15441738 | 1 | 1 | 1 | 1 | I_TT(1) |
| Chr9:15873914 | 0 | 0 | 0 | 0 | NA |
| Chr9:16024218 | 0 | 0 | 0 | 0 | NA |
| Chr9:16330820 | 1 | 1 | 1 | 1 | D_3(1) |
| Chr9:16698141 | 0 | 0 | 0 | 0 | NA |
| Chr9:17452747 | 0 | 0 | 0 | 0 | NA |
| Chr9:17901601 | 0 | 0 | 0 | 0 | NA |
| Chr9:18354394 | 0 | 0 | 0 | 0 | NA |

| | | | | | |
|---|---|---|---|---|---|
| Chr9:18705899 | 2 | 1 | 2 | 1 | I_T(1);Perfect(1) |
| Chr9:19596969 | 1 | 1 | 1 | 1 | I_T(1) |
| Chr9:19832667 | 2 | 1 | 55 | 1 | D_1(1);Perfect(54) |
| Chr9:20177227 | 1 | 0 | 1 | 0 | Perfect(1) |
| Chr9:2094378 | 1 | 0 | 1 | 0 | Perfect(1) |
| Chr9:21458344 | 0 | 0 | 0 | 0 | NA |
| Chr9:21744263 | 0 | 0 | 0 | 0 | NA |
| Chr9:21859387 | 1 | 1 | 1 | 1 | D_7(1) |
| Chr9:21884649 | 0 | 0 | 0 | 0 | NA |
| Chr9:22186805 | 0 | 0 | 0 | 0 | NA |
| Chr9:22706823 | 1 | 0 | 1 | 0 | Perfect(1) |
| Chr9:6030900 | 0 | 0 | 0 | 0 | NA |
| Chr9:643091 | 0 | 0 | 0 | 0 | NA |
| Chr9:694483 | 0 | 0 | 0 | 0 | NA |
| Chr9:7117395 | 3 | 2 | 6 | 3 | I_A-D_19(1);D_3(2);Perfect(3) |
| Chr9:7248751 | 1 | 0 | 2 | 0 | Perfect(2) |
| Chr9:968867 | 1 | 1 | 1 | 1 | D_3(1) |

Footprints are represented by insertions (I) or deletions (D) followed by base pairs inserted or length of base pairs deleted. Numbers in parentheses are number of RILs have the same footprint at this locus.

Supplementary Data 4. Primers used in this study

| Primers | Sequence (5'-3') |
|---|---|
| Primers for transposon display | |
| MseI+0 | GACGATGAGTCCTGAGTAA |
| MseI+T/A/G/C | GACGATGAGTCCTGAGTAAT/A/G/C |
| mPing P1 | GTAGCCGTGCAATGACACTAG |
| mPing P2 | TGACACTAGCCATTGTGACTG |
| Primers for qRT-PCR | |
| PingORF1_F | CAACAAGACCCAAATTCATGTTTG |
| PingORF1_R | ACTTGGTGGCTGTGGAGATATTGG |
| PingTPASE_F | TACGTATTGTGGACGCATTAGGC |
| PingTPASE_R | CATCAGCACCACTACCAGTAGCC |
| Actin_F | CCAGTGGTCGTACCACAGGTATTG |
| Actin_R | AGTAACCACGCTCCGTCAGG |
| Primers for validation of excisions | |
| RME1F | ATAAGCAAGCTAGTTTGGGCCT |
| RME1R | AAAGCTCTAGATTGACGGCCAA |
| RME2F | TGACACTACTGTGACAGCATCC |
| RME2R | TAACTCACTCACCATGTGGCAA |
| RME3F | TGGGAAGTGATGAGGAGGAGAT |
| RME3R | GCGCGGGGGATTAGAATACTTA |
| RME4F | GCACTCCGAAGAAGCAAAAAGT |
| RME4R | GGCCCCGCGATTAGTTACTAAT |
| RME5F | CACTACTCATCAGCAAGGTGGT |
| RME5R | GCTACATGCTACAGTGACGAGA |
| RME6F | GCCGTACGTGGTTTCAGAAAAA |
| RME6R | GGAGAAAAAGGACTAGCTGGCT |
| RME7F | ATCCGATTCTCAGCTCAGCTTC |
| RME7R | CACCATCAGCCAGTACCTTCTT |
| RME8F | TAAACACAACCTTTGTCACGGC |
| RME8R | CCGCAATCAATACCGCATTCAT |
| RME9F | CCATCCCTTGATCACCCCTAAG |
| RME9R | AACACGAAACAACAGAACACCT |
| RME10F | CAAGATAACTGCCACCAGGAGT |
| RME10R | TGAAGGGTGTTGCATGGGATAA |
| RME11F | ATGTTCCATGGGCCAGAGAATT |
| RME11R | TGTGGCCGAGTATATTGGGATG |
| RME12F | CTGTTGTACTCCTTTCGTCCCA |
| RME12R | TACCGTGCCAAGTGATGAATGA |
| RME13F | AACAAACGATGCTTTCTGCTCC |
| RME13R | ATTTTCTGTGTGCACCGTTCTG |
| RME14F | GCCTCGTCAGAAAAACAGCTTT |

| | |
|---|---|
| RME14R | TGATGAATGGAGTGCTGCTGAT |
| RME15F | TCAAACACACCTACACGCTGAT |
| RME15R | GAACTTCCTGTCTCTGCACTCA |
| RME16F | ATTCTCTTTTCGCTTTGTCGGC |
| RME16R | TTAATACATGGCCCCACCTGTC |
| RME17F | GGTGACGTGTTACCAAATACCG |
| RME17R | GCAGTCTGCGTCCATGATAAAC |
| RME18F | TGAATGAACCATCTCTCGTGCA |
| RME18R | TGAATTGGAGCCAGTAGTTGACT |
| RME19F | GGACAGGTGGAGTATTCCCTTG |
| RME19R | ATTCTTCTCTCCTGCTTGTGGG |
| RME20F | CAACTTTCCATCAGTTGCAGGA |
| RME20R | GAGGTTGGGAAGAGAGAATGGG |
| RME21F | ATTTAGGGGAGTGTGTCGTGTC |
| RME21R | TTCTTCTCCGGGTGTGATGAAG |
| RME22F | CCAATGGAAGCAGAGGAGGAAT |
| RME22R | CCACGATCGATGTCACACATTG |
| RME23F | CTTCCTCTAGATCTAGCCCCGA |
| RME23R | CATCAACGGATGCAGATGGAAC |
| RME24F | AATGAGATGAGACAGTGCAGCA |
| RME24R | TCACTGGCACACACTAGCTTAG |
| RME25F | TGGGCTCACATGCTAATCAACT |
| RME25R | CAACCCAGATCTCTTCAGCCTT |
| RME26F | ACAACCTAGGGTTTAGTGTGCT |
| RME26R | GATGACACGAGGCTAGGTGAAA |
| RME27F | AATCAAGTTCGGGAGCTTGGAT |
| RME27R | TCCATCCAGAACGGCCAATA |
| RME28F | GCCGCTGCTTTGTTTTACTCTT |
| RME28R | TTGCATCAATGATGACATGGCA |
| RME29F | GTTGTCAAGTCGTCAGTTGTCG |
| RME29R | ACGATACAATTGCCTGGACAGT |
| RME30F | GTGCGTCCAAAAGGTAGCTTTT |
| RME30R | ACGATACGAAGGAAGCAGGATC |

Primers for validation of structrual variations associated with *mPing* loci

| | |
|---|---|
| SV22_C1F | GTGGCATTTTATCCGGTGCATT |
| SV22_C1R | TGTGAGAGTGTGAGTGCTTTGT |
| SV_R26_2F | TCAGTGAACATGGAGAGGGTTG |
| SV_R26_2R | GCAGACATGGCAGTTTCACTTT |
| SV_R26R | TAGCAGACATGGCAGTTTCACT |
| SV_R22F | ATGACAAAAGCAGGCAACACTC |
| SV_R22R | TATCCTCCAATTTGCCGTCGAA |

| | |
|---|---|
| SV198_C9aF | ACCTCTGTCTCCATACCATCCA |
| SV198_C9aR | CGGGTGAGAAGGTACATCGAAA |
| SV198_C9bF | ACGTGGACCGAATAGGAAACAT |
| SV198_C9bR | AGCAACCGTGTTATAGCTGGAA |
| SV_C1_29LF | TGCTTGCAGTATACGAGGGTTT |
| SV_C1_29LR | ACTCCGGCAAGTCCTAATCAAG |
| SV_C1_29RF | GATTTTGAAAGCTGTGGTGGCT |
| SV_C1_29RR | GAACACATTATCCGAACCAGCG |
| SV_C7_10LF | TGCCACGAAATAGTTAACCCGA |
| SV_C7_10LR | CTCACCTGATGCCTGAACGATA |
| SV_C7_10RF | GACCAGTGAACAAACAGCAACT |
| SV_C7_10RR | TGGAATTGCTAGTGCTGTCTGT |
| SV_C10_13LF | CCTTGTACTTTTGGCTGGTTCG |
| SV_C10_13LR | GAGTGATTGATGCCCCGAAAAC |
| SV_C10_13RF | AGTACTTTGTACTTCCCTGTGCT |
| SV_C10_13RR | TCATGCAAGCGTGTGTTTTGAT |
| SV_C10_17LF | AACTAAGCGGAGGTAGAAGCAC |
| SV_C10_17LR | CACAAGCGAAGAAACACTCTGG |
| SV_C10_17RF | TGGCTGTTCCTCTTGTCTCTTG |
| SV_C10_17RR | TGACGGCCGATAGTTTAACGAA |
| SV_C9_22LF | AAACTGTCGCATCCGATTCTCT |
| SV_C9_22LR | TGTCCCCTTTGTCGATAACTGG |
| SV_C9_22RF | TAGGTCCAACGCTATGCTGAAG |
| SV_C9_22RR | AAAGATGAGGTCCAACGTCACA |

Primers for genotyping RILs using genome-wide markers developed using *mPing* polymorphysim between HEG4 and Nipponbare

| | |
|---|---|
| C1_29MF | AATGTCACCATGGCTCCTGTAG |
| C1_29MR | GCTTGGGCGTTGTCAAACTAAT |
| C4_26MF | CTCCTCTCATCTTGCGCCTATG |
| C4_26MR | GATTCCCGCTTATCCTCCAGTT |
| C6_27MF | ATGAATCATGGCACTGTCTCGT |
| C6_27MR | CTGGGAAAGCAGAGTAGTGGTT |
| C7_7MF | TGGCCGGAGTAGTTTTACAGAG |
| C7_7MR | CAACTTCTCTCAGGACACGGAA |
| C10_6MF | GTTGCGGATTTCCCTATCATGC |
| C10_6MR | AGGCTGAAGTTACTGCTTTGGT |
| C11_7MF | GTTGCAAGACACAATCCTCCAC |
| C11_7MR | ATGGCTTTGATCCTCCACTAGC |
| C12_3MF | AAATTCCGACACTCTCTGGCAT |
| C12_3MR | GGATGTGCTCCGAATGATGTTG |
| C12_19MF | AGAGACATAGACTTGGCCAACG |

| | |
|---|---|
| C12_19MR | GAGTACATATGACCGGGGGAAC |
| MADS50F | AAAAGTGGGTAGTGTTGGCTCT |
| MADS50R | CTGCTCTTCTTCTTGTCCCCAT |
| 37180F | GCAGGAGAGTGAACACCAGTAA |
| 37180R | AACCTTGAATCGGTCGACATGA |
| 3850bF | ACAGACAGATCGTGATGATGCA |
| 3850bR | TCAAACCGGCTGAAGATTAGCT |
